# Single-cell based elucidation of molecularly-distinct glioblastoma states and drug sensitivity

**DOI:** 10.1101/675439

**Authors:** Hongxu Ding, Danielle M. Burgenske, Wenting Zhao, Prem S. Subramaniam, Katrina K. Bakken, Lihong He, Mariano J. Alvarez, Pasquale Laise, Evan O. Paull, Eleonora F. Spinazzi, Athanassios Dovas, Tamara Marie, Pavan Upadhyayula, Filemon Dela Cruz, Daniel Diolaiti, Andrew Kung, Jeffrey N. Bruce, Peter Canoll, Peter A. Sims, Jann N. Sarkaria, Andrea Califano

## Abstract

Glioblastoma heterogeneity and plasticity remain controversial, with proposed subtypes representing the average of highly heterogeneous admixtures of independent transcriptional states. Single-cell, protein-activity-based analysis allowed full quantification of >6,000 regulatory and signaling proteins, thus providing a previously unattainable single-cell characterization level. This helped identify four novel, molecularly distinct subtypes that successfully harmonize across multiple GBM datasets, including previously published bulk and single-cell profiles and single cell profiles from seven orthotopic PDX models, representative of prior subtype diversity. GBM is thus characterized by the plastic coexistence of single cells in two mutually-exclusive developmental lineages, with additional stratification provided by their proliferative potential. Consistently, all previous subtypes could be recapitulated by single-cell mixtures drawn from newly identified states. Critically, drug sensitivity was predicted and validated as highly state-dependent, both in single-cell assays from patient-derived explants and in PDX models, suggesting that successful treatment requires combinations of multiple drugs targeting these distinct tumor states.

**Significance:** We propose a new, 4-subtype GBM classification, which harmonizes across bulk and single-cell datasets. Single-cell mixtures from these subtypes effectively recapitulate all prior classifications, suggesting that the latter are a byproduct of GBM heterogeneity. Finally, we predict single-cell level activity of three clinically-relevant drugs, and validate them in patient-derived explant.

## Introduction

Glioblastoma (GBM) represents the most common malignant brain tumor in adults and is associated with a dismal outcome (Ohgaki and Kleihues, 2005). Prognosis of GBM was shown to be associated with molecular features, including gene expression, as well as genetic and epigenetic variants (Ceccarelli et al., 2016; Freije et al., 2004; Liang et al., 2005; Murat et al., 2008; Nutt et al., 2003; Phillips et al., 2006; Verhaak et al., 2010; Wang et al., 2017). Based on these features, GBM has been clustered into distinct subtypes based on two largely non-overlapping schemas arising from bulk-tissue studies, including (Phillips et al., 2006) and (Ceccarelli et al., 2016; Verhaak et al., 2010), with further refinements in (Wang et al., 2017). Specifically, cluster analysis using outcome-associated genes identified three subtypes including proneural (PN^P^), mesenchymal (MES^P^), and proliferative (PRO^P^) (Phillips et al., 2006). In contrast, by further leveraging genetic and epigenetic information, complementary analyses identified four subtypes, including proneural (PN^W^), mesenchymal (MES^W^), classical (CL^W^), and neural (NEU^W^) (Verhaak et al., 2010). The latter was later attributed to normal tissue infiltration (Gill et al., 2014) and subsequently dropped (Wang et al., 2017). However, despite ultimately proposing the same number of subtypes, the two classification schemas are largely non-overlapping when applied to the same GBM cohorts. Specifically, while samples classified as MES^W^ (Wang) and MES^P^ (Phillips) are reasonably concordant, those classified in the other subtypes have only minimal overlap.

We propose that—once microenvironment and normal tissue infiltration are discarded—the major difference in subtype classification arises from two key contributions: (a) the plastic coexistence of single cell populations representing distinct transcriptional states within each tumor (Burrell et al., 2013) and (b) the use of *ad hoc* gene sets for classifier training purposes. Indeed, both the Phillips and the Wang classifiers rely on upfront bioinformatic filters to select small, informative genes subsets, based on either outcome (Phillips et al., 2006) or on subtraction of microenvironment contributions (Wang et al., 2017).

This issue could be effectively resolved by leveraging single-cell based classifiers, which would not be affected by tissue heterogeneity. However, while several single-cell GBM datasets have become available (Darmanis et al., 2017; Patel et al., 2014; Tirosh et al., 2016; Wang et al., 2017; Yuan et al., 2018), rather than being used to generate more universal, single-cell classifiers that could harmonize across all datasets, these studies either clustered the data for further classification using prior, bulk-tissue classifiers or focused on biologically-relevant subpopulations identified by classical developmental markers. Indeed, we argue that single cell RNASeq profiles lack sufficient depth to allow training high-quality single cell classifiers, while bulk-tissue subtypes are largely ineffective for single-cell classification, resulting in high rate of ambiguously or non-statistically significantly classified cells. This suggests that single-cell GBM subtypes capable of generalizing to multiple datasets are still elusive and that bulk-tissue classifiers are poorly suited to single-cell analysis. To address this issue, we propose a novel, fully unsupervised clustering approach based on protein-activity rather than gene expression, which can be applied to either single cells or bulk tissue, producing results that are not only consistent with, but also recapitulate the activity of critical lineage marker proteins, playing critical roles in GBM biology, despite the fact that their encoding genes may be undetectable in single cell transcriptomes.

Training a single-cell classifier must address a critical “gene-dropout” issue, associated with the shallow nature of single-cell RNA-sequencing (scRNA-Seq) (Kharchenko et al., 2014). This severely limits both the number and, more importantly, the dynamics of gene expression, with the majority of genes either not detected or detected by a single mRNA read (Kolodziejczyk et al., 2015). To address this issue, we decided to rely on a fully unsupervised classification methodology, based on the metaVIPER algorithm (Alvarez et al., 2016; Ding et al., 2018a). MetaVIPER computes the activity of all proteins that mechanistically regulate the transcriptional state of the cell—including transcription factors (TFs) and co-factors (co-TFs)—based on the expression of their transcriptional targets, as inferred by ARACNe (Accurate Reconstruction of Cellular Networks algorithm) (Basso et al., 2005). This has critical consequences, as while Pearson correlation between a 30M- and a 50K-read gene expression profile (typical of single cells) is quite low (ρ < 0.3), correlation between metaVIPER-inferred protein activity at the two depths is very high (ρ > 0.85) (Alvarez et al., 2016; Ding et al., 2018a). This makes the analysis extremely robust and allows full quantitative characterization of >6,000 regulatory and signaling proteins, independent of whether their encoding gene is detectable in single-cell transcriptomes.

Several studies have shown that protein activity profiles produced by metaVIPER-—or by its precursors VIPER and MARINa—represent more robust descriptors of transcriptional cell state compared to gene expression (Alvarez et al., 2016; Aubry et al., 2015; Bisikirska et al., 2016; Brichta et al., 2015; Della Gatta et al., 2012; Ikiz et al., 2015; Kushwaha et al., 2015; Lefebvre et al., 2010; Repunte-Canonigo et al., 2015; Rodriguez-Barrueco et al., 2015; Talos et al., 2017), including in GBM (Carro et al., 2010; Chen et al., 2014; Ding et al., 2018a). Here, we show that unsupervised metaVIPER-based cluster analysis produces four distinct GBM subtypes that harmonize across both single cells and bulk tissue profiles.

The molecular subtypes identified by the analysis are markedly distinct from prior classifications and represent a novel stratification of the disease that fully integrates bulk and single-cell analyses. However, to show their consistency with legacy studies, we show that all prior subtypes are effectively recapitulated by cell mixtures drawn from the newly identified subtypes. This was accomplished using single-cell profiles from six orthotopic, fluorescently-labeled patient-derived xenograft (PDX) models established at the Mayo Clinic, which were specifically selected to represent the full gamut of prior subtypes. Prior subtypes may thus reflect tissue heterogeneity more than transcriptional identity.

Consistent with these findings, a single-cell-based classifier was able to unambiguously classify >90% of the single cells from three distinct cohorts, while prior classification schemas only achieve ∼50%. Moreover, compared to previous clustering efforts, protein-activity-based analysis significantly improved both cluster tightness—as assessed by silhouette score—and cross-cohort reproducibility. Furthermore, metaVIPER analysis of single cell could effectively quantify the activity of proteins playing a critical role in GBM, including those reported by (Carro et al., 2010) and (Wang et al., 2017), most of which were either not detectable by transcriptional analysis or had inconsistent cross-subtype expression. Finally, the analysis identified novel determinants of tumor cell state that contribute to better understanding of their biology.

Critically, the four new transcriptional states identified by metaVIPER-based cluster analysis were predicted to have differential sensitivity to therapeutic agents by OncoTreat analysis (Alvarez et al., 2018), thus providing a key rationale for the need of combination therapy to manage this disease. To experimentally validate these predictions, we focused of three clinically-relevant drugs—etoposide (a topoisomerase II inhibitor), R04929097 (a gamma-secretase inhibitor), and panobinostat (a pan-HDAC inhibitor). Analysis of single cells from patient-derived tumor explants and PDX models treated with these drugs fully confirmed their predicted subtype-specific activity.

## Results

A workflow of the approach used to harmonize tumor subtype identification across distinct bulk-tissue and single-cell GBM datasets is shown in Fig1SupFig. Specifically, individual gene expression profiles were first transformed into protein activity profiles using the metaVIPER algorithm, using ARACNe-inferred networks from five distinct GBM datasets (see Fig2SupTable and Methods). Unsupervised analysis of multiple datasets shows use of protein-activity significantly outperforms gene expression, both in terms of cluster tightness, as assessed by silhouette score (*SS*) analysis, and in terms of reproducibility across datasets.

### Characterizing inter-tumor heterogeneity by cross-cohort analysis

We first compared the subtypes proposed by *Phillips* (Phillips et al., 2006) and *Wang* (Wang et al., 2017) to subtypes produced by cluster analysis of metaVIPER-inferred protein-activity. Notably, the latter is fully unsupervised as it uses all transcription factors and co-factors, without any outcome or biology-based filtering. For consistency with the Wang and Phillips analyses we chose *K* = 3 (i.e., 3-cluster analysis). Consistent with prior studies (Alvarez et al., 2018; Califano and Alvarez, 2017), protein-activity-based clusters produced much tighter clusters (based on silhouette scores, *SS*), than gene-expression (Figure 1AB vs. DE). Specifically, following gene expression clustering, we observed *SS* ≤ 0.25 for most of Phillips and Wang samples (76% and 96%, respectively), compared to only 23% and 35% of samples clustered by protein-activity-based analysis. Typically, *SS* = 0.25 is used as a threshold for effective cluster analysis (Rousseeuw, 1987).

To further compare Phillips and Wang subtypes, we used their published classifiers to annotate the complementary cohort (i.e. Wang cohort with Phillips classifier and vice-versa). Then, for each dataset, we used Gene Set Enrichment Analysis (GSEA) (Subramanian et al., 2005) to compare the enrichment of the top 50 most differentially expressed genes in each Phillips cluster in genes differentially expressed in each Wang cluster, and vice-versa (see Methods). We chose 50 genes to both have sufficient power for GSEA analysis and to avoid bias due to different size gene-sets. Finally, we integrated p-values from complementary analyses in the two datasets (e.g., PN^P^ vs PN^W^ was integrated with PN^W^ vs. PN^P^) using the Stouffer’s test. The analysis produced a 3×3 heatmap comparing the genes identified by the two classification schemas in both cohorts (Figure 1C). As previously reported, the only subtype with statistically significant cross-cohort enrichment was the mesenchymal one (–log *p* = 4.5), suggesting poor cross-cohort reproducibility of gene expression-based clustering. This was also reflected by the equally dismal overlap of samples classified based on the Wang and Phillips subtypes across the two datasets (Figure 1GH).

**Figure 1:**
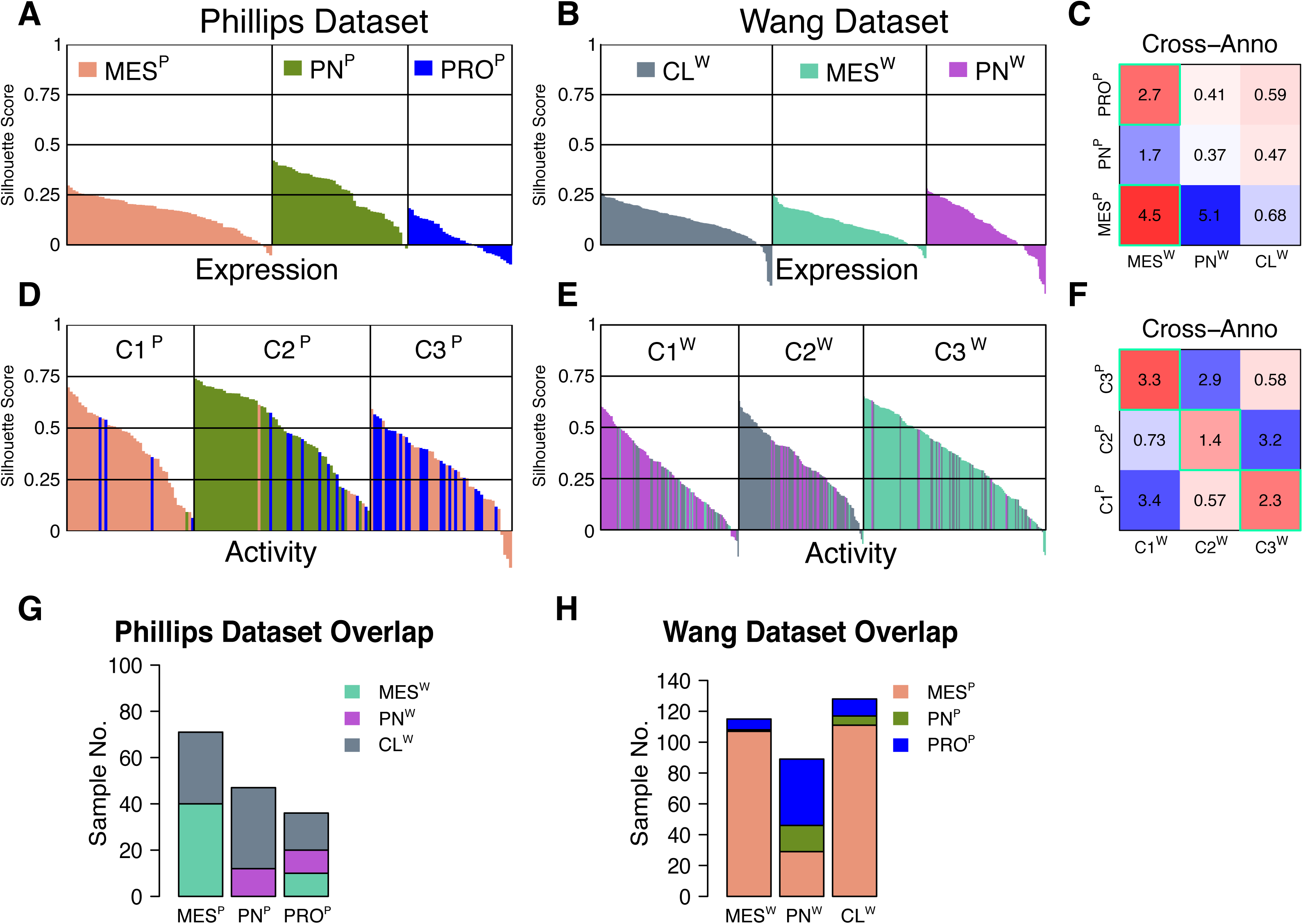
Comparison of gene expression and protein activity-based clustering in the Wang and Phillips datasets. (A,B) Reproducibility assessment of gene-expression-based cluster analyses. Analysis of *Phillips* samples (Phillips et al., 2006) classified as MES^P^ (salmon), PN^P^ (green), and PRO^P^ (blue) dataset and of *Wang* samples (Wang et al., 2017) classified as MES^W^ (teal), CL^W^ (gray), and PN^W^ (purple) revealed average silhouette scores *SS* < 0.25. (D,E) In contrast, (*K* = 3) protein-activity-based PAM cluster analysis of the same datasets generated much tighter clusters with *SS* ≫ 0.25. Samples in each protein-activity-based cluster are color-coder according to their original classification. (C,F) Cross-cohort annotation analysis based on gene expression shows poor conservation across datasets. In contrast, cross-cohort cluster conservation based on protein-activity was consistent and statistically significant. For each pairwise subtype comparison, -log *p*-values were computed by enrichment analysis of the top 50 differentially expressed genes or differentially active proteins, by GSEA. Statistically significant enrichments are shown with a cyan border. (G,H) Number of overlapping samples by cross-cohort annotation analysis shows poor overlap of Wang and Phillips classification.

In contrast, protein-activity based clusters showed significant and highly-specific overlap of differentially-active proteins by GSEA analysis. Specifically, clusters C1^W^, C2^W^, and C3^W^ (Wang dataset) matched clusters C3^P^, C2^P^, and C1^P^ (Phillips dataset), respectively (*p* < 0.05), with no additional matches, producing a one-to-one correspondence (Figure 1F). Based on protein-activity, C1^P^ and C3^W^ were mostly comprised of samples previously classified as mesenchymal. However, C2^P^ and C2^W^ included a mix of multiple subtypes, enriched in proneural and classical samples, respectively. Similarly, C3^P^ and C1^W^ included a mix of mesenchymal/proliferative and proneural/classical samples, respectively, thus making the new classification quite distinct from prior ones.

For the previous analysis, we assessed a *K* = 3 cluster solution for compatibility with prior classifications. However, unbiased cluster optimality analysis (Rousseeuw, 1987) identified a *K* = 2 cluster solution as the best choice for both gene-expression and protein-activity-based cluster analysis (Fig2SupFig A-D). Indeed, protein-activity-based, 2-cluster analysis produced much more significant subtype separation in both datasets (Fig2SupFig EF), with high average silhouette scores (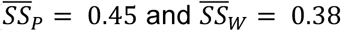, for the Wang and Phillips data respectively), suggesting the existence of only two major transcriptional states in GBM. In sharp contrast, 2-cluster gene expression analysis produced poor results, with low silhouette score and ineffective biological classification. Specifically, the Phillips cohort yielded a 102-sample cluster, mostly classified as MES^P^ and PRO^P^ and a 52-sample cluster mostly classified as PN^P^ (Fig2SupFig G), both with poor average silhouette score (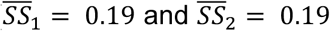, respectively). The Wang cohort produced a large 300-sample cluster, containing 90% of the samples, and a small 32-sample cluster, both of which comprised a mixture of all three Wang classes, essentially failing to stratify the data in any biologically meaningful way (Fig2SupFig H).

To confirm the high cross-cohort consistency of protein-activity-based subtypes at the molecular level, we trained a random-forest classifier on either the Phillips or the Wang subtypes and tested it on the independently inferred subtypes in the other cohort, achieving almost perfect cross-cohort classification, (AUC_PW_ = 0.966 and AUC_WP_ = 0.969, by based on Area Under the Curve analysis), respectively (Fig2SupFig IJ). This confirms that a 2-cluster, protein-activity-based classification provides more robust and universal stratification compared to prior subtypes. Consistently, in terms of specific proteins, independent analysis of each dataset confirmed high cross-cohort overlap of most differentially active proteins (*p*_PW_ = 10^-20^ and *p*_WP_ = 10^-13^, by GSEA), including established drivers of proneural and mesenchymal state (Fig2SupFig KL). For instance, *CEBPB*, *FOSL2*, *RUNX1*, and several *STAT* proteins— previously validated as determinant of the mesenchymal subtype (Carro et al., 2010)—were most activated in one cluster, while *OLIG2*, *MYCN*, and *NEUROD2*—established markers of the proneural subtype—were most activated in the other. This suggests that the principal stratification across multiple GBM cohorts occurs along a proneural to mesenchymal axis. In contrast to prior protein-activity-based reports (Carro et al., 2010) the analysis presented here is fully unsupervised and does not rely on prior, gene-expression-based subtypes.

### Iterative clustering identifies a finer-grain 4-cluster solution

we then used the iterClust method (Ding et al., 2018b) to assess whether the initial 2-cluster solution could be further stratified, resulting in a 4-cluster solution, with consistently high average silhouette scores in the range 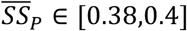 and 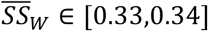 (Fig2SupFig MN). In terms of previously defined subtypes, clusters inferred from the Wang dataset were mixed; in contrast, clusters inferred from the Phillips dataset were clearly delineated. Specifically, clusters P1 and P3 were comprised almost exclusively of mesenchymal and proneural samples, respectively, while P2 included a mixture of mesenchymal and proliferative samples, and P4 a mixture of proneural and proliferative samples (Fig2SupFig M). This suggests the existence of four subtypes including quiescent proneural (PN), quiescent mesenchymal (MES), proliferative proneural (PPN), and proliferative mesenchymal (PMES) states, which are not effectively recapitulated by previous gene-expression-based subtypes.

### Single-cell classification reveals GBM intra-tumor heterogeneity and recapitulates inter-tumor heterogeneity

Despite recent availability of single-cell dataset from GBM patients (Darmanis et al., 2017; Levitin et al., 2019; Patel et al., 2014; Tirosh et al., 2016; Wang et al., 2017; Yuan et al., 2018), there have been no attempts to identify GBM subtypes de novo at the single-cell level to address the limitations imposed by bulk-tissue heterogeneity. Rather, single-cell cluster analysis was followed either by classification with prior bulk-level classifiers (Patel et al., 2014; Wang et al., 2017) or by cluster characterization with classical lineage markers (Tirosh et al., 2016; Yuan et al., 2018) or by focusing on differences between infiltrating and non-infiltrating cells (Darmanis et al., 2017). Indeed, scRNA-Seq profiles have very low depth— generally in the 20K to 100K-read range—and are thus poorly suited to generating robust classifiers, since many genes are either undetected or detected based on a single mRNA molecule.

Prior publications and previous section’s analyses show that metaVIPER-inferred TF and co-TF activity represents a more robust and reproducible descriptor of the cell transcriptional state, compared to gene expression (Alvarez et al., 2016; Califano and Alvarez, 2017; Ding et al., 2018a). This is especially important in single-cell analyses, where metaVIPER can accurately measure the activity of >6,000 regulatory proteins from the expression of their transcriptional targets—from as little as 20K-reads—independent of expression of their encoding genes (Ding et al., 2018a).

To test the existence of molecularly-distinct single-cell states, we used metaVIPER to generate protein activity profiles from published gene expression datasets representative of all three Wang subtypes (Patel et al., 2014; Wang et al., 2017) (see Methods), using ARACNe networks generated from five different GBM datasets (Fig2SupTable). Consistent with bulk-tissue analysis, IterClust (Ding et al., 2018b) identified an optimal 4-cluster solution with 3 clusters inferred in the first iteration (C1-C3) and C2 further split into C21 and C22 in the second iteration. Each cluster presented highly significant average silhouette scores (Figure 2A-C and Fig3SupTable-1). Supporting the approach’s robustness, using *K* = 2 in the first iteration (similar to the bulk sample analysis) produced a virtually identical 4-cluster IterClust solution (Fig3SupFig).

**Figure 2:**
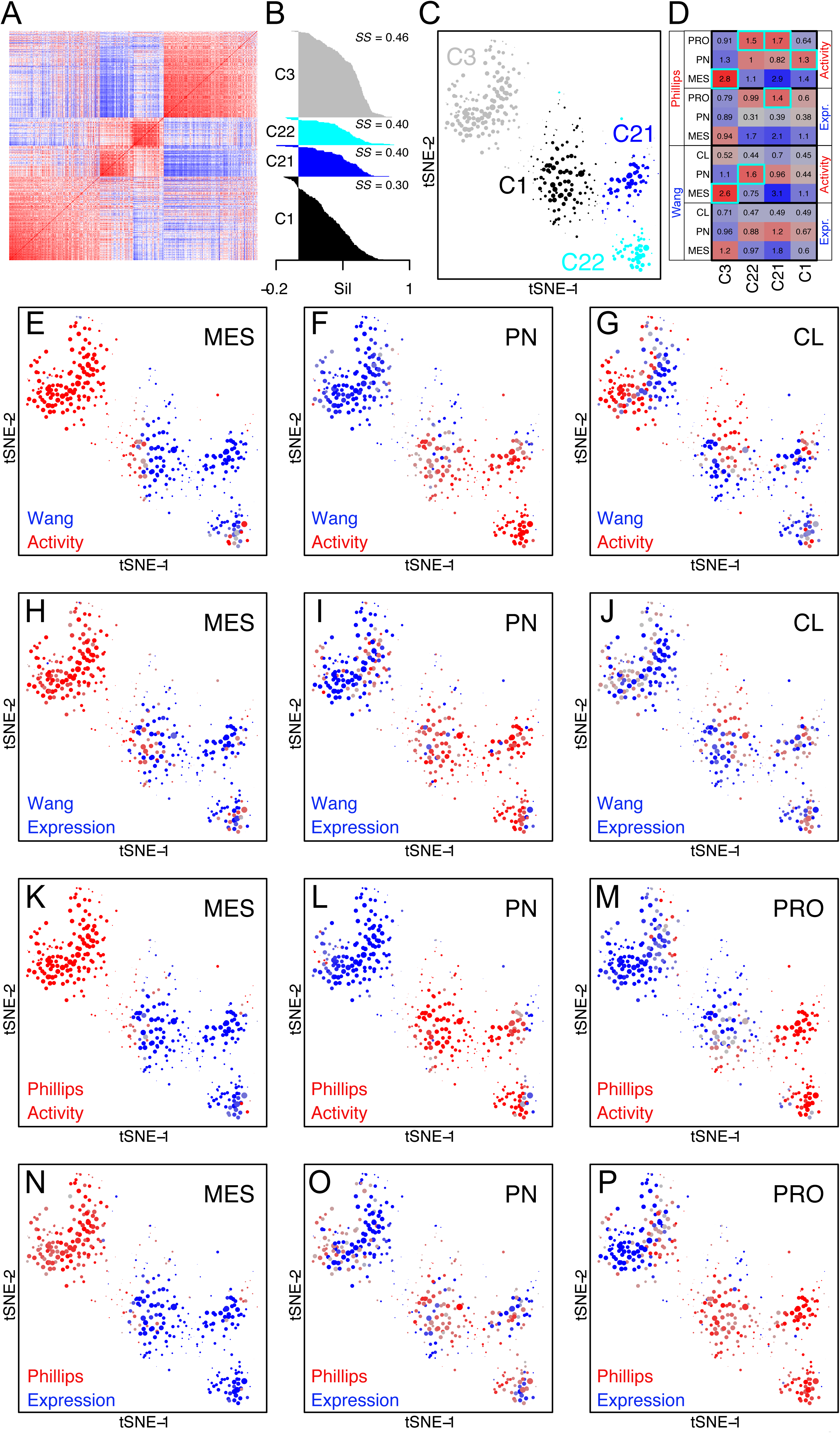
Protein-activity-based cluster analysis of the Wang and Phillips datasets. (A) Protein-activity-based, single-cell IterClust analysis (Ding et al., 2018b) identified four subtypes that are significantly distinct from previously reported Phillips and Wang subtypes at the molecular level. (B) Single-cell subtypes identified by IterClust analysis produced highly significant average Silhouette Scores, *SS* ≫ 0.25. (C) t-SNE representation of the four molecularly-distinct subtypes. Each dot corresponds to a single cell, with dot size proportional to its cluster consistency by silhouette score (D) Based on gene expression enrichment analysis, the four subtypes identified by single-cell IterClust analysis have no statistically significant overlap with previously reported subtypes, with the exception of C21 which has borderline statistically significant overlap with the Phillips Proliferative subtype (statistically significant overlaps is highlighted by thick cyan borders). Values in each cell of the heatmap represent the -log_10_(*p*) of the overlap of the 50 most differentially expressed genes in each subtype pair by Fisher Exact Test (FET). Protein-activity-based overlap analysis, also by FET analysis of the 50 most differentially active proteins, shows a higher number of statistically significant overlaps. (E-G) Individual samples colored from blue to red based on their Wang subtype classification as (E) mesenchymal, (F) proneural, and (G) classical (dot size proportional to silhouette score). Classification scores were computed based on the enrichment of the 50 most differentially active proteins in each cell in differentially active proteins in each Wang subtype, by GSEA analysis (Subramanian et al., 2005). (H-J) Identical plots using classification scores based on gene expression. (K-M) Equivalent protein-activity-based plots for Phillips subtypes, including (K) mesenchymal, (L) proneural, and (M) proliferative. (N-P) Identical plots using classification scores based on gene expression. As shown, none of the previously proposed subtypes, except for the Phillips MES subtype, effectively co-segregates with the individual clusters identified by unsupervised IterClust analysis.

**Figure 3:**
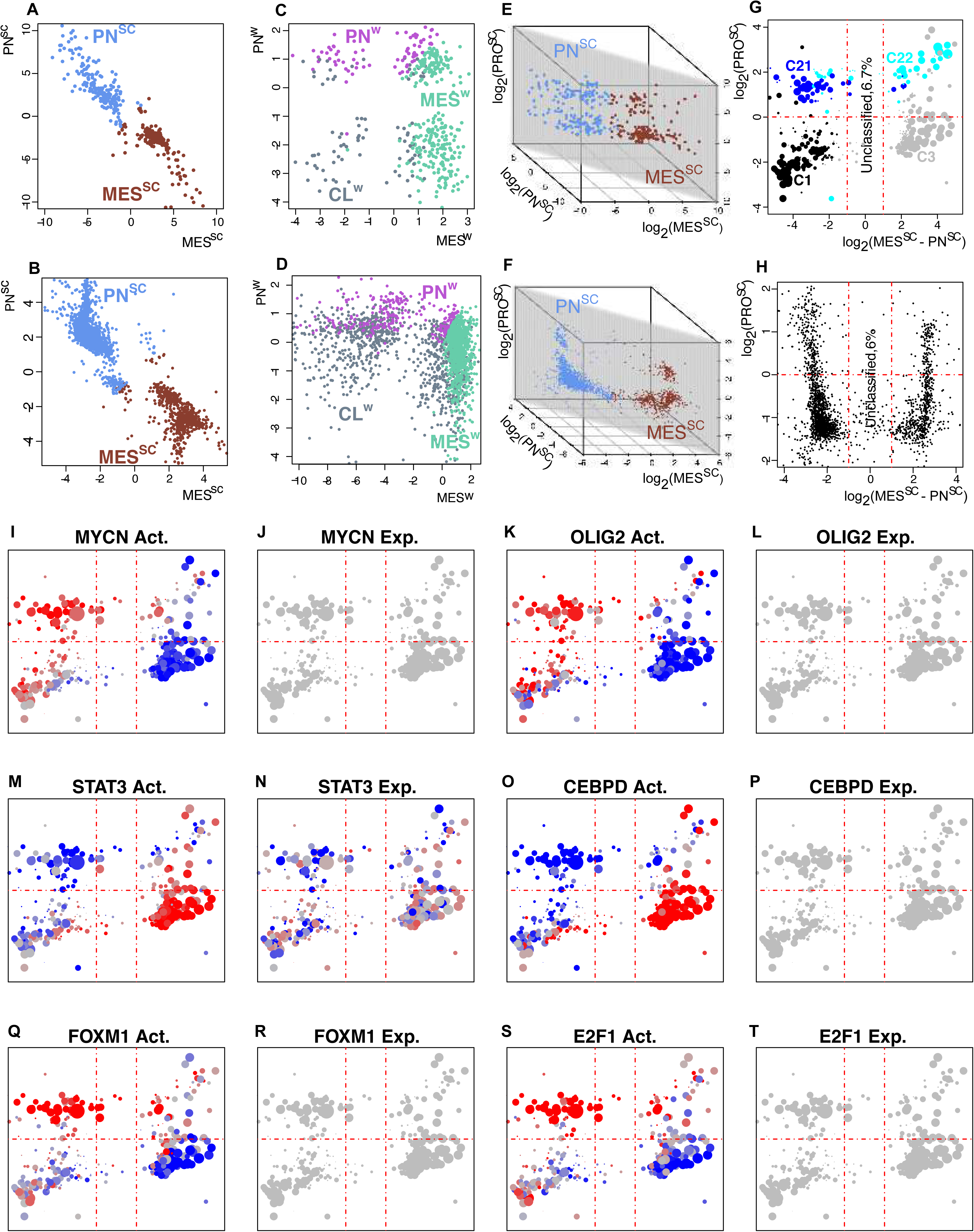
Single cell cluster analysis. Protein-activity-based, single-cell cluster analysis identified four strongly conserved cell states (*subtypes*) in the *Patel* and *Darmanis* datasets. (A,B) the MES^SC^ vs. PN^SC^ scores provide the only mutually-exclusive classification of the single cells in both datasets. (C,D) None of the previously proposed subtype-pairs could provide a mutually exclusive classification at the single cell level, including the Wang PN vs. MES (for all comparisons see Fig4SupFig 1 and 2). (E,F) An additional proliferative axis (PRO^SC^) further divides single cells into four quadrants, with only 6.7% and 6% of the cells ambiguously classified in each dataset, respectively. (G,H) The MES^SC^ vs. PN^SC^ and PRO^SC^ axes effectively separate single cells associated with each of the four subtypes identified by unsupervised cluster analysis into four quadrants, which strongly co-segregate with IterClust inferred clusters, effectively separating proneural and mesenchymal cells into quiescent or proliferative. (I, K) high activity of established proneural marker proteins identifies cells in the two left quadrants, even though they are undetectable at the gene expression level (J, L). (M, O) high activity of established mesenchymal marker proteins identifies cells in the two right quadrants, even though they are undetectable or mixed at the gene expression level (N, P). Finally, (Q, S) high activity of established proliferative marker proteins identifies cells in the top two quadrants, even though they are undetectable at the gene expression level (R, T). All dot sizes are proportional to Silhouette Scores.

We then assessed enrichment of Phillips (P) and Wang (W) subtype-specific markers in each of the four clusters identified by IterClust analysis, based both on gene expression and protein activity, as previously done in Figure 1C. Differentially active proteins were computed by metaVIPER analysis of gene expression signature associated with each cluster or subtype. Subtypes identified by single-cell cluster analysis emerged as substantially different from previously published ones (Figure 2D). Specifically, in terms of differentially expressed genes, only one cluster (C21) showed borderline significant overlap with the PRO^P^ subtype (-log *p* = 1.4). Overlap was slightly more significant at the protein activity level, with clusters C21 and C22 weakly but significantly overlapping with the PRO^P^ subtype (–log *p* = 1.7 and 1.5, respectively), C3 strongly overlapping with the MES^P^ and MES^W^ subtypes (–log *p* = 2.8 and 2.6, respectively), and C1 with borderline significant overlap with the PN^P^ subtype (–log *p* = 1.3). Overall, based on this analysis, four molecularly distinct states emerged at the single-cell level, including one quiescent/proneural state (PN*^SC^*= C1), a quiescent/mesenchymal state (MES*^SC^*= *C*3), and two distinct proliferative states (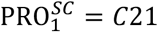 and 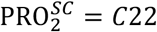). This result was consistent with the four subtypes identified by unsupervised, metaVIPER analysis of bulk-tissue profiles (Fig2SupFig M). Interestingly, the classical subtype showed no significant overlap to any of the single-cell clusters, either at gene expression or protein activity level, suggesting that it may be better suited to bulk-tissue classification due to both heterogeneity of neoplastic cells and infiltration by normal tissue. Taken together, these data suggest that, with the exception of the mesenchymal one, subtypes identified by single-cell analysis are substantially distinct from those identified from bulk-tissue analysis. For consistency, we use nomenclature taken from prior classification schemas. However, the new subtypes are quite molecularly distinct from previous ones as will be further discussed below.

Interestingly, the proteins that were most differentially active in each single-cell cluster (see Methods and Fig3SupTable-2) were consistent with those previously reported as Master Regulators of GBM subtypes (Carro et al., 2010). For instance, aberrantly activated TF/co-TF proteins in MES*^SC^* cells included experimentally validated Master Regulators of mesenchymal state, such as STAT3, C/EBPB, FOSL2, bHLH-B2 and RUNX1. Aberrantly activated TF/co-TF proteins in the PN*^SC^* and in proliferative subtypes (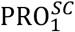 and 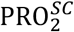) also recapitulated Master Regulators of the proneural and proliferative subtypes, respectively (Fig3SupTable-3).

### An optimal framework for single-cell-based GBM stratification

Among those identified by unsupervised single-cell cluster analysis, we then tried to identify subtypes representing mutually-exclusive transcriptional states that would optimally stratify the subtypes. If available, these would provide unambiguous classification and suggest the existence of truly distinct developmental-like states, thus providing a natural framework for single-cell GBM representation.

To accomplish this goal, we considered every possible subtype (PN*^SC^*= C1, 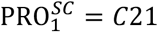, 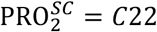, and MES*^SC^*= *C*3; Fig3SupFig) and ranked all proteins based on their differential activity in that subtype, giving us four ranked protein lists (i.e., one per subtype). For each single cell, we then computed subtype-specific classification scores for each of the four subtypes by Gene Set Enrichment Analysis (Subramanian et al., 2005), resulting in a four-score vector per cell. Finally, we plotted the entire single-cell population on the axes defined by each of six possible subtype pairs (e.g., MES*^SC^* vs. 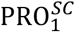) using the corresponding subtype scores, in log scale (Fig4SupFig 1 A1-A6).

Consistent with 2-cluster bulk sample analysis (Fig2SupFig L), the MES*^SC^* and PN*^SC^* related scores provided the most mutually-exclusive classification and most consistent co-segregation of the four subtypes (Figure 4A). Specifically, 93% of the cells were unambiguously classified as either mesenchymal or proneural, resulting in optimal cluster co-segregation, with 98% of PN*^SC^* cells classified as proneural, 90% of MES*^SC^* cells as mesenchymal, 86% of 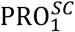 cells as proneural, and 58% of 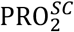 cells as mesenchymal, with remaining cells either unclassified or having mixed classification. In contrast, all other subtype-pairs produced a much greater number of ambiguously classified cells (i.e., in bottom-left and top-right quadrants) and/or poorer cluster co-segregation (Fig4SupFig-1 A1-A6).

**Figure 4:**
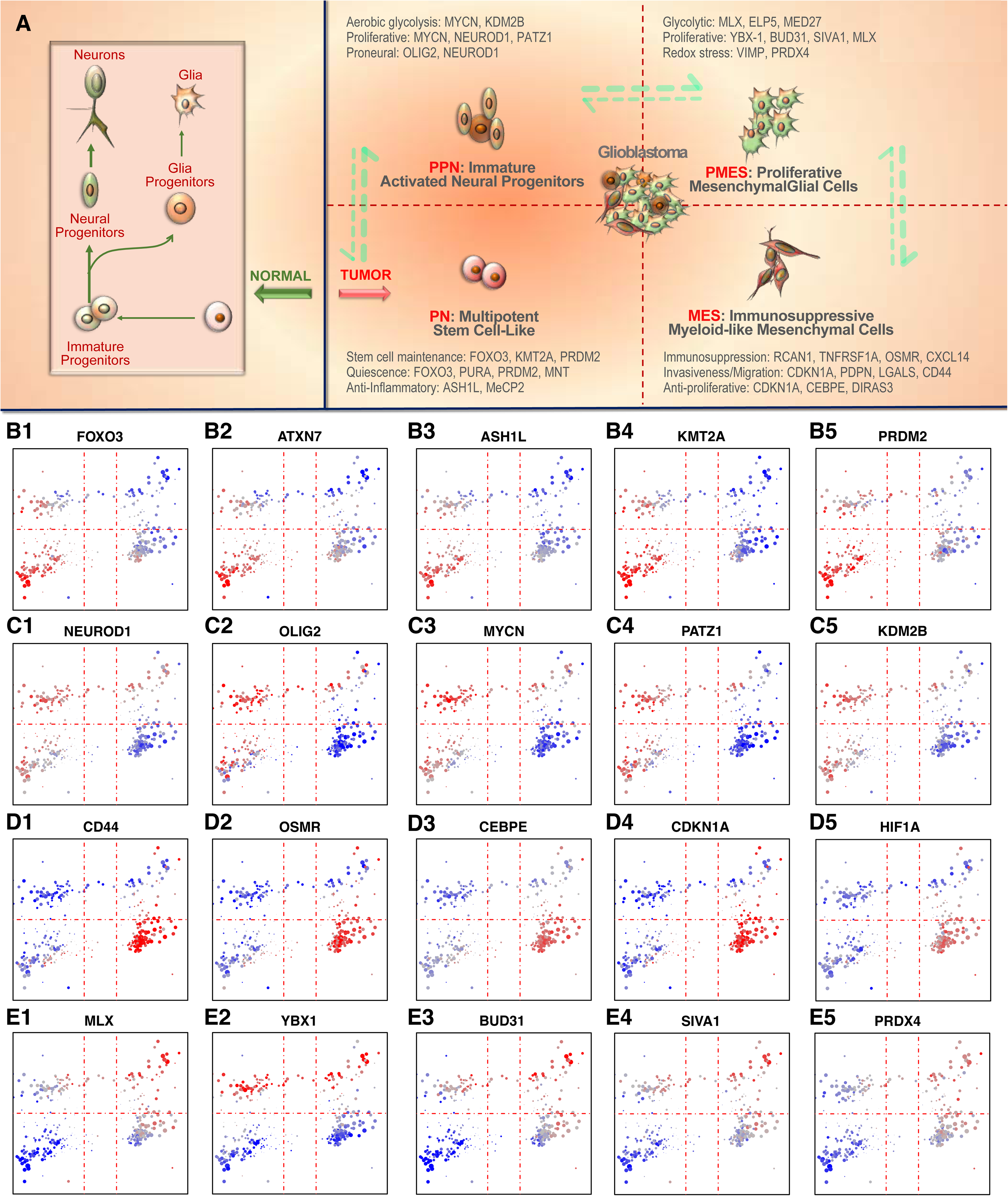
Quadrant-specific biology of individual GBM cells. (A) Each quadrant identified by unsupervised cluster analysis is associated with a distinct single-cell population that recapitulates normal lineage development of neural stem cells as well as their plastic differentiation along an aberrant mesenchymal lineage. Protein activity of key markers associated with: (B1-B5) a pluripotent, stem-like progenitor state of cells in the PN quadrant; (C1-C5) the immature, activated glial progenitor state of cells in the PPN quadrant; (D1-D5) the immunosuppressive (myeloid-like), mesenchymal, quiescent state of cells in the MES quadrant; and (E1-E5) the transient mesenchymal1 glial state of cells in the PMES quadrant. See also Fig5SupFig1-5 for additional differentially activated proteins and differentially expressed genes.

Analysis of an independent single-cell GBM dataset (Darmanis et al., 2017), using scores trained on the (Patel et al., 2014) data, confirmed the robustness of these results, with virtually identical mutually-exclusive PN*^SC^* vs. MES*^SC^* classification (Figure 4B), compared to all other subtype-pairs (Fig4SupFig-2 D1-D6). In sharp contrast, classifier scores based on Wang and Phillips subtypes failed to provide mutually-exclusive classification for any subtype-pair, for both the Patel (Figure 4C and Fig4SupFig-1 B1-B3, C1-C3) and the Darmanis datasets (Figure 4D and Fig4SupFig-2 E1-E3, F1-F3), including when using their respective MES and PN subtypes, resulting in large numbers of ambiguously classified cells.

Since the PN*^SC^* vs. MES*^SC^* axis defines an optimal classification metric for both bulk-tissue and single cells, we assessed whether a secondary axis, defined by a score determined by the enrichment analysis of proteins differentially active in 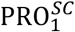 and 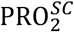 cells vs. PN*^SC^* and MES*^SC^* cells, may further stratify the four subtypes. Since these proteins were highly enriched in proliferative markers (e.g. *E2F1*, *E2F2*, *E2F3*, *FOXM1*, etc.), we defined this as proliferative, single-cell score axis (PRO*^SC^*). Consistently, analysis of a 3D scatter plot based on the PN*^SC^*, MES*^SC^*, and PRO*^SC^* score of each single cell (Figure 4EF) shows most cells laying on a diagonal plane (shaded) defined by the log_2_ (MES*^SC^* − PN*^SC^*) and by the log_2_ (PRO*^SC^*) scores, suggesting that GBM cells are optimally represented on this plane, see Figure 4GH for the Patel and Darmanis datasets, respectively.

Taken together, these data show that GBM cells coexist in plastic equilibrium between two aberrant developmental lineages (MES*^SC^* and PN*^SC^*) that are mutually exclusive, with additional stratification provided by their proliferative potential. The four quadrants identified by the S*_MES_*_/*PN*_= log_2_ (MES*^SC^* − PN*^SC^*) and *S_PRO_*= log_2_ (PRO*^SC^*) scores identify distinct single-cell populations—in both Patel and Darmanis datasets—virtually overlapping with the four clusters identified by unsupervised analysis including: quiescent-proneural (QPN), quiescent-mesenchymal (QMES), proliferative-proneural (PPN), and proliferative-mesenchymal (PMES) cells (Figure 4GH), thus providing a comprehensive cross-cohort characterization of GBM cell state in single cells (Figure 4A). Notably, while the 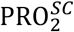 subtype is represented in both the PMES (58%) and PPN (42%) quadrants, suggesting a potential transitional nature of these cells. Remarkably, the same quadrants effectively characterize bulk-tissue profiles from the Wang and Phillips datasets, consistent with the four subtypes identified by unsupervised cluster analysis (Fig4SupFig-3). As a result, the four GBM states defined by the S*_MES_*_/*PN*_ and SPRO scores harmonize the molecular subtypes of both bulk-tissue and single cells.

### Elucidating GBM biology at the single-cell level

A critical advantage of using protein activity in single-cell analyses is that the role of key proteins that mechanistically determine cell state through their transcriptional targets can be effectively assessed on a quantitative basis. For instance, established markers of proneural subtype (e.g., *OLIG2* and *MYCN*) were undetectable at the gene expression level in the Patel dataset. However, as expected, they are activated at the protein level in the proneural quadrants (QPN and PPN) (Figure 4IJ, 4KL), proportional to proliferative potential. Consistently, established markers of mesenchymal subtype (e.g., *CEBP/D* and *STAT3*) are also undetectable at the gene expression level but are activated at the protein level in the mesenchymal quadrants (MES and QMES), inversely proportional to proliferative potential (Figure 4MN, 4OP). Finally, classical markers of proliferation (e.g. *FOXM1* and *E2F1*) are also invisible at the gene expression level but are significantly activated in proliferative quadrants (PPN and PMES) (Figure 4QR, 4ST). This dramatic increase in dynamic range allows in-depth investigation of GBM biology at the single-cell level, which would be otherwise challenging based on gene expression alone.

We further support this point by reporting on relevant GBM markers—both by at the protein activity and gene expression level—including those: (a) previously associated with or mutated in the Wang subtypes (Fig5SupFig-1) (b) most differentially active in each of the four quadrants, (i.e., QPN, PPN, QMES, and PMES) (Fig5SupFig-2) (c) previously reported as Master Regulators of GBM subtypes (Carro et al., 2010) (Fig5SupFig-3) (d) differentially activated in mesenchymal vs. proneural cells (Fig5SupFig-4) and (e) differentially activated in proliferative vs. quiescent cells (Fig5SupFig-5). In the following section, we discuss key proteins differentially active in the four quadrants that help characterize the biology of these cells. <u>Selected proteins</u> <u>(in boldface) are shown in the main text figure for easier reference (</u>Figures 5B-5E<u>), while a</u> <u>comprehensive repertoire of quadrant-specific proteins is provided in Fig5SupFig-2</u>.

**Figure 5:**
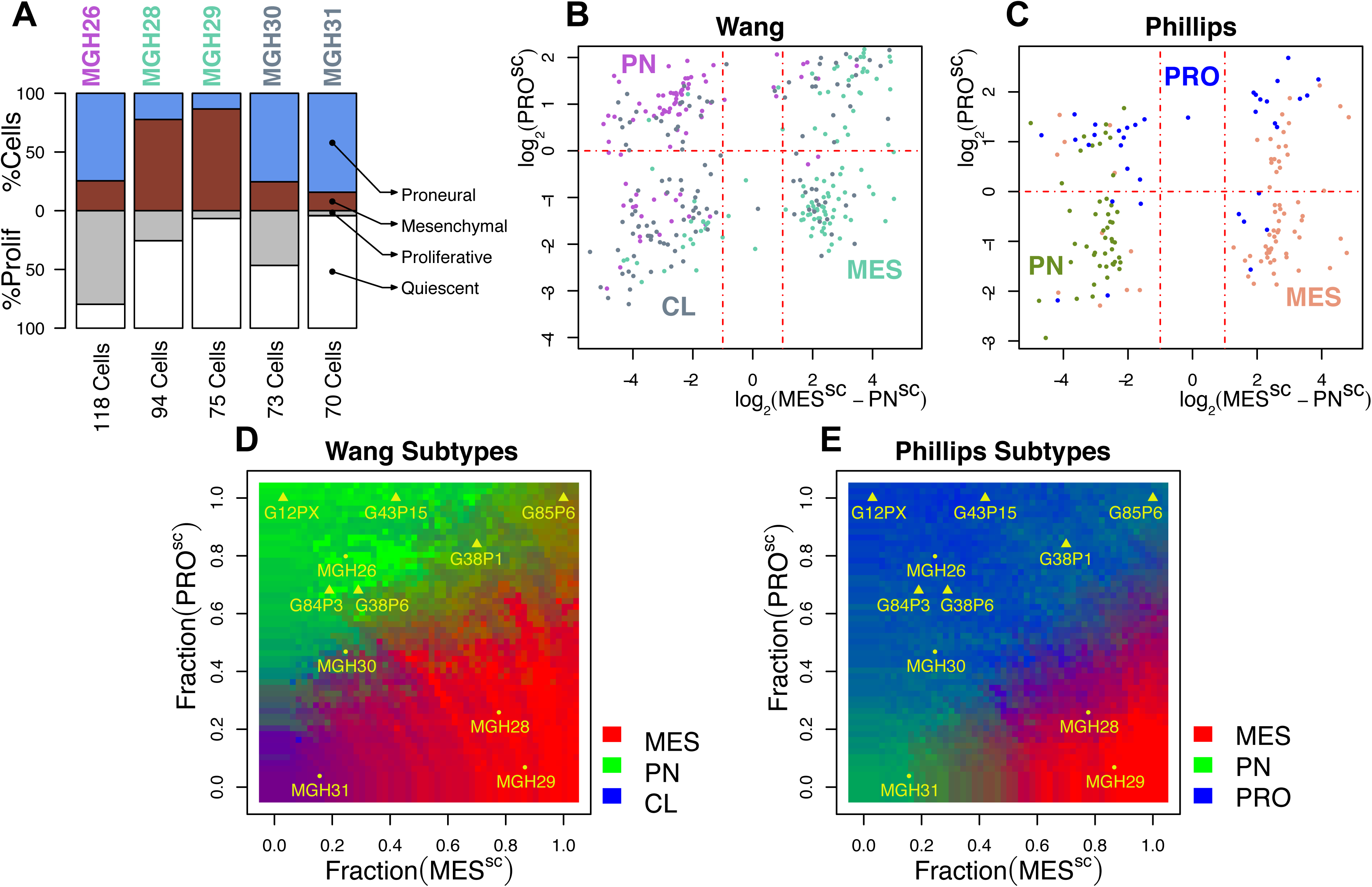
Single-cell composition of bulk samples. (A) Fraction of mesenchymal, proneural, proliferative, and quiescent single cells measured in Patel samples, including in MGH26 (PN), MGH28 (MES), MGH29 (MES), MGH30 (CL), and MGH31 (CL). (B,C) Annotation of Wang and Phillips samples using the single-cell-based classifier (see Methods). Samples were color-coded according to corresponding subtypes determined in the original studies. (D,E) Synthetic bulk samples with different fractions of proneural vs. mesenchymal (x-axis) and proliferative vs. quiescent (y-axis) single cells effectively recapitulate all prior *Wang* and *Phillips* subtypes (see Methods). Bulk-tissue samples from Patel, as well as from the PDX models profiled in this study, are projected according to their single cell composition (yellow text).

### Quiescent-Proneural subtype (PN, lower left quadrant)

cells in this quadrant are quiescent resembling neural stem cells, with activation of classic multipotent stem cell programs (Fig5SupFig-2, pp.2-5). Specifically, **FOXO3** is critical for maintaining self-renewal and for suppressing ubiquitination and proteasomal degradation of PTEN in neural stem cells (Ikeda and Toyoshima, 2017; Renault et al., 2009). Consistently, the **ATXN7/**USP22 deubiquination module of the SAGA histone acetyltransferase complex (Hu et al., 2012; Zhuang et al., 2015) and OTUD7B deubiquitinase regulate PTEN and mTORC stability, respectively (Sacco et al., 2014; Zhuang et al., 2015). Additional proteins in this compartment, contributing to maintenance of an undifferentiated cell state, include **ASH1L**, a stem cell factor that regulates *HOX* gene expression (Jones et al., 2015; Kanellopoulou et al., 2015; Miyazaki et al., 2013), MeCP2, a methyl CpG-binding repressor protein that effectively prevents transcription of glial genes (Hsieh and Zhao, 2016; Kohyama et al., 2008; Tsujimura et al., 2009), **KMT2A**/MLL1, a required factor for transcription of neuronal-specific genes (Hsieh and Zhao, 2016; Lim et al., 2009), and **PRDM2**, a cognate binding partner of RB-1 critical for maintenance of adult stem cell quiescence (Cheedipudi et al., 2015). The latter is an established central epigenetic regulator that fine-tunes the balance between stem cell quiescence and their entry into a differentiation program. It represses *MYC/MYCN* expression, which is consistently reduced in this subtype compared to the PPN^SC^ subtype. Finally, MNT is a MYC/MYCN cognate binding partners and cooperates in suppressing its activity via a feedback control loop and by binding and sequestering MAX, the canonical MYC/MYCN binding partner required to form the transcriptional activator heterodimer (Si et al., 2010).

### Proliferative-Proneural subtype (PPN, upper-left quadrant)

this quadrant is characterized by cells associated with established proneural progenitor markers, including **NEUROD1** and **OLIG2** (Figure 4C1,C2) (Christie et al., 2013; Hsieh, 2012; Hsieh and Zhao, 2016), SOX11 and SOX12 (Bergsland et al., 2011; Wegner, 2011), and EPHB3 (Baumann et al., 2013; Theus et al., 2010), suggesting that these cells behave like immature glial progenitors, preceding commitment to the glial lineage, consistent with (Yuan et al., 2018) (Fig5SupFig-2, pp.7-10). Such aberrant progenitor state is coupled with activation of established proliferative markers (e.g., **NEUROD1**, **MYCN**, **PATZ1**, E2F1, FOXM1) coupled with established regulators of aerobic glycolysis and glutaminolysis, such as **MYCN** (Dang, 2011; Wise et al., 2008) and **KDM2B** (Yu et al., 2015) (Figure 4C3,C5). **KDM2B**, in particular, is an H3-lysine 36 (H3K36) di-demethylase whose role in stem cells is to maintain status by marking sites recruiting the Polycomb Repressive Complex 1 (PRC1) to the CpG islands of developmental genes (Farcas et al., 2012; He et al., 2013). This protein plays a key role in maintaining the glioma stem cell pool and is a critical regulator of the *Ink4a/Ink4b/Arf* locus, *RB1*, and *p53* (David, 2012; Pfau et al., 2008; Tzatsos et al., 2009). Consistently, additionally highly activated proteins include the PCGF2/MEL18 sub-unit, which is required for PRC1 formation (Di Croce and Helin, 2013) and cooperates with PATZ1—an established PN marker and transcriptional activator of *MYC/MYCN* (Kobayashi et al., 2000) that also interacts with and regulates PRC1 proteins (Fedele et al., 2017)—and SCML2, a core subunit of the PRC1 complex.

### Quiescent-Mesenchymal subtype (MES, lower-right quadrant)

The protein activity profile of cells in this quadrant suggests they are quiescent and characterized by an invasive and immunosuppressive phenotype with a myeloid-like features (Fig5SupFig-2, pp. 12-15). Consistent with (Yuan et al., 2018), we find significant activation of proteins from their myeloid-like signature of single GBM cells: **CD44**, **OSMR**, CXCL14, S100A10, PDPN, PTRF, and LGALS1. Indeed, this quadrant is enriched in cells from samples similarly highlighted by them (e.g., MGH29, Fig5SupFig-6), even though virtually every sample contributes cells to this quadrant. In terms of differentially active proteins, these cells appear to phenocopy myeloid-derived suppressor cells (MDSCs), which accumulate in solid tumors and represent major immunosuppression modulators. Specifically, **CEBPE** is an established, myeloid-specific isoform and a critical factor for immature myeloid precursor differentiation (Akagi et al., 2010; Bedi et al., 2008; Rodriguez-Barrueco et al., 2015); TNFRSF1A, a prototypical TNF-alpha receptor, and TNF-alpha represent a critical checkpoint in MDSCs generation (Raveney et al., 2010; Sade-Feldman et al., 2013; Zhao et al., 2012); **CD44**, a maker of normal myelopoeisis and acute myeloid leukemia (Charrad et al., 1999; Jin et al., 2006) is expressed in mesenchymal-like GBM cells (Johansson et al., 2017; Mooney et al., 2016); CXCL14, is also an established MDSC marker (Shurin et al., 2005). Independently, aberrant activation of PDPN, LGALS1, and **CDKN1A** is consistent with the invasive phenotype presented by these cells (Ilarregui et al., 2009; Krishnan et al., 2018; Okuma et al., 2017; Shiina et al., 2016). RCAN1 is a critical regulator of the calcineurin-NFAT pathway in MDSCs, whose activity is mimicked by the classic immunosuppressive drug cyclosporine A that promotes MDSCs recruitment (Crabtree, 2001; Crabtree and Schreiber, 2009; Wang et al., 2015; Yang et al., 2018a). The cell cycle inhibitor **CDKN1A** also induces migration of classical MDSCs into the tumor microenvironment by inducing the chemokine receptor CX3R1 (Okuma et al., 2017), and likely coordinating the shift of these cells from a proliferative program to a non-proliferative, migratory one (Matus et al., 2015), possibly hypoxia-mediated, consistent with **HIF1A** activation. In this regard, the quadrant-specific protein **OSMR** is a canonical IL-6-like gp130 utilizing receptor that activates STAT3 (Chen and Benveniste, 2004), which in turn regulates ***CDKN1A*** (Bellido et al., 1998; Lundquist et al., 2003).

OSMR is also an established regulator of the mesenchymal *vs.* proneural subtype in glioma, where it regulates neural precursor activity (Natesh et al., 2015) and can heterodimerize with the EGFRvIII mutant *EGFR* receptor, a hallmark GBM mutation, which in turns partners with STAT3 (Jahani-Asl et al., 2016; Mohan et al., 2017). **OSMR**-STAT3 activation also regulates *VEGF/VEGFR* expression (Repovic et al., 2003; Tawara et al., 2019), while WWTR1 (TAZ), a Hippo pathway effector, is in turn activated by VEGF/VEGFR (Elaimy and Mercurio, 2018), suggesting a positive **OSMR**-STAT3-WWTR1 angiogenic feedback loop. Finally, the classical glioblastoma mesenchymal markers CEBP/B, STAT3, FOSL1, RUNX1, and BHLHE40 reported in (Carro et al., 2010) are all highly and specifically activated in this quadrant (Fig5SupFig-3).

### Proliferative-Mesenchymal subtype (PMES, upper-right quadrant)

finally, this quadrant appears to represent cells undergoing reprogramming to a more proliferative state via an aerobic to hypoxic glycolytic shift in metabolism but not yet overtly cycling, as shown by dramatically lower activity of proliferative markers such as FOXM1 and E2F1 compared to the PPN quadrant (Figure 4Q-T) (Fig5SupFig-2, pp.17-20). Incidentally, FOXM1 is a classic regulator of aerobic glycolysis (“Warburg effect”) (Cui et al., 2014; Shang et al., 2017; Wang et al., 2016; Zheng et al., 2016). **MLX** and **YBX-1** are known glycolytic shift regulators—controlling retrograde mitochondrial signaling to the nucleus, expression of glycolytic and lipogenic genes, and integrating *PI3K/AKT* signaling to glycolysis (Amal et al., 2015; Billin and Ayer, 2006; Butow and Avadhani, 2004; Diolaiti et al., 2015; Lasham et al., 2013; Sans et al., 2006; Stoltzman et al., 2008; Suresh et al., 2018; Xu et al., 2017). MED27, MRPL12 and ELP5 further contribute to these activities (Alexander et al., 2013; Catarina et al., 2014; Close et al., 2012; Fisher, 2018; Frei et al., 2005; Hawer et al., 2018; Rapino and Close, 2018). The transition to a more proliferative phenotype is highlighted by the activation of **BUD31, YBX-1,** SARNP, TARBP2, **SIVA1**, and PHB. The first three allow rapid mRNA splicing, processing, and transport for increased translation of proteins under proliferative stress (Climente-Gonzalez et al., 2017; Dufu et al., 2010; Eliseeva et al., 2011; Hsu et al., 2015; Lyabin et al., 2014; Marcelino Meliso et al., 2017; Saha et al., 2012; Wu et al., 2015). TARBP2 and **YBX-1** are central regulators of *DICER/Ago2* mediated miRNAs processing in GBM (Frohn et al., 2012; Wu et al., 2015), consistent with the particular heightened role of epigenetic mechanisms, including those mediated by miRNAs and non-coding RNAs, play in GBM plasticity (Balci et al., 2016; Li et al., 2016; Peng et al., 2018; Zhang et al., 2015). **SIVA1** is an E3 ubiquitin ligase that promotes degradation of the tumor suppressor and cell cycle checkpoint protein ARF, which in turn downregulates *p53* (Vachtenheim et al., 2018; Van Nostrand et al., 2015; Wang et al., 2013). Loss of *p53* and *ARF* are classical features of glioblastoma (Kim et al., 2012; Mao et al., 2012; Zheng et al., 2008). **PHB** is required for correct localization of Raf-1 in response to Ras activation and couples EGFR signaling to Ras-Raf-MEK-ERK activation, as well as for mitochondrial functions (Rajalingam and Rudel, 2005; Wei et al., 2017; Yang et al., 2018b). Activation of oxidative and ER stress programs are captured by aberrant activation of proteins such as **PRDX4** (Bulleid and Ellgaard, 2011; Zito, 2013). Interestingly, several of the MES and PPN markers, such as OLIG2, MYCN, FOXM1, E2F2, and CD44, for instance, have mixed differential activity in these cells, while proteins activated in this quadrant, such **YBX-1** or **MLX**, are also activated in subsets of PPN or MES cells. Taken together, and considering also their lower abundance, this is consistent with PMES cells representing a transitional state of cells undergoing reprogramming between the PPN and QMES states, with some possible enrichment of a distinct population of stem-like mesenchymal cells (Chandran et al., 2015), activated by glycolytic metabolism (Mao et al., 2013), which may explain the ability of migratory mesenchymal cells to reseed GBM tumors at distal sites, including repopulating the aberrant glial lineage.

Based on these findings, we propose to change the nomenclature of GBM subtypes as follows: the quiescent proneural quadrant represents a Neural *Stem-like Precursor Cell* (NSPC) subtype, the proliferative proneural quadrant represents a *Neural Progenitor* Cell (NPC) subtype. The Quiescent Mesenchymal quadrant represents an *Immunosuppressive, Invasive Mesenchymal Cell* (IIMC) subtype. Finally, proliferative mesenchymal cells define a transitional Mesenchymal-to-Proneural Cell (M2PC) subtype.

### Relating intra- and inter-tumor heterogeneity

we further assessed the single-cell composition of the 5 Patel samples, which had been deliberately selected to represent all three Wang subtypes, including PN^W^ (MGH26), MES^W^ (MGH28/29) and CL^W^ (MGH30/31). Figure 5A shows the proportion of mesenchymal vs. proneural cells, as well as the proportion of proliferative vs. quiescent cells, based on the single cell classifier scores. Samples classified as MES^W^ at the bulk level (MGH28/29) are dominated by mesenchymal cells (∼80%), while samples classified as PN^W^ and CL^W^ (MGH26 and MGH30/31, respectively) are dominated by proneural cells (∼70%). In addition, PN^W^ samples (MGH26) have a significantly higher proportion of proliferative cells (∼80%) compared to both CL^W^ and MES^W^ samples (∼30%).

We projected samples from the Wang and Phillips cohorts, color-coded according to the corresponding subtype classification, on the two principal axes identified by single-cell cluster analysis. The hypothesis, similar to what has been proposed by computational deconvolution methods (Maurer et al., 2019), is that such an analysis could identify the fractional contribution of a tissue in terms of single-cell representing distinct transcriptional states. Consistent with Figure 5A, MES^P^ and MES^W^ samples were classified as representing a predominantly quiescent mesenchymal state; PN^P^ and PN^W^ samples as representing either a quiescent or a proliferative proneural state, respectively; CL^W^ samples as representing a quiescent proneural state; finally, PRO^P^ samples as representing proliferative proneural and mesenchymal states (Figure 5BC).

Finally, we assessed whether single cell populations comprising different fractions of mesenchymal vs proneural and quiescent vs. proliferative cells could recapitulate the Wang (Figure 5D) and Phillips (Figure 5E) subtypes at the bulk tissue level (see Methods). Specifically, we created synthetic bulk samples with a variable fraction of mesenchymal vs. proneural and proliferative vs. quiescent cells, and classified them along the published Wang and Phillips subtypes. These heatmaps show that different combination of single cells distributed along the two axes could produce bulk samples classified according to all 6 subtypes.

From these analyses the CL^W^ and PN^W^ subtypes appear to represent proneural cells in either a quiescent or proliferative state, respectively, while the PRO^P^ subtype represents both proneural and mesenchymal cells in a proliferative state, thus resolving a standing controversy and providing cross-cohort harmonization.

### Passage-dependent Drift in Patient-derived Mouse Xenograft Models

To further confirm these findings in a high-quality (Fig7SupFig-1), independent dataset, we generated and analyzed scRNA-Seq profiles from six orthotopic GBM transplants (PDX models), including an early (P1) and late (P6) passage of the same PDX model (Figure 6). Human cells were fluorescently labeled for FACS-based isolation and models were selected to represent all four of the original Verhaak subtypes, based on bulk-tissue classification, including neural (G12, G22, G84), proneural (G85), mesenchymal (G43) and classical (G38, both at P1 and P6) (Mayo dataset, see Methods and Fig7SupTable).

**Figure 6:**
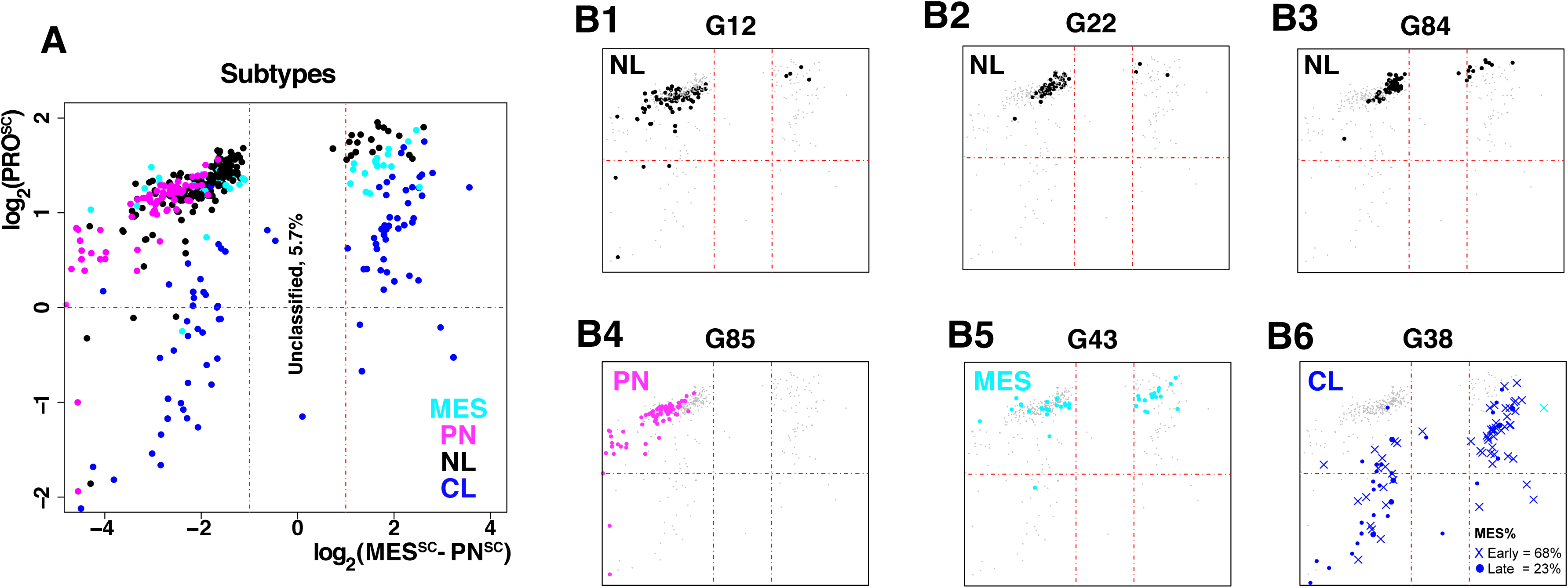
Single cell composition of orthotopic, patient-derived xenograft models. (A) Single cells are color-coded according to the corresponding PDX model subtypes, which were determined using classifier described in (Verhaak et al., 2010). (B1-B6) Single cells from each individual PDX model. Single cells from an early (P1) and late (P6) passages of the G38 model are shown together (B6).

Single cells were dissociated from these brain lesions and then scRNA-Seq profiled on the Fluidigm C1 (see Methods). Profiles were then characterized using the single-cell classifier scores (Figure 6A, B1-B6), as well as the Phillips and Wang classifier scores (Fig7SupFig-2).

Consistent with previous dataset, only the single-cell classifier produced mutually exclusive classification, along the proneural vs. mesenchymal axis. The analyses further confirmed that virtually every GBM tumor—regardless of its bulk-level classification—is comprised of cells representing all four states identified by single-cell analysis, suggesting that, despite significant bias, PDX models may effectively recapitulate the heterogeneity of the original tumor along these two axes. However, especially in later passages, PDX tumors present a significantly reduced mesenchymal vs. proneural cell fraction and a larger proliferative vs. quiescent cell fraction (Figure 6A, B1-B6).

To quantitatively assess these differences, we profiled both an early (P1) passage and a late (P6) passage of the G38 model (Figure 6B-6), originally classified as classical based on bulk-tissue analysis. Our analysis showed a highly significant decrease of mesenchymal cells between the P1 and P6 tumors (*p* = 1.5E-4, by Fisher Exact Test, Fig7). In addition, our analysis showed an even more significant decrease of mesenchymal cells in PDX models compared to primary tumors from the Patel et al. dataset (*p* = 7.1E-15). Similarly, compared to primary tumors from the Patel et al. dataset, PDX cells were significantly enriched in proliferative cells. Specifically, in primary tumors, (Figure 4G) ∼60% of the cells were classified as non-proliferative, while 97% of the PDX-derived cells were proliferative (Figure 6A) (*p* = 6.5E-69). Taken together, these findings suggest significant molecular subtype drift following serial PDX passaging.

### Multiple drugs are required to target cells in distinct quadrants

The presence of four molecularly-distinct quadrants within each tumor suggests that a single drug approach may not be effective in GBM, as each compartment may present differential drug sensitivity. To quantify potential differences, we used the recently published OncoTreat methodology (Alvarez et al., 2018) to predict sensitivity to three clinically-relevant GBM drugs at the single-cell level in the Patel et al. dataset. OncoTreat assesses the ability of a drug to revert the coordinated activity of the top 50 most statistically significant master regulators (MRs) of tumor state based on perturbational profiles of drugs in cell lines whose MR profile is significantly similar to that of the target tissue (p < 1E-5). In this case, we used perturbational profile data from two cell lines, including U87 (mesenchymal) and HF2597 (proneural) to assess a compound’s ability to modulate the activity of master regulator proteins (see Methods). We generated perturbational profiles for three drugs, including the topoisomerase-II inhibitor etoposide, the gamma-secretase inhibitor RO4929097, and the pan-HDAC inhibitor panobinostat (see Methods), all previously used in GBM clinical trials.

Consistent with our expectation, OncoTreat analysis of Patel et al. cells revealed that individual drugs are predicted to have highly quadrant-specific sensitivity. For instance, sensitivity to etoposide was predicted to be highest in proliferative proneural cells (Figure 7A), confirming that etoposide is an effective treatment for proneural GBM mouse models (Sonabend et al., 2014); in contrast, sensitivity to RO4929097 was higher in mesenchymal vs. proneural cells (Figure 7B). Finally, panobinostat sensitivity was highest in proliferative cells, independent of their mesenchymal vs. proneural status (Figure 7C). Mesenchymal-specific sensitivity to RO4929097 may result from activation of gamma-secretase-dependent signals in the mesenchymal quadrants, including *NOTCH3* (Fig5SupFig-1, pp.3) and CD44 (Figure 4D1).

**Figure 7:**
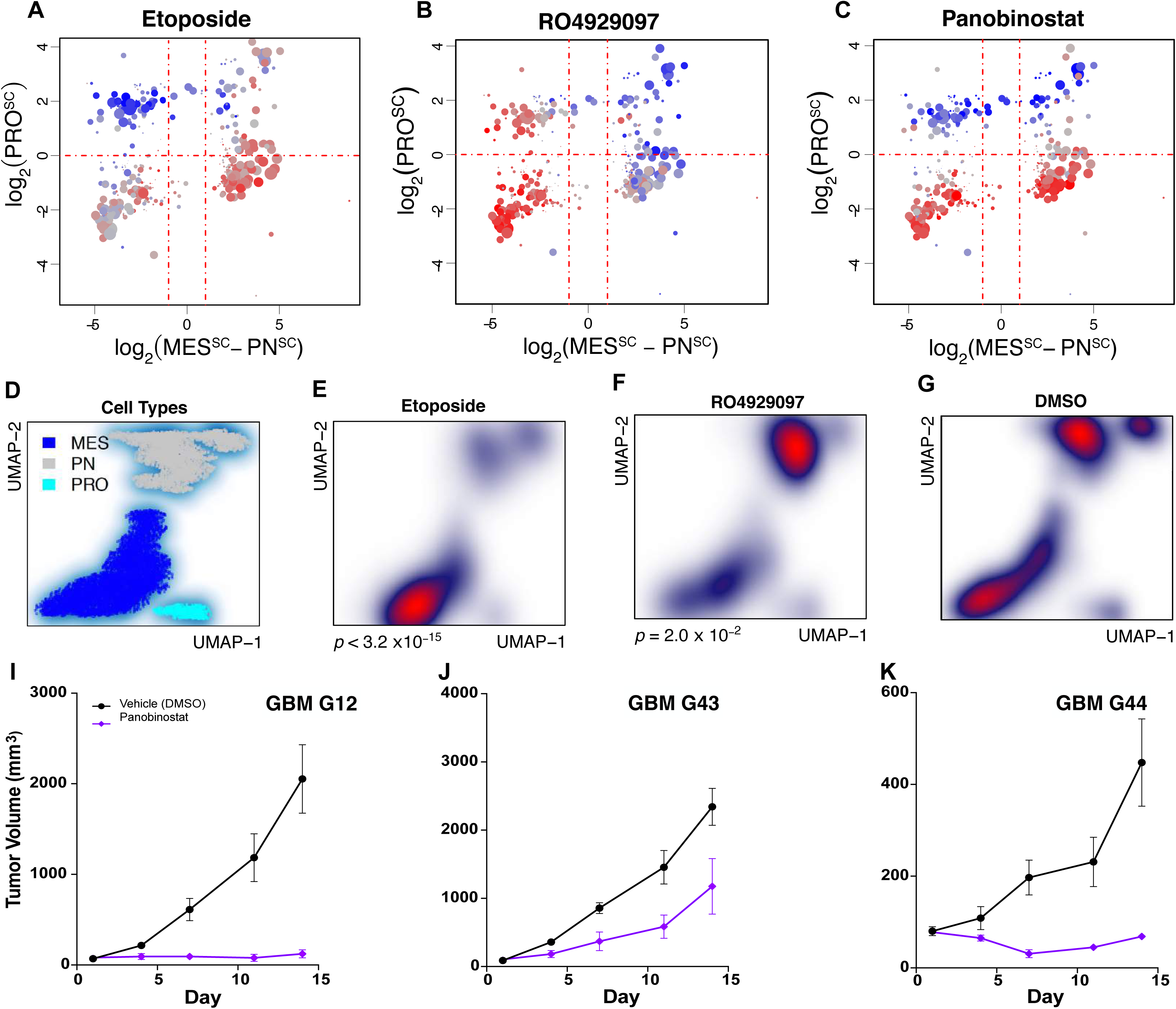
Quadrant specific drug-sensitivity prediction and experimental validation. (A-C) Subtype-specific prediction of sensitivity to etoposide, RO4929097, and panobinostat by OncoTreat analysis of single cells from the Patel dataset (see Methods). Blue dots indicate predicted inversion of single-cell-specific Master Regulator activity and thus higher drug sensitivity. (D) Single neoplastic cells, from patient derived explants, annotated according to their mesenchymal, proneural, or proliferative state (see Methods and Fig8SupFig). Single cell density following treatment with (E) etoposide, inducing significant loss of proneural cells, (F) RO4929097, inducing significant loss of mesenchymal cells, and (G) DMSO as vehicle control (see Methods). (I-K) Tumor volume curves of proliferative PDX models G12, G43 and G44, following treatment with panobinostat or 5% dextrose water, as vehicle, confirm its preferential activity on proliferative cells (see Methods).

To experimentally validate these predictions, we performed scRNA-Seq profiles of single cells from patient-derived explants. Acute slices generated from a surgical resection specimen were placed in culture conditions for 6h, and then treated for 18h with each drug, with DMSO as negative control. Non-tumor cells were removed by expression based CNV analysis (Patel et al., 2014), which effectively identified tumor cells presenting highly significant Chr7 amplification and Chr10 deletion. Cells were then further classified into mesenchymal, proneural and proliferative using the MES^SC^, PN^SC^, and PRO^SC^ single cell classifier scores (Fig8SupFig A-F). Most of the cells in these tumors are non-proliferative, with a 57/43 representation of mesenchymal vs. proneural cells (Figure 7D).

Consistent with OncoTreat predictions, etoposide and RO4929097 induced significant selection of mesenchymal and proneural cells, respectively (Figure 7E-G) (*p*_Eto_ = 3.2E-15, *p*_RO4_ = 2.0E-2). Since only a minority of cells were proliferative (2%, Figure 7D) we evaluated the effect of panobinostat in *in vivo*, in subcutaneous transplants from orthotopically established PDX models (G12, G43, and G44, established at the Mayo Clinic), which were shown to be highly enriched for proliferative cells. These models were treated with panobinostat 10 mg/kg intraperitoneal daily, 5 days on/2 days off, for 10 doses) and 5% dextrose water, as vehicle. All three models showed significant reduction in tumor growth at 15 days, compared to DMSO, including stable disease for model G12 and G44 and ∼50% tumor volume reduction for G43 (*p* = 0.034, 0.03, 0.04, respectively, by AUC analysis, see Methods).

## Discussion

Past bulk-tissue based studies have highlighted the complexity and heterogeneity of glioblastoma, in particular by suggesting the plastic coexistence of an aberrant glial developmental lineage with a less structured aberrant mesenchymal lineage and creating the premise for further studies at the single cell level (Carro et al., 2010; Ceccarelli et al., 2016; Phillips et al., 2006; Verhaak et al., 2010; Wang et al., 2017). Their strength has been the ability to monitor the expression of virtually all expressed genes first via gene expression microarrays and then RNA sequencing. At the single cell level, we have just started to look further into the complex cascade of aberrant and inter-related lineages that comprise a GBM mass (Darmanis et al., 2017; Patel et al., 2014; Tirosh et al., 2016; Wang et al., 2017; Yuan et al., 2018). However, given the low-depth of sequencing, these studies have been mostly successful in elucidating similarities between GBM cell states and their corresponding physiologic states— from neural stem cells to glial progenitor to committed glia, including reprogramming along an aberrant mesenchymal lineage—using previously established markers. Yet, with only 1,000 to 2,000 genes detected per cell, on average, most of which via a single read, these studies could not rival the depth of characterization of bulk-tissue based ones. We have thus been left at an impasse between characterizing bulk-tissue properties in detailed, genome-wide fashion and dissecting GBM’s heterogeneity by leveraging only 10% of the total number of genes.

In this manuscript we propose a novel approach that, by leveraging the network-based algorithm metaVIPER, can achieve virtually the same resolution of bulk-tissue profile analysis in single cell studies. Critically, metaVIPER does not average over cells located proximally on gene expression manifold, which is generally non-informative for a majority of under-sampled genes, see Figure 4, for instance. Rather, it uses only the low-depth gene expression profile of a single cell to infer the activity of >6,000 regulatory and signaling proteins, including transcription factors, co-factors/chromatin-remodeling enzymes, and signaling proteins. Specifically, by inferring protein activity based on the expression of its context-specific regulatory targets, metaVIPER can accurately measure the activity even of proteins whose encoding gene is not detected across the entire single-cell dataset. We show critical examples of this in Figure 4 and 5. For instance, while in high-depth datasets, such as (Yuan et al., 2018), single-cell expression of MYCN and OLIG2 can be effectively quantitated, in low-depth dataset, such as (Patel et al., 2014), their expression could not be detected in virtually any cells (Figure 4JL). Yet, activity of the encoded proteins could be effectively quantitated by metaVIPER to be highest in the proneural quadrants, proportional to the cell’s proliferative potential (Figure 4IK). This dramatic increase in gene-product dynamic range resulted in four distinct advantages.

First, the significant increase in signal-to-noise ratio allowed by conversion of gene expression profiles into protein activity profiles allowed us to generate a classification system that harmonizes across virtually all published and new datasets, including both bulk-tissue and single-cell. This is important because previous classifications were largely inconsistent when applied to distinct datasets and produced largely ambiguous on non-significant single-cell classification (Figure 4CD and s-1,2).

Second, the new classification system emerging from the analysis is for the first time organized along two orthogonal axes, one unambiguously capturing the alternative lineages identified in bulk GBM tissues (i.e., glial and mesenchymal), the other capturing the cell’s proliferative potential (Figure 4GH). Organization of both bulk-tissue samples and single cells from different datasets along these two axes provided virtually unambiguous classification of independent GBM states detected by fully-unsupervised cluster analysis. The latter is a critical advantage, since to produce reasonable classifications, all prior approaches were based on a reduced gene-set repertoire selected by *ad hoc* criteria and were not thus completely unsupervised. The newly proposed classification also provides full continuity with priori bulk-level subtypes by showing that each of them, including both Phillips and Verhaak subtypes, can be recapitulated by different mixtures GBM cells drawn from the four newly proposed subtypes.

Third, the ability to accurately quantify the activity of virtually all regulatory and signaling proteins was instrumental in providing in-depth characterization of GBM biology at the single-cell level, which is simply unattainable in the gene expression profiles. For instance, identification of specific activation of established proteins preventing expression of glial markers in the QPN quadrant suggests that these may be the cells that most critically represent a stem-like state in glioma, rather than the more proliferative PPN cells, which appear to have undergone at least partial activation of key glial programs and are most similar to glial lineage progenitors. To that respect, Figure 4 and Fig5SupFig-1,2,3,4,5 contain an inventory of all of the most differentially active proteins in each compartment, lineage, or proliferative state, including markers that could be detected from bulk-tissue but not at the single cell level. Indeed, not surprisingly, the vast majority of these proteins has activity that is clearly compartment specific, even though the expression of their encoding genes is either undetectable (e.g. OLIG2, RELB, etc.) or completely inconsistent with prior knowledge about subtype-specific expression (e.g., CD44, CDKN1A, etc.) in the Patel dataset (Fig5SupFig-1).

Finally, and most critical, the ability to study single cells with virtually the same dynamic range of bulk-tissue, supported the use of previously published methodologies, such as OncoTreat (Alvarez et al., 2018), to evaluate the sensitivity of individual tissues (in this case individual single cells) to distinct therapeutic agents. Indeed, OncoTreat predicted orthogonal, compartment-specific sensitivity of GBM cells to three clinically-relevant agents, including a topoisomerase II inhibitor (etoposide), a gamma-secretase inhibitor (R04929097), and a pan-HDAC inhibitor (panobinostat). Predictions were experimentally validated in patient-derived GBM explants (etoposide and R04929097) and in PDX models (panobinostat). An interesting corollary was that, while the overall single-cell heterogeneity of GBM is overall preserved in PDX models, these present a marked shift toward the more proneural and proliferative state, which could potentially significantly bias the effect of drugs that have marked compartment specific activity. Taken together, these data suggest that treatment of this complex and highly plastic disease will require combination of drugs targeting the individual compartments, possibly via metronomic schedules, which we plan to explore in follow-up research.

Clearly, a number of limitations remain, which should be acknowledged. In particular, while extensive experimental assays have shown that VIPER and metaVIPER can accurately measure activity of about 80% of transcription factors, co-factors and chromatin remodeling enzymes and about 70% of signaling proteins (Alvarez et al., 2016), there is still a small pool of proteins, whose activity will be mis-quantified. This requires further refinement of their set of regulatory targets (regulons), using epigenetic information (e.g. ATACSeq data), DNA-binding motifs, and perturbational data, all of which are currently being investigated. In addition, it has been established that while VIPER and metaVIPER may correctly infer the absolute value of the differential activity of most proteins, the sign may be inverted for about 30% of them, due to complex negative autoregulatory loop that, at equilibrium, may inversely couple mRNA expression and protein activity. Whether the sign of a protein activity should be inverted can be effectively assessed based on the overall correlation between its activity and the expression of its encoding gene or by relatively mundane experimental assays, including CRISPRi-mediated gene silencing followed by RNASeq profiling. Given a regulatory network and protein of interest, directionality must be assessed only once.

In summary, while the metaVIPER methodology is not perfect, it should be recognized that at this time there are no genome wide alternatives for the accurate measurement of protein activity profiles in single cells. As a result, the striking overall concordance between our metaVIPER based findings and previously established markers of GBM cell state, suggest that the analyses reported in this manuscript will provide at least some novel information that would be otherwise unattainable for the study of GBM heterogeneity, for the key molecular determinants of GBM cell state at the single cell level, and finally for the study of drug sensitivity in individual GBM cells.

## Methods

### Patient derived xenograft establishment

Studies were approved by the Mayo Clinic Institutional Animal Care and Use Committee, and all animal care procedures were followed. PDXs were established using six to seven week old female athymic nude mice (Hsd:Athymic Nude-Foxn1^nu^, Envigo, Indianapolis, IN) as previously described (Carlson et al., 2011).

### Establishing PDX models for scRNA-Seq

GFP-expressing derivatives of patient derived xenografts were established by transducing short-term explant cultures with lentivirus as previously described (Gupta et al., 2016; Gupta et al., 2014). Transduction efficiency was evaluated by examining the extent of GFP positivity by fluorescent microscopy before cell expansion in an athymic nude mouse. GFP-expressing cell lines were harvested from flank tumors of athymic nude mouse and placed on Matrigel (BD Biosciences, Billerica, MA) coated plates. Once cells were adherent, GFP positivity was confirmed by fluorescent microscopy. Cells were prepped and injected intracranially as previously described (Carlson et al., 2011). Animals were monitored daily and sacrificed by cervical dislocation at the onset of neurological decline.

### PDX model tumor dissociation

Following cervical dislocation, brains were removed and tumors carved using goggles equipped with GFP visualization (BLS Limited, Budapest, Hungary). Tumors were placed into Eppendorf tubes containing DMEM media (Corning, 10-013-CV) and immediately processed for dissociation. Tumor specimens were rinsed with PBS and trypsinized (TrypLE, ThermoFisher, Waltham, MA). The tissue was then minced into small fragments using dissection scissors and placed in a 37 degree waterbath, vortexing periodically. The solution was spun at 1200 rpm for 3 minutes, trypsin was removed, and the tissue pellet was resuspended in DMEM. The specimens were aspirated into 1 mL syringes and passed through 21G and 23G needles (Cardinal Health, Dublin, OH) until no clumps were visible. This solution was then filtered using 100 and 40 µm filters (ThermoFisher, Waltham, MA). Viability and single cell state of isolated cells was conducted by trypan blue exclusion and light microscropy. Finally, cells were spun down, washed twice in PBS, and immediately submitted to Mayo Clinic’s Core for Single Cell sorting.

### scRNA-Seq of PDX models

Cells were counted and measured for size and viability using the Vi-Cell XR Cell Viability Analyzer (Beckman-Coulter, Brea, CA). A C1 Single-Cell Array Integrated Fluidics Circuit (IFC) for mRNA-Seq (cell size 5-10 uM, Fluidigm product number 100-5759; cell size 10-17 uM, Fluidigm part number 100-5760) was primed in the C1 Single-Cell Auto Prep System (Fluidigm product number 100-7 000). While the IFC was being primed, the lysis, reverse transcription and PCR reagents were thawed and the respective chemistries were mixed in a clean room DNA-free hood. After priming, the IFC was taken to a cell culture hood and the cells were pipetted into the IFC. The IFC was placed back into the C1 System to load and separate the cells. Once the cells were sorted into up to 96 separate chambers, the IFC was removed from the C1 System and imaged on a microscope. Cell number and viability were noted on a log sheet. The lysis, reverse transcription and pre-amplification chemistries were pipetted into the IFC in the cell culture hood. The IFC was loaded into the C1 a final time to run the mRNA-Seq script overnight.

The following morning, up to 96 individual cDNA samples were harvested from the IFC. All samples were quality control tested and quantified on a 96-capillary array Fragment Analyzer (Advanced Analytical, Ankeny, IA). Smear analyses were run using the PROSize® 2.0 software (Advanced Analytical). Only samples that passed the smear analysis thresholds were selected for library construction. Each of these samples was diluted to between 200-250 pg/ul. The Illumina Nextera XT DNA Library Preparation Kit (Illumina product number FC-131-1096) was used to create individually indexed cDNA libraries for sequencing.

### Preparation and treatment of patient-derived brain tumor slices (explants) for scRNA-Seq

This work was approved by the Columbia University Medical Center Institutional Review Board before commencing the study. Acute slices were generated from a surgical resection specimen collected from an adult patient with radiographic and intraoperative histopathological characteristics consistent with primary HGG, (the final diagnosis was Glioblastoma, IDH-wildtype, WHO grades IV). The surgically excised specimen was immediately placed in a sterile 50mL canonical tube ¾ filled ice-cold high-sucrose low-sodium artificial cerebrospinal fluid (ACSF) solution containing 210 mM sucrose, 10 mM glucose, 2.5 mM KCl, 1.25 mM NaH_2_PO_4_, 0.5 mM CaCl_2_, 7 mM MgCl_2_ and 26 mM NaHCO_3_, and were kept on ice for transportation (transit time was approximately 10 minutes from operating room to laboratory). Preparation of ex vivo tissue slices was modified from methods described previously (Gahwiler et al., 1997; Kohling et al., 1999; Shimizu et al., 2011; Stoppini et al., 1991; Ting et al., 2018). Briefly, the tissue specimen was placed in ice-cold ACSF solution and 500 µm slices were generated using a tissue chopper (McIlwain TC752). The slices were immediately transferred to the ice-cold ACSF solution in 6-well plates using a sterile plastic Pasteur pipette half filled with ice-cold ACSF solution followed by a 15 minutes recovery in ACSF to reach room temperature. Slices were then placed on top of a porous membrane insert (0.4 µm, Millipore). The membrane inserts were placed into 6-well plates containing 1.5 ml maintenance medium consisted of F12/DMEM (Gibco) supplemented with N-2 Supplement (Gibco) and 1% penicillin/streptomycin (Sigma). To ensure proper diffusion into the slice, culture medium was placed under the membrane insert without bubbles. A drop of 10 µl of culture medium was added directly on top of each slice to prevent the slice surface from drying. The slices were first rested for 6 hrs with the maintenance medium in a humidified incubator at 37**℃** and 5% CO_2_. Then, the medium was replaced with pre-warmed medium containing 5 µM Etoposide (Tocris Bioscience, 100 mM stock), 50 nM RO4929097 (Selleck Chem, 10 mM stock), or corresponding volume of vehicle (DMSO). Slices were cultured with the treatment medium in a humidified incubator at 37**℃** and 5% CO_d_ for 18 hrs before being collected for dissociation.

### Dissociation of patient-derived brain tumor slices

Collected slices were dissociated using Adult brain dissociation kit (Miltenyi Biotec) on gentleMACS Octo Dissociator with Heaters (Miltenyi Biotec) according to the manufacturer’s instructions.

### Microwell-based scRNA-Seq of patient-derived brain tumor slices

Dissociated cells from each slice were subjected to microwell-based single-cell RNA-seq (Yuan and Sims, 2016) as previously described (Yuan et al., 2018) with modifications to the reverse transcription step and the sequencing method. Once the RNA-capture step was finished, sealing oil was flushed out of the devices by pipetting 1ml wash buffer supplemented with 0.04 U/µl RNase inhibitor (Thermo Fisher Scientific) and then beads were extracted from the device and resuspended in 200 µl of reverse transcription mixture. Bead-suspensions were divided into 50µl aliquots and placed into PCR tubes (Corning) followed by incubation at 25°C for 30 min and at 42°C for 90 min in a thermocycler. For sequencing, we pooled all libraries derived from the same donor, each of which was barcoded with a unique Illumina sample index. The pooled library sequenced on 1) an Illumina NextSeq 500 with an 8-base index read, a 21-base read 1 containing cell-identifying barcodes (CB) and unique molecular identifiers (UMIs), and a 63-base read 2 containing the transcript sequence, and 2) an Illumina NovaSeq 6000 with an 8-base index read, a 26-base read 1 containing CB and UMI, and a 91-base read 2 containing the transcript sequence.

### Animal studies

Patient-derived xenograft (PDX) models (GBM G12, G43, G44) for panobinostat validation studies were developed by implanting ∼10^7^ cells resuspended in Matrigel (1:1, v/v, Corning) into the subcutaneous flank of NOD (NOD.Cg-Prkdcscid Il2rgtm1Wjl/SzJ) scid gamma (NSG) mice to generate heterotopic models. Tumor growth was measured biweekly using calipers. Treatment started when flank tumors were ∼ 150 mm^3^ (tumor volume, TV = width2 × 0.5 length). Orthotopic brain tumor models were generated by stereotactic injection of ∼10^5^ cells resuspended in phosphate-buffered saline (PBS) into the right striatum. Orthotopic models were monitored biweekly or monthly by magnetic resonance imaging (MRI) to track tumor engraftment and growth. Animals were dosed with Vehicle consisting of 5% dextrose water (vehicle control for panobinostat treatments). Panobinostat (10 mg/kg IP daily) was administered on a 5-day on, 2-day off schedule for a minimum of 14 days. An area-under-the-curve (AUC) analysis was performed to compare differences in tumor volume curves between Vehicle- and drug-treated cohorts. All experiments were performed in accordance with institutional guidelines and under an approved protocol from the Memorial Sloan Kettering Cancer Center (MSKCC) Institutional Animal Care and Use Committee (Protocol 16-08-011).

### scRNA-Seq gene expression analysis for PDX models

After demultiplexing, the resulting raw reads were aligned to hg19 reference index by Bowtie2-2.2.6 (Langmead and Salzberg, 2012). Aligned reads were sorted and indexed by samtools-1.2 (Li et al., 2009). Counts matrix was measured with R package GenomicFeatures-1.24.5 and GenomicAlignments-1.8.4 (Lawrence et al., 2013) and TxDb.Hsapiens.UCSC.hg19.knownGene-3.2.2 (Carlson and Maintainer, 2015) from Bioconductor.

### scRNA-Seq gene expression analysis for patient-derived brain tumor slices

Raw data obtained from the Illumina NextSeq 500 was trimmed and aligned as described in (Yuan et al., 2018). For each read with a unique, strand specific alignment to exonic sequence, we constructed an address comprised of the CB, UMI barcode, and gene identifier. Raw data obtained from the Illumina NovaSeq 6000 was first corrected for index swapping to avoid cross-talk between sample index sequences as described in (Szabo et al., 2019). The resulting address file only contained the addresses from reads with corresponding single index. We combined the addresses from the NextSeq 500 and the corrected addresses from the NovaSeq 6000 to generate a digital gene expression matrix and filter CBs as described previously (Yuan et al., 2018). Briefly, reads with the same CB, UMI and aligned gene were collapsed and sequencing errors in the CB and UMI were corrected to generate a preliminary matrix of molecular counts for each cell. Then we removed CBs 1) with excessive (> 10%) molecules expressed from mitochondrial genome, 2) where the ratio of molecules aligning to whole gene bodies (including introns) to molecules aligning exclusively to exons was >1.5, and 3) for which the average number of reads per molecule or average number of molecules per gene deviated by >2.5 standard deviations from the mean for a given sample.

### Microarray gene expression analysis

Informative probe clusters were assembled with the cleaner algorithm (Alvarez et al., 2009) and the expression data was summarized and normalized with the MAS5 algorithm as implemented in the R package affy-1.54.0 from Bioconductor (Gautier et al., 2004). Differences in sample distributions were removed with the robust spline normalization procedure implemented in the R package lumi-2.28.0 from Bioconductor (Du et al., 2007, 2008; Du et al., 2010; Lin et al., 2008).

### Regulatory networks and protein activity analysis

All 5 brain tumor regulatory networks were reverse engineered by ARACNe (Basso et al., 2005; Lachmann et al., 2016) with 100 bootstrap iterations using 1,813 transcription factors (genes annotated in Gene Ontology molecular function database, as GO:0003700, ‘transcription factor activity’, or as GO:0003677, ‘DNA binding’, and GO:0030528, ‘transcription regulator activity’, or as GO:00034677 and GO: 0045449, ‘regulation of transcription’), 969 transcriptional cofactors (a manually curated list, not overlapping with the transcription factor list, built upon genes annotated as GO:0003712, “transcription cofactor activity”, or GO:0030528 or GO:0045449), and 3370 signaling pathway related genes (annotated in GO biological process database as GO:0007165 “signal transduction” and in GO cellular component database as GO:0005622, “intracellular”, or GO:0005886, “plasma membrane”). Parameters were set to 0 DPI (Data Processing Inequality) tolerance and MI (Mutual Information) p-value threshold, *p* = 10^-8^. Protein activity profiles were inferred from expression matrix by integrating the 5 regulatory networks with the metaVIPER algorithm (Alvarez et al., 2016; Ding et al., 2018).

### Gene set enrichment analysis (GSEA)

Genes over expressed in each of the Phillips- and Wang-specific subtypes were downloaded from the original studies (Phillips et al., 2006; Wang et al., 2017). Sets of aberrantly activated proteins in subtypes (C1, C21, C22, and C3) inferred by cluster analysis of the single-cell profiles from dataset GSE57872 (Patel et al., 2014) were assembled by metaVIPER analysis. Specifically, MES and PN-specific protein sets were generated by t-test analysis of differentially activated proteins between the C1 (PN cells) and the C3 (MES cells) cluster, with a Bonferroni corrected threshold p < 10^-10^. To identify PRO-specific proteins, we integrated the ranked list of proteins differentially activated between (a) the C21 and C1 clusters (proliferative and quiescent PN cells) and (b) the C22 and C3 clusters (proliferative and quiescent MES cells). Integration was performed by reciprocal GSEA leading edge analysis. Specifically, we first identified highly differentially active proteins in the C21 vs. C1 clusters, (*p* < 10^-10^ by t-test analysis, Bonferroni-corrected) and computed their GSEA enrichment in protein differentially active in the C22 vs. C3 clusters. We then identified highly differentially active proteins in the C22 vs. C3 clusters, (*p* < 10^-10^ by t-test analysis, Bonferroni-corrected) and computed the reciprocal GSEA enrichment in protein differentially active in the C21 vs. C1 clusters. Since both analyses were highly statistically significant (*p_C21→C1_* = 2.8E-41 and p_C22→C3_ = *p* = 2.8E-18, respectively), suggesting a highly-conserved repertoire of differentially activated proteins, we combined the proteins in the leading edge of both analyses to generate a final list of PRO-specific proteins. For these analyses, (see Fig3SupTable-2).

### Synthetic bulk analysis

Synthetic bulk samples were constructing by summing the reads from *n* = 40 single-cell samples from the GSE57872 (Patel et al., 2014) dataset picked randomly from clusters C1 (PN cells), C3 (MES cells); C21 (PPN cells) and C22 (PMES cells) with probabilities *p_C_*_1_, *p_C_*_12_, *p_C_*_22_ and *p_C_*_3_ determined by two fractional numbers *p_PN_*_/*MES*_ ∈ [0,1] and *p_PRO_*_/*QUI*_ ∈ [0,1] (represented on the x and y axis of Figure 5DE), such that:

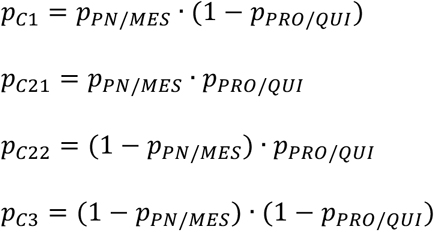

Such that *p_C_*_1_+*p_C_*_12_+*p_C_*_22_+*p_C_*_3_ = 1. We then annotated each synthetic bulk samples according to either the Wang (Figure 5D) or the Phillips (Figure 5E) classification, by GSEA analysis.

### OncoTreat analysis

To assess the sensitivity of single cells representative of each quadrant to specific agents, we performed OncoTreat analysis (Alvarez et al., 2018) on each cell profile in the GSE57872 dataset (Patel et al., 2014), using perturbational data generated by PLATESeq RNA-Seq profile analysis of GBM cell lines at 24h following treatment with etoposide, R04929097, and panobinostat. Specifically, OncoTreat assesses the ability of a drug to invert the activity of the master regulator proteins that mechanistically control a specific transcriptional state based on RNA-Seq following perturbational assays in cells representative of the tumor lineage. To effectively represent drug response in both mesenchymal and proneural cells, we performed drug perturbation assays in two complementary cell lines, including HF2597 (proneural) and U87 (mesenchymal). For each cell line, drug concentration was set to the drug’s 48h IC_20_ (i.e. highest sub-lethal dose). This prevents gene expression profiles of drug perturbation to be overly contaminated by cell stress and death programs and has been shown to be optimally suitable to reveal drug mechanisms of action (Woo et al., 2015). For each cell, OncoTreat p-values—assessing the ability of a drug to invert the activity of master regulator proteins of that cell, based on HF2597 and U87 cells perturbational data—were then integrated using the Stouffer’s method (Figure 7A-C). Interestingly, OncoTreat predictions from the two cell lines were in significant agreement across all single cells, suggesting a high reproducibility of the analysis.

### Single cell drug perturbation analysis

Protein activity profiles for drug- and vehicle-treated single cells (see section “single cell drug perturbation”) were inferred as previously described (see section “regulatory networks and protein activity analysis”). Based on such profiles, UMAP dimension reduction (Becht et al., 2018) was performed to project high-dimensional single-cell data on a 2D representation. Based on UMAP projection, DBSCAN clustering analysis (Ester et al., 1996) was performed, giving 4 major clusters, among which one non-tumor population was detected according to autosome copy number analysis: for each autosome, average expression of corresponding genes with expression rate greater the 10% were considered the “pseudo copy number”. Considering GBM hallmark CNV alterations in chromosome 7 (gain) and chromosome 10 (loss), we normalized the “pseudo copy number” of chromosome 7 and 10 against the average “pseudo copy number” of all other autosomes, as the representation of tumor/noon-tumor cells, following a similar rationale as described in (Patel et al., 2014). The other three tumor-related clusters were annotated as PN, MES and PRO respectively, based on GSEA analysis using single cell classifier (see section “gene set enrichment analysis”).

Finally, for each drug, Fisher’s Exact Text (FET) was used to quantify subtype-specific depletion at the single cell level. FET tables were computed as following, with P and Q representing the number of cells in Subtype #1 and #2, following drug treatment, and P’ and Q’ represent number of cells in Subtype #1 and #2 in vehicle-treated (DMSO) controls. Single cell subtype classification was determined by cluster analysis, as described above, including for PN, MES and PRO.

|  | Drug | DMSO |
| --- | --- | --- |
| Subtype #1 | P | $(P+Q) \times P'/(P'+Q')$ |
| Subtype #2 | Q | $(P+Q) \times Q'/(P'+Q')$ |

### Data and software availability

The scRNA-Seq data reported in this paper has been deposited in GEO under the accession number GSE107978 and GSE129671. All relevant data can also be downloaded from the NCI-CTD^2^ data portal (http://ctd2.nci.hih.gov).

## Author Contributions

AC and JNS conceived and oversaw the project. HD designed and performed all the analysis with help from MJA, and EOP, while PL performed the unsupervised cluster analyses. PSS provided biological interpretation of differentially active proteins. DMB, KKB and LH were responsible for all work related to single cell profile generation from the Mayo PDX models. JNB, PC, EFS, PU and AD procured patient-derived tumor surgical specimens and prepared, cultured, and performed ex-vivo drug treatment assays in patient-derived GBM explants. PAS and WZ profiled and analyzed single-cells from drug-treated explants, and generated the corresponding single-cell profiles. AK, FDC, and DD conceived the PDX validation assays and FDC and DD were responsible for its implementation and analysis. AC, HD, JNS, and PSS wrote the manuscript, with help from PL, DMB, PC, JNB, and PAS.

## Acknowledgments

We thank Erik P. Sulman and Roel G, W, Verhaak for annotating the Verhaak subtype of primary patient samples from which the mouse PDX models were derived. This work was supported by US National Institute of Health grants: R35 CA197745 (NCI) to AC for the single cell methodology development and analyses, Cancer Target Discovery and Development U01 CA217858 (NCI) to AC for the overall study design and bulk-tissue analyses and for the drug perturbation profiles. S10 OD012351 and S10 OD021764 for the High-Performance Computing infrastructure necessary to perform the analyses, R01 NS103473-01 (NCI) to PC, PS, and JNB, and P30 CA008748 (NCI) for the use of genomic and single cell shared resources.

## References

Akagi, T., Thoennissen, N.H., George, A., Crooks, G., Song, J.H., Okamoto, R., Nowak, D., Gombart, A.F., and Koeffler, H.P. (2010). In Vivo Deficiency of Both C/EBPβ and C/EBPε Results in Highly Defective Myeloid Differentiation and Lack of Cytokine Response. 5, e15419.

Alexander, Kristin, Yu, W., Derek, Ebru, Melissa, Joshua, Tonelli, M., Allison, Alan, et al. (2013). Calorie Restriction and SIRT3 Trigger Global Reprogramming of the Mitochondrial Protein Acetylome. 49, 186–199.

Alvarez, M.J., Sumazin, P., Rajbhandari, P., and Califano, A. (2009). Correlating measurements across samples improves accuracy of large-scale expression profile experiments. Genome Biol 10, R143.

Alvarez, M.J., Shen, Y., Giorgi, F.M., Lachmann, A., Ding, B.B., Ye, B.H., and Califano, A. (2016). Functional characterization of somatic mutations in cancer using network-based inference of protein activity. Nature genetics 48, 838–847.

Alvarez, M.J., Subramaniam, P.S., Tang, L.H., Grunn, A., Aburi, M., Rieckhof, G., Komissarova, E.V., Hagan, E.A., Bodei, L., Clemons, P.A., et al. (2018). A precision oncology approach to the pharmacological targeting of mechanistic dependencies in neuroendocrine tumors. Nature genetics 50, 979–989.

Amal, Chansey, Cheng, H., Thomas, Gian, Syam, Dale, Tirode, F., Mathers, J., Khan, D., et al. (2015). Translational Activation of HIF1α by YB-1 Promotes Sarcoma Metastasis. 27, 682–697.

Aubry, S., Shin, W., Crary, J.F., Lefort, R., Qureshi, Y.H., Lefebvre, C., Califano, A., and Shelanski, M.L. (2015). Assembly and interrogation of Alzheimer’s disease genetic networks reveal novel regulators of progression. PLoS One 10, e0120352.

Balci, T., Yilmaz Susluer, S., Kayabasi, C., Ozmen Yelken, B., Biray Avci, C., and Gunduz, C. (2016). Analysis of dysregulated long non-coding RNA expressions in glioblastoma cells. 590, 120–122.

Basso, K., Margolin, A.A., Stolovitzky, G., Klein, U., Dalla-Favera, R., and Califano, A. (2005). Reverse engineering of regulatory networks in human B cells. Nature genetics 37, 382–390.

Baumann, G., Travieso, L., Liebl, D.J., and Theus, M.H. (2013). Pronounced hypoxia in the subventricular zone following traumatic brain injury and the neural stem/progenitor cell response. 238, 830–841.

Becht, E., McInnes, L., Healy, J., Dutertre, C.A., Kwok, I.W.H., Ng, L.G., Ginhoux, F., and Newell, E.W. (2018). Dimensionality reduction for visualizing single-cell data using UMAP. Nat Biotechnol.

Bedi, R., Du, J., Sharma, A.K., Gomes, I., and Ackerman, S.J. (2008). Human C/EBP-activator and repressor isoforms differentially reprogram myeloid lineage commitment and differentiation. 113, 317–327.

Bellido, T., O’Brien, C.A., Roberson, P.K., and Manolagas, S.C. (1998). Transcriptional activation of the p21(WAF1,CIP1,SDI1) gene by interleukin-6 type cytokines - A prerequisite for their pro-differentiating and anti-apoptotic effects on human osteoblastic cells. Journal of Biological Chemistry 273, 21137–21144.

Bergsland, M., Ramskold, D., Zaouter, C., Klum, S., Sandberg, R., and Muhr, J. (2011). Sequentially acting Sox transcription factors in neural lineage development. Genes & development 25, 2453–2464.

Billin, A.N., and Ayer, D.E. (2006). The Mlx network: evidence for a parallel Max-like transcriptional network that regulates energy metabolism. Curr Top Microbiol Immunol 302, 255–278.

Bisikirska, B., Bansal, M., Shen, Y., Teruya-Feldstein, J., Chaganti, R., and Califano, A. (2016). Elucidation and Pharmacological Targeting of Novel Molecular Drivers of Follicular Lymphoma Progression. Cancer Res 76, 664–674.

Brichta, L., Shin, W., Jackson-Lewis, V., Blesa, J., Yap, E.L., Walker, Z., Zhang, J., Roussarie, J.P., Alvarez, M.J., Califano, A., et al. (2015). Identification of neurodegenerative factors using translatome-regulatory network analysis. Nat Neurosci 18, 1325–1333.

Bulleid, N.J., and Ellgaard, L. (2011). Multiple ways to make disulfides. Trends in Biochemical Sciences 36, 485–492.

Burrell, R.A., McGranahan, N., Bartek, J., and Swanton, C. (2013). The causes and consequences of genetic heterogeneity in cancer evolution. Nature 501, 338–345.

Butow, R.A., and Avadhani, N.G. (2004). Mitochondrial Signaling. 14, 1–15.

Califano, A., and Alvarez, M.J. (2017). The recurrent architecture of tumour initiation, progression and drug sensitivity. Nat Rev Cancer 17, 116–130.

Carro, M.S., Lim, W.K., Alvarez, M.J., Bollo, R.J., Zhao, X., Snyder, E.Y., Sulman, E.P., Anne, S.L., Doetsch, F., Colman, H., et al. (2010). The transcriptional network for mesenchymal transformation of brain tumours. Nature 463, 318–325.

Carlson, B.L., Pokorny, J.L., Schroeder, M.A., and Sarkaria, J.N. (2011). Establishment, maintenance and in vitro and in vivo applications of primary human glioblastoma multiforme (GBM) xenograft models for translational biology studies and drug discovery. Current protocols in pharmacology / editorial board, SJ Enna Chapter 14, Unit 14 16.

Carlson, M., and Maintainer, B.P. (2015). TxDb.Hsapiens.UCSC.hg19.knownGene: Annotation package for TxDb object(s) (Bioconductor), pp. R package.

Catarina, Steinmann, V., Thomas, Jais, A., Esterbauer, H., and Juergen (2014). Ecdysone and Mediator Change Energy Metabolism to Terminate Proliferation in Drosophila Neural Stem Cells. 158, 874–888.

Ceccarelli, M., Floris, Tathiane, Thais, Sofie, Bradley, Morozova, O., Newton, Y., Radenbaugh, A., Stefano, et al. (2016). Molecular Profiling Reveals Biologically Discrete Subsets and Pathways of Progression in Diffuse Glioma. Cell 164, 550–563.

Chandran, U.R., Luthra, S., Santana-Santos, L., Mao, P., Kim, S.H., Minata, M., Li, J., Benos, P.V., DeWang, M., Hu, B., et al. (2015). Gene expression profiling distinguishes proneural glioma stem cells from mesenchymal glioma stem cells. Genom Data 5, 333–336.

Charrad, R.S., Li, Y., Delpech, B., Balitrand, N., Clay, D., Jasmin, C., Chomienne, C., and Smadja-Joffe, F. (1999). Ligation of the CD44 adhesion molecule reverses blockage of differentiation in human acute myeloid leukemia. Nat Med 5, 669–676.

Cheedipudi, S., Puri, D., Saleh, A., Gala, H.P., Rumman, M., Pillai, M.S., Sreenivas, P., Arora, R., Sellathurai, J., Schroder, H.D., et al. (2015). A fine balance: epigenetic control of cellular quiescence by the tumor suppressor PRDM2/RIZ at a bivalent domain in the cyclin a gene. Nucleic Acids Res 43, 6236–6256.

Chen, J.C., Alvarez, M.J., Talos, F., Dhruv, H., Rieckhof, G.E., Iyer, A., Diefes, K.L., Aldape, K., Berens, M., Shen, M.M., et al. (2014). Identification of Causal Genetic Drivers of Human Disease through Systems-Level Analysis of Regulatory Networks. Cell 159, 402–414.

Chen, S.-H., and Benveniste, E.N. (2004). Oncostatin M: a pleiotropic cytokine in the central nervous system. 15, 379–391.

Christie, K.J., Emery, B., Denham, M., Bujalka, H., Cate, H.S., and Turnley, A.M. (2013). Transcriptional Regulation and Specification of Neural Stem Cells. In Transcriptional and Translational Regulation of Stem Cells, G. Hime, and H. Abud, eds. (Dordrecht: Springer Netherlands), pp. 129–155.

Climente-Gonzalez, H., Porta-Pardo, E., Godzik, A., and Eyras, E. (2017). The Functional Impact of Alternative Splicing in Cancer. Cell Rep 20, 2215–2226.

Close, P., Gillard, M., Ladang, A., Jiang, Z., Papuga, J., Hawkes, N., Nguyen, L., Chapelle, J.P., Bouillenne, F., Svejstrup, J., et al. (2012). DERP6 (ELP5) and C3ORF75 (ELP6) Regulate Tumorigenicity and Migration of Melanoma Cells as Subunits of Elongator. 287, 32535–32545.

Crabtree, G.R. (2001). Calcium, calcineurin, and the control of transcription. J Biol Chem 276, 2313–2316.

Crabtree, G.R., and Schreiber, S.L. (2009). SnapShot: Ca2+-Calcineurin-NFATSignaling. 138, 210–210.e211.

Cui, J., Shi, M., Xie, D., Wei, D., Jia, Z., Zheng, S., Gao, Y., Huang, S., and Xie, K. (2014). FOXM1 Promotes the Warburg Effect and Pancreatic Cancer Progression via Transactivation of LDHA Expression. 20, 2595–2606.

Dang, C.V. (2011). Therapeutic Targeting of Myc-Reprogrammed Cancer Cell Metabolism. 76, 369–374.

Darmanis, S., Sloan, S.A., Croote, D., Mignardi, M., Chernikova, S., Samghababi, P., Zhang, Y., Neff, N., Kowarsky, M., Caneda, C., et al. (2017). Single-Cell RNA-Seq Analysis of Infiltrating Neoplastic Cells at the Migrating Front of Human Glioblastoma. Cell Rep 21, 1399–1410.

David, G. (2012). Regulation of oncogene-induced cell cycle exit and senescence by chromatin modifiers. 13, 992–1000.

Della Gatta, G., Palomero, T., Perez-Garcia, A., Ambesi-Impiombato, A., Bansal, M., Carpenter, Z.W., De Keersmaecker, K., Sole, X., Xu, L., Paietta, E., et al. (2012). Reverse engineering of TLX oncogenic transcriptional networks identifies RUNX1 as tumor suppressor in T-ALL. Nat Med 18, 436–440.

Di Croce, L., and Helin, K. (2013). Transcriptional regulation by Polycomb group proteins. Nat Struct Mol Biol 20, 1147–1155.

Ding, H., Douglass, E.F., Jr., Sonabend, A.M., Mela, A., Bose, S., Gonzalez, C., Canoll, P.D., Sims, P.A., Alvarez, M.J., and Califano, A. (2018a). Quantitative assessment of protein activity in orphan tissues and single cells using the metaVIPER algorithm. Nat Commun 9, 1471.

Ding, H., Wang, W., and Califano, A. (2018b). iterClust: a statistical framework for iterative clustering analysis. Bioinformatics 34, 2865–2866.

Diolaiti, D., McFerrin, L., Carroll, P.A., and Eisenman, R.N. (2015). Functional interactions among members of the MAX and MLX transcriptional network during oncogenesis. Bba-Gene Regul Mech 1849, 484–500.

Du, P., Kibbe, W.A., and Lin, S.M. (2007). nuID: a universal naming scheme of oligonucleotides for illumina, affymetrix, and other microarrays. Biol Direct 2, 16.

Du, P., Kibbe, W.A., and Lin, S.M. (2008). lumi: a pipeline for processing Illumina microarray. Bioinformatics 24, 1547–1548.

Du, P., Zhang, X., Huang, C.C., Jafari, N., Kibbe, W.A., Hou, L., and Lin, S.M. (2010). Comparison of Beta-value and M-value methods for quantifying methylation levels by microarray analysis. BMC bioinformatics 11, 587.

Dufu, K., Livingstone, M.J., Seebacher, J., Gygi, S.P., Wilson, S.A., and Reed, R. (2010). ATP is required for interactions between UAP56 and two conserved mRNA export proteins, Aly and CIP29, to assemble the TREX complex. 24, 2043–2053.

Elaimy, A.L., and Mercurio, A.M. (2018). Convergence of VEGF and YAP/TAZ signaling: Implications for angiogenesis and cancer biology. Sci Signal 11.

Eliseeva, I.A., Kim, E.R., Guryanov, S.G., Ovchinnikov, L.P., and Lyabin, D.N. (2011). Y-box-binding protein 1 (YB-1) and its functions. 76, 1402–1433.

Ester, M., Kriegel, H.P., Sander, J., and Xu, X. (1996). A density-based algorithm for discovering clusters in large spatial databases with noise. Kdd 96, 226–231.

Farcas, A.M., Blackledge, N.P., Sudbery, I., Long, H.K., McGouran, J.F., Rose, N.R., Lee, S., Sims, D., Cerase, A., Sheahan, T.W., et al. (2012). KDM2B links the Polycomb Repressive Complex 1 (PRC1) to recognition of CpG islands. Elife 1, e00205.

Fedele, M., Crescenzi, E., and Cerchia, L. (2017). The POZ/BTB and AT-Hook Containing Zinc Finger 1 (PATZ1) Transcription Regulator: Physiological Functions and Disease Involvement. Int J Mol Sci 18.

Fisher, R.P. (2018). Taking Aim at Glycolysis with CDK8 Inhibitors. Trends in Endocrinology & Metabolism 29, 281–282.

Frei, C., Galloni, M., Hafen, E., and Edgar, B.A. (2005). The Drosophila mitochondrial ribosomal protein mRpL12 is required for Cyclin D/Cdk4-driven growth. 24, 623–634.

Freije, W.A., Castro-Vargas, F.E., Fang, Z., Horvath, S., Cloughesy, T., Liau, L.M., Mischel, P.S., and Nelson, S.F. (2004). Gene Expression Profiling of Gliomas Strongly Predicts Survival. Cancer Research 64, 6503–6510.

Frohn, A., Eberl, H.C., Stohr, J., Glasmacher, E., Rudel, S., Heissmeyer, V., Mann, M., and Meister, G. (2012). Dicer-dependent and -independent Argonaute2 Protein Interaction Networks in Mammalian Cells. 11, 1442–1456.

Gahwiler, B.H., Capogna, M., Debanne, D., McKinney, R.A., and Thompson, S.M. (1997). Organotypic slice cultures: a technique has come of age. Trends Neurosci 20, 471–477.

Gill, B.J., Pisapia, D.J., Malone, H.R., Goldstein, H., Lei, L., Sonabend, A., Yun, J., Samanamud, J., Sims, J.S., Banu, M., et al. (2014). MRI-localized biopsies reveal subtype-specific differences in molecular and cellular composition at the margins of glioblastoma. Proc Natl Acad Sci U S A 111, 12550–12555.

Gupta, S.K., Kizilbash, S.H., Carlson, B.L., Mladek, A.C., Boakye-Agyeman, F., Bakken, K.K., Pokorny, J.L., Schroeder, M.A., Decker, P.A., Cen, L., et al. (2016). Delineation of MGMT Hypermethylation as a Biomarker for Veliparib-Mediated Temozolomide-Sensitizing Therapy of Glioblastoma. J Natl Cancer Inst 108.

Gupta, S.K., Mladek, A.C., Carlson, B.L., Boakye-Agyeman, F., Bakken, K.K., Kizilbash, S.H., Schroeder, M.A., Reid, J., and Sarkaria, J.N. (2014). Discordant in vitro and in vivo chemopotentiating effects of the PARP inhibitor veliparib in temozolomide-sensitive versus - resistant glioblastoma multiforme xenografts. Clin Cancer Res 20, 3730–3741.

Hawer, H., Hammermeister, A., Ravichandran, K., Glatt, S., Schaffrath, R., and Klassen, R. (2018). Roles of Elongator Dependent tRNA Modification Pathways in Neurodegeneration and Cancer. Genes 10, 19.

He, J., Shen, L., Wan, M., Taranova, O., Wu, H., and Zhang, Y. (2013). Kdm2b maintains murine embryonic stem cell status by recruiting PRC1 complex to CpG islands of developmental genes. 15, 373–384.

Hsieh, J. (2012). Orchestrating transcriptional control of adult neurogenesis. Genes & development 26, 1010–1021.

Hsieh, J., and Zhao, X. (2016). Genetics and Epigenetics in Adult Neurogenesis. Cold Spring Harb Perspect Biol 8.

Hsu, T.Y.T., Simon, L.M., Neill, N.J., Marcotte, R., Sayad, A., Bland, C.S., Echeverria, G.V., Sun, T., Kurley, S.J., Tyagi, S., et al. (2015). The spliceosome is a therapeutic vulnerability in MYC-driven cancer. 525, 384–388.

Hu, J., Liu, Y.L., Piao, S.L., Yang, D.D., Yang, Y.M., and Cai, L. (2012). Expression patterns of USP22 and potential targets BMI-1, PTEN, p-AKT in non-small-cell lung cancer. Lung Cancer 77, 593–599.

Ikeda, M., and Toyoshima, F. (2017). Dormant Pluripotent Cells Emerge during Neural Differentiation of Embryonic Stem Cells in a FoxO3-Dependent Manner. Molecular and Cellular Biology 37, MCB.00417-00416.

Ikiz, B., Alvarez, M.J., Re, D.B., Le Verche, V., Politi, K., Lotti, F., Phani, S., Pradhan, R., Yu, C., Croft, G.F., et al. (2015). The Regulatory Machinery of Neurodegeneration in In Vitro Models of Amyotrophic Lateral Sclerosis. Cell Rep 12, 335–345.

Ilarregui, J.M., Croci, D.O., Bianco, G.A., Toscano, M.A., Salatino, M., Vermeulen, M.E., Geffner, J.R., and Rabinovich, G.A. (2009). Tolerogenic signals delivered by dendritic cells to T cells through a galectin-1-driven immunoregulatory circuit involving interleukin 27 and interleukin 10. 10, 981–991.

Jahani-Asl, A., Yin, H., Soleimani, V.D., Haque, T., Luchman, H.A., Chang, N.C., Sincennes, M.-C., Puram, S.V., Scott, A.M., Lorimer, I.A.J., et al. (2016). Control of glioblastoma tumorigenesis by feed-forward cytokine signaling. 19, 798–806.

Jin, L.Q., Hope, K.J., Zhai, Q.L., Smadja-Joffe, F., and Dick, J.E. (2006). Targeting of CD44 eradicates human acute myeloid leukemic stem cells. Nature Medicine 12, 1167–1174.

Johansson, E., Grassi, E.S., Pantazopoulou, V., Tong, B., Lindgren, D., Berg, T.J., Pietras, E.J., Axelson, H., and Pietras, A. (2017). CD44 Interacts with HIF-2α to Modulate the Hypoxic Phenotype of Perinecrotic and Perivascular Glioma Cells. Cell Reports 20, 1641–1653.

Jones, M., Chase, J., Brinkmeier, M., Xu, J., Weinberg, D.N., Schira, J., Friedman, A., Malek, S., Grembecka, J., Cierpicki, T., et al. (2015). Ash1l controls quiescence and self-renewal potential in hematopoietic stem cells. J Clin Invest 125, 2007–2020.

Kanellopoulou, C., Gilpatrick, T., Kilaru, G., Burr, P., Nguyen, C.K., Morawski, A., Lenardo, M.J., and Muljo, S.A. (2015). Reprogramming of Polycomb-Mediated Gene Silencing in Embryonic Stem Cells by the miR-290 Family and the Methyltransferase Ash1l. Stem Cell Reports 5, 971–978.

Kharchenko, P.V., Silberstein, L., and Scadden, D.T. (2014). Bayesian approach to single-cell differential expression analysis. Nat Methods 11, 740–742.

Kim, H.S., Woolard, K., Lai, C., Bauer, P.O., Maric, D., Song, H., Li, A., Kotliarova, S., Zhang, W., and Fine, H.A. (2012). Gliomagenesis Arising from Pten- and Ink4a/Arf-Deficient Neural Progenitor Cells Is Mediated by the p53-Fbxw7/Cdc4 Pathway, Which Controls c-Myc. 72, 6065–6075.

Kobayashi, A., Yamagiwa, H., Hoshino, H., Muto, A., Sato, K., Morita, M., Hayashi, N., Yamamoto, M., and Igarashi, K. (2000). A Combinatorial Code for Gene Expression Generated by Transcription Factor Bach2 and MAZR (MAZ-Related Factor) through the BTB/POZ Domain. 20, 1733–1746.

Kohling, R., Qu, M., Zilles, K., and Speckmann, E.J. (1999). Current-source-density profiles associated with sharp waves in human epileptic neocortical tissue. Neuroscience 94, 1039–1050.

Kohyama, J., Kojima, T., Takatsuka, E., Yamashita, T., Namiki, J., Hsieh, J., Gage, F.H., Namihira, M., Okano, H., Sawamoto, K., et al. (2008). Epigenetic regulation of neural cell differentiation plasticity in the adult mammalian brain. Proc Natl Acad Sci U S A 105, 18012–18017.

Kolodziejczyk, A.A., Kim, J.K., Tsang, J.C., Ilicic, T., Henriksson, J., Natarajan, K.N., Tuck, A.C., Gao, X., Buhler, M., Liu, P., et al. (2015). Single Cell RNA-Sequencing of Pluripotent States Unlocks Modular Transcriptional Variation. Cell Stem Cell 17, 471–485.

Krishnan, H., Rayes, J., Miyashita, T., Ishii, G., Retzbach, E.P., Sheehan, S.A., Takemoto, A., Chang, Y.W., Yoneda, K., Asai, J., et al. (2018). Podoplanin: An emerging cancer biomarker and therapeutic target. Cancer Sci 109, 1292–1299.

Kushwaha, R., Jagadish, N., Kustagi, M., Tomishima, M.J., Mendiratta, G., Bansal, M., Kim, H.R., Sumazin, P., Alvarez, M.J., Lefebvre, C., et al. (2015). Interrogation of a context-specific transcription factor network identifies novel regulators of pluripotency. Stem Cells 33, 367–377.

Lachmann, A., Giorgi, F.M., Lopez, G., and Califano, A. (2016). ARACNe-AP: gene network reverse engineering through adaptive partitioning inference of mutual information. Bioinformatics 32, 2233–2235.

Langmead, B., and Salzberg, S.L. (2012). Fast gapped-read alignment with Bowtie 2. Nat Methods 9, 357–359.

Lasham, A., Cristin, Adele, Sandra, and Antony (2013). YB-1: oncoprotein, prognostic marker and therapeutic target? Biochemical Journal 449, 11–23.

Lawrence, M., Huber, W., Pages, H., Aboyoun, P., Carlson, M., Gentleman, R., Morgan, M.T., and Carey, V.J. (2013). Software for computing and annotating genomic ranges. PLoS Comput Biol 9, e1003118.

Lefebvre, C., Rajbhandari, P., Alvarez, M.J., Bandaru, P., Lim, W.K., Sato, M., Wang, K., Sumazin, P., Kustagi, M., Bisikirska, B.C., et al. (2010). A human B-cell interactome identifies MYB and FOXM1 as master regulators of proliferation in germinal centers. Molecular systems biology 6, 377.

Levitin, H.M., Yuan, J., Cheng, Y.L., Ruiz, F.J., Bush, E.C., Bruce, J.N., Canoll, P., Iavarone, A., Lasorella, A., Blei, D.M., et al. (2019). De novo gene signature identification from single-cell RNA-seq with hierarchical Poisson factorization. Molecular systems biology 15, e8557.

Li, H., Handsaker, B., Wysoker, A., Fennell, T., Ruan, J., Homer, N., Marth, G., Abecasis, G., Durbin, R., and Genome Project Data Processing, S. (2009). The Sequence Alignment/Map format and SAMtools. Bioinformatics 25, 2078–2079.

Li, Q., Jia, H., Li, H., Dong, C., Wang, Y., and Zou, Z. (2016). LncRNA and mRNA expression profiles of glioblastoma multiforme (GBM) reveal the potential roles of lncRNAs in GBM pathogenesis.

Liang, Y., Diehn, M., Watson, N., Bollen, A.W., Aldape, K.D., Nicholas, M.K., Lamborn, K.R., Berger, M.S., Botstein, D., Brown, P.O., et al. (2005). Gene expression profiling reveals molecularly and clinically distinct subtypes of glioblastoma multiforme. Proceedings of the National Academy of Sciences 102, 5814–5819.

Lim, D.A., Huang, Y.C., Swigut, T., Mirick, A.L., Garcia-Verdugo, J.M., Wysocka, J., Ernst, P., and Alvarez-Buylla, A. (2009). Chromatin remodelling factor Mll1 is essential for neurogenesis from postnatal neural stem cells. Nature 458, 529–533.

Lin, S.M., Du, P., Huber, W., and Kibbe, W.A. (2008). Model-based variance-stabilizing transformation for Illumina microarray data. Nucleic Acids Res 36, e11.

Lundquist, A., Barre, B., Bienvenu, F., Hermann, J., Avril, S., and Coqueret, O. (2003). Kaposi sarcoma-associated viral cyclin K overrides cell growth inhibition mediated by oncostatin M through STAT3 inhibition. Blood 101, 4070–4077.

Lyabin, D.N., Eliseeva, I.A., and Ovchinnikov, L.P. (2014). YB-1 protein: functions and regulation. 5, 95–110.

Mao, H., Lebrun, D.G., Yang, J., Zhu, V.F., and Li, M. (2012). Deregulated Signaling Pathways in Glioblastoma Multiforme: Molecular Mechanisms and Therapeutic Targets. Cancer Investigation 30, 48–56.

Mao, P., Joshi, K., Li, J., Kim, S.H., Li, P., Santana-Santos, L., Luthra, S., Chandran, U.R., Benos, P.V., Smith, L., et al. (2013). Mesenchymal glioma stem cells are maintained by activated glycolytic metabolism involving aldehyde dehydrogenase 1A3. Proc Natl Acad Sci U S A 110, 8644–8649.

Marcelino Meliso, F., Hubert, C.G., Favoretto Galante, P.A., and Penalva, L.O. (2017). RNA processing as an alternative route to attack glioblastoma. Human Genetics 136, 1129–1141.

Matus, D.Q., Lohmer, L.L., Kelley, L.C., Schindler, A.J., Kohrman, A.Q., Barkoulas, M., Zhang, W., Chi, Q., and Sherwood, D.R. (2015). Invasive Cell Fate Requires G1 Cell-Cycle Arrest and Histone Deacetylase-Mediated Changes in Gene Expression. Dev Cell 35, 162–174.

Maurer, C., Holmstrom, S.R., He, J., Laise, P., Su, T., Ahmed, A., Hibshoosh, H., Chabot, J.A., Oberstein, P.E., Sepulveda, A.R., et al. (2019). Experimental microdissection enables functional harmonisation of pancreatic cancer subtypes. Gut.

Miyazaki, H., Higashimoto, K., Yada, Y., Endo, T.A., Sharif, J., Komori, T., Matsuda, M., Koseki, Y., Nakayama, M., Soejima, H., et al. (2013). Ash1l methylates Lys36 of histone H3 independently of transcriptional elongation to counteract polycomb silencing. PLoS Genet 9, e1003897.

Mohan, S., Bonni, A., and Jahani-Asl, A. (2017). Targeting OSMR in glioma stem cells. Oncotarget 8.

Mooney, K.L., Choy, W., Sidhu, S., Pelargos, P., Bui, T.T., Voth, B., Barnette, N., and Yang, I. (2016). The role of CD44 in glioblastoma multiforme. 34, 1–5.

Murat, A., Migliavacca, E., Gorlia, T., Lambiv, W.L., Shay, T., Hamou, M.-F., De Tribolet, N., Regli, L., Wick, W., Kouwenhoven, M.C.M., et al. (2008). Stem Cell–Related “Self-Renewal” Signature and High Epidermal Growth Factor Receptor Expression Associated With Resistance to Concomitant Chemoradiotherapy in Glioblastoma. Journal of Clinical Oncology 26, 3015–3024.

Natesh, K., Bhosale, D., Desai, A., Chandrika, G., Pujari, R., Jagtap, J., Chugh, A., Ranade, D., and Shastry, P. (2015). Oncostatin-M Differentially Regulates Mesenchymal and Proneural Signature Genes in Gliomas via STAT3 Signaling. 17, 225–237.

Nutt, C.L., Mani, D.R., Betensky, R.A., Tamayo, P., Cairncross, J.G., Ladd, C., Pohl, U., Hartmann, C., McLaughlin, M.E., Batchelor, T.T., et al. (2003). Gene expression-based classification of malignant gliomas correlates better with survival than histological classification. Cancer Research 63, 1602–1607.

Ohgaki, H., and Kleihues, P. (2005). Epidemiology and etiology of gliomas. 109, 93–108.

Okuma, A., Hanyu, A., Watanabe, S., and Hara, E. (2017). p16(Ink4a) and p21(Cip1/Waf1) promote tumour growth by enhancing myeloid-derived suppressor cells chemotaxis. Nat Commun 8, 2050.

Patel, A.P., Tirosh, I., Trombetta, J.J., Shalek, A.K., Gillespie, S.M., Wakimoto, H., Cahill, D.P., Nahed, B.V., Curry, W.T., Martuza, R.L., et al. (2014). Single-cell RNA-seq highlights intratumoral heterogeneity in primary glioblastoma. Science 344, 1396–1401.

Peng, Z., Liu, C., and Wu, M. (2018). New insights into long noncoding RNAs and their roles in glioma. Molecular Cancer 17.

Pfau, R., Tzatsos, A., Kampranis, S.C., Serebrennikova, O.B., Bear, S.E., and Tsichlis, P.N. (2008). Members of a family of JmjC domain-containing oncoproteins immortalize embryonic fibroblasts via a JmjC domain-dependent process. 105, 1907–1912.

Phillips, H.S., Kharbanda, S., Chen, R., Forrest, W.F., Soriano, R.H., Wu, T.D., Misra, A., Nigro, J.M., Colman, H., Soroceanu, L., et al. (2006). Molecular subclasses of high-grade glioma predict prognosis, delineate a pattern of disease progression, and resemble stages in neurogenesis. Cancer cell 9, 157–173.

Rajalingam, K., and Rudel, T. (2005). Ras-Raf Signaling Needs Prohibitin. 4, 1503–1505.

Rapino, F., and Close, P. (2018). Wobble uridine tRNA modification: a new vulnerability of refractory melanoma. Molecular & Cellular Oncology, 1–3.

Raveney, B.J.E., Copland, D.A., Calder, C.J., Dick, A.D., and Nicholson, L.B. (2010). TNFR1 signalling is a critical checkpoint for developing macrophages that control of T-cell proliferation. 131, 340–349.

Renault, V.M., Rafalski, V.A., Morgan, A.A., Salih, D.A., Brett, J.O., Webb, A.E., Villeda, S.A., Thekkat, P.U., Guillerey, C., Denko, N.C., et al. (2009). FoxO3 regulates neural stem cell homeostasis. Cell Stem Cell 5, 527–539.

Repovic, P., Fears, C.Y., Gladson, C.L., and Benveniste, E.N. (2003). Oncostatin-M induction of vascular endothelial growth factor expression in astroglioma cells. 22, 8117–8124.

Repunte-Canonigo, V., Shin, W., Vendruscolo, L.F., Lefebvre, C., van der Stap, L., Kawamura, T., Schlosburg, J.E., Alvarez, M., Koob, G.F., Califano, A., et al. (2015). Identifying candidate drivers of alcohol dependence-induced excessive drinking by assembly and interrogation of brain-specific regulatory networks. Genome Biol 16, 68.

Rodriguez-Barrueco, R., Yu, J., Saucedo-Cuevas, L.P., Olivan, M., Llobet-Navas, D., Putcha, P., Castro, V., Murga-Penas, E.M., Collazo-Lorduy, A., Castillo-Martin, M., et al. (2015). Inhibition of the autocrine IL-6-JAK2-STAT3-calprotectin axis as targeted therapy for HR-/HER2+ breast cancers. Genes & development 29, 1631–1648.

Rousseeuw, P.J. (1987). Silhouettes - a Graphical Aid to the Interpretation and Validation of Cluster-Analysis. J Comput Appl Math 20, 53–65.

Sacco, J.J., Yau, T.Y., Darling, S., Patel, V., Liu, H., Urbe, S., Clague, M.J., and Coulson, J.M. (2014). The deubiquitylase Ataxin-3 restricts PTEN transcription in lung cancer cells. Oncogene 33, 4265–4272.

Sade-Feldman, M., Kanterman, J., Ish-Shalom, E., Elnekave, M., Horwitz, E., and Baniyash, M. (2013). Tumor Necrosis Factor-α Blocks Differentiation and Enhances Suppressive Activity of Immature Myeloid Cells during Chronic Inflammation. 38, 541–554.

Saha, D., Banerjee, S., Bashir, S., and Vijayraghavan, U. (2012). Context dependent splicing functions of Bud31/Ycr063w define its role in budding and cell cycle progression. 424, 579–585.

Sans, C.L., Satterwhite, D.J., Stoltzman, C.A., Breen, K.T., and Ayer, D.E. (2006). MondoA-Mlx Heterodimers Are Candidate Sensors of Cellular Energy Status: Mitochondrial Localization and Direct Regulation of Glycolysis. Molecular and Cellular Biology 26, 4863–4871.

Shang, R., Pu, M., Li, Y., and Wang, D. (2017). FOXM1 regulates glycolysis in hepatocellular carcinoma by transactivating glucose transporter 1 expression. Oncology Reports 37, 2261–2269.

Shiina, S., Ohno, M., Ohka, F., Kuramitsu, S., Yamamichi, A., Kato, A., Motomura, K., Tanahashi, K., Yamamoto, T., Watanabe, R., et al. (2016). CAR T Cells Targeting Podoplanin Reduce Orthotopic Glioblastomas in Mouse Brains.

Shimizu, F., Hovinga, K.E., Metzner, M., Soulet, D., and Tabar, V. (2011). Organotypic explant culture of glioblastoma multiforme and subsequent single-cell suspension. Curr Protoc Stem Cell Biol Chapter 3, Unit3 5.

Shurin, G.V., Ferris, R., Tourkova, I.L., Perez, L., Lokshin, A., Balkir, L., Collins, B., Chatta, G.S., and Shurin, M.R. (2005). Loss of New Chemokine CXCL14 in Tumor Tissue Is Associated with Low Infiltration by Dendritic Cells (DC), while Restoration of Human CXCL14 Expression in Tumor Cells Causes Attraction of DC Both In Vitro and In Vivo. 174, 5490–5498.

Si, J., Yu, X., Zhang, Y., and DeWille, J.W. (2010). Myc interacts with Max and Miz1 to repress C/EBPdelta promoter activity and gene expression. Mol Cancer 9, 92.

Sonabend, A.M., Carminucci, A.S., Amendolara, B., Bansal, M., Leung, R., Lei, L., Realubit, R., Li, H., Karan, C., Yun, J., et al. (2014). Convection-enhanced delivery of etoposide is effective against murine proneural glioblastoma. Neuro Oncol 16, 1210–1219.

Stoltzman, C.A., Peterson, C.W., Breen, K.T., Muoio, D.M., Billin, A.N., and Ayer, D.E. (2008). Glucose sensing by MondoA:Mlx complexes: A role for hexokinases and direct regulation of thioredoxin-interacting protein expression. 105, 6912–6917.

Stoppini, L., Buchs, P.A., and Muller, D. (1991). A simple method for organotypic cultures of nervous tissue. J Neurosci Methods 37, 173–182.

Subramanian, A., Tamayo, P., Mootha, V.K., Mukherjee, S., Ebert, B.L., Gillette, M.A., Paulovich, A., Pomeroy, S.L., Golub, T.R., Lander, E.S., et al. (2005). Gene set enrichment analysis: a knowledge-based approach for interpreting genome-wide expression profiles. Proc Natl Acad Sci U S A 102, 15545–15550.

Suresh, P.S., Tsutsumi, R., and Venkatesh, T. (2018). YBX1 at the crossroads of non-coding transcriptome, exosomal, and cytoplasmic granular signaling. European Journal of Cell Biology 97, 163–167.

Szabo, P.A., Mendes Levitin, H., Miron, M., Snyder, M.E., Senda, T., Jinzhou, Y., Cheng, Y.L., and al., e. (2019). A single-cell reference map for human blood and tissue T cell activation reveals functional states in health and disease. bioRxiv

Talos, F., Mitrofanova, A., Bergren, S.K., Califano, A., and Shen, M.M. (2017). A computational systems approach identifies synergistic specification genes that facilitate lineage conversion to prostate tissue. Nat Commun 8, 14662.

Tawara, K., Scott, H., Emathinger, J., Ide, A., Fox, R., Greiner, D., Lajoie, D., Hedeen, D., Nandakumar, M., Oler, A.J., et al. (2019). Co-Expression of VEGF and IL-6 Family Cytokines is Associated with Decreased Survival in HER2 Negative Breast Cancer Patients: Subtype-Specific IL-6 Family Cytokine-Mediated VEGF Secretion. Translational Oncology 12, 245–255.

Theus, M.H., Ricard, J., Bethea, J.R., and Liebl, D.J. (2010). Ephb3 Inhibits the Expansion of Neural Progenitor Cells in the SVZ by Regulating p53 During Homeostasis and Following Traumatic Brain Injury. N/A-N/A.

Ting, J.T., Kalmbach, B., Chong, P., de Frates, R., Keene, C.D., Gwinn, R.P., Cobbs, C., Ko, A.L., Ojemann, J.G., Ellenbogen, R.G., et al. (2018). A robust ex vivo experimental platform for molecular-genetic dissection of adult human neocortical cell types and circuits. Sci Rep 8, 8407.

Tirosh, I., Venteicher, A.S., Hebert, C., Escalante, L.E., Patel, A.P., Yizhak, K., Fisher, J.M., Rodman, C., Mount, C., Filbin, M.G., et al. (2016). Single-cell RNA-seq supports a developmental hierarchy in human oligodendroglioma. Nature 539, 309–313.

Tsujimura, K., Abematsu, M., Kohyama, J., Namihira, M., and Nakashima, K. (2009). Neuronal differentiation of neural precursor cells is promoted by the methyl-CpG-binding protein MeCP2. Exp Neurol 219, 104–111.

Tzatsos, A., Pfau, R., Kampranis, S.C., and Tsichlis, P.N. (2009). Ndy1/KDM2B immortalizes mouse embryonic fibroblasts by repressing the Ink4a/Arf locus. 106, 2641–2646.

Vachtenheim, J.J., Lischke, R., and Vachtenheim, J. (2018). Siva-1 emerges as a tissue-specific oncogene beyond its classic role of a proapoptotic gene. OncoTargets and Therapy Volume 11, 6361–6367.

Van Nostrand, J.L., Brisac, A., Mello, S.S., Jacobs, S.B.R., Luong, R., and Attardi, L.D. (2015). The p53 Target Gene SIVA Enables Non-Small Cell Lung Cancer Development. 5, 622–635.

Verhaak, R.G.W., Hoadley, K.A., Purdom, E., Wang, V., Qi, Y., Wilkerson, M.D., Miller, C.R., Ding, L., Golub, T., Mesirov, J.P., et al. (2010). Integrated Genomic Analysis Identifies Clinically Relevant Subtypes of Glioblastoma Characterized by Abnormalities in PDGFRA, IDH1, EGFR, and NF1. Cancer cell 17, 98–110.

Wang, Q., Hu, B., Hu, X., Kim, H., Squatrito, M., Scarpace, L., deCarvalho, A.C., Lyu, S., Li, P., Li, Y., et al. (2017). Tumor Evolution of Glioma-Intrinsic Gene Expression Subtypes Associates with Immunological Changes in the Microenvironment. Cancer cell 32, 42–56 e46.

Wang, X., Bi, Y., Xue, L., Liao, J., Chen, X., Lu, Y., Zhang, Z., Wang, J., Liu, H., Yang, H., et al. (2015). The calcineurin-NFAT axis controls allograft immunity in myeloid-derived suppressor cells through reprogramming T cell differentiation. Mol Cell Biol 35, 598–609.

Wang, X., Zha, M., Zhao, X., Jiang, P., Du, W., Tam, A.Y.H., Mei, Y., and Wu, M. (2013). Siva1 inhibits p53 function by acting as an ARF E3 ubiquitin ligase. Nature Communications 4, 1551.

Wang, Y., Yun, Y., Wu, B., Wen, L., Wen, M., Yang, H., Zhao, L., Liu, W., Huang, S., Wen, N., et al. (2016). FOXM1 promotes reprogramming of glucose metabolism in epithelial ovarian cancer cells via activation of GLUT1 and HK2 transcription. Oncotarget 7, 47985–47997.

Wegner, M. (2011). SOX after SOX: SOXession regulates neurogenesis. 25, 2423–2428.

Wei, Y., Chiang, W.C., Sumpter, R., Jr., Mishra, P., and Levine, B. (2017). Prohibitin 2 Is an Inner Mitochondrial Membrane Mitophagy Receptor. Cell 168, 224–238 e210.

Wise, D.R., Deberardinis, R.J., Mancuso, A., Sayed, N., Zhang, X.Y., Pfeiffer, H.K., Nissim, I., Daikhin, E., Yudkoff, M., McMahon, S.B., et al. (2008). Myc regulates a transcriptional program that stimulates mitochondrial glutaminolysis and leads to glutamine addiction. Proceedings of the National Academy of Sciences 105, 18782–18787.

Woo, J.H., Shimoni, Y., Yang, W.S., Subramaniam, P., Iyer, A., Nicoletti, P., Rodriguez Martinez, M., Lopez, G., Mattioli, M., Realubit, R., et al. (2015). Elucidating Compound Mechanism of Action by Network Perturbation Analysis. Cell 162, 441–451.

Wu, S.-L., Fu, X., Huang, J., Jia, T.-T., Zong, F.-Y., Mu, S.-R., Zhu, H., Yan, Y., Qiu, S., Wu, Q., et al. (2015). Genome-wide analysis of YB-1-RNA interactions reveals a novel role of YB-1 in miRNA processing in glioblastoma multiforme. 43, 8516–8528.

Xu, L., Li, H., Wu, L., and Huang, S. (2017). YBX1 promotes tumor growth by elevating glycolysis in human bladder cancer. Oncotarget.

Yang, F., Li, Y., Zhang, Q., Tan, L., Peng, L., and Zhao, Y. (2018a). The Effect of Immunosuppressive Drugs on MDSCs in Transplantation. J Immunol Res 2018, 5414808.

Yang, J., Li, B., and He, Q.-Y. (2018b). Significance of prohibitin domain family in tumorigenesis and its implication in cancer diagnosis and treatment. Cell Death & Disease 9.

Yu, X., Wang, J., Wu, J., and Shi, Y. (2015). A systematic study of the cellular metabolic regulation of Jhdm1b in tumor cells. 11, 1867–1875.

Yuan, J., and Sims, P.A. (2016). An Automated Microwell Platform for Large-Scale Single Cell RNA-Seq. Scientific reports 6, 33883.

Yuan, J., Levitin, H.M., Frattini, V., Bush, E.C., Boyett, D.M., Samanamud, J., Ceccarelli, M., Dovas, A., Zanazzi, G., Canoll, P., et al. (2018). Single-cell transcriptome analysis of lineage diversity in high-grade glioma. Genome Med 10, 57.

Zhang, X.-Q., Kiang, K.M.-Y., Wang, Y.-C., Pu, J.K.-S., Ho, A., Cheng, S.Y., Lee, D., Zhang, P.-D., Chen, J.-J., Lui, W.-M., et al. (2015). IDH1 mutation-associated long non-coding RNA expression profile changes in glioma. 125, 253–263.

Zhao, X., Rong, L., Zhao, X., Li, X., Liu, X., Deng, J., Wu, H., Xu, X., Erben, U., Wu, P., et al. (2012). TNF signaling drives myeloid-derived suppressor cell accumulation. 122, 4094–4104.

Zheng, H., Ying, H., Yan, H., Kimmelman, A.C., Hiller, D.J., Chen, A.-J., Perry, S.R., Tonon, G., Chu, G.C., Ding, Z., et al. (2008). p53 and Pten control neural and glioma stem/progenitor cell renewal and differentiation. 455, 1129–1133.

Zheng, X., Boyer, L., Jin, M., Mertens, J., Kim, Y., Ma, L., Ma, L., Hamm, M., Gage, F.H., and Hunter, T. (2016). Metabolic reprogramming during neuronal differentiation from aerobic glycolysis to neuronal oxidative phosphorylation. Elife 5.

Zhuang, Y.J., Liao, Z.W., Yu, H.W., Song, X.L., Liu, Y., Shi, X.Y., Lin, X.D., and Zhou, T.C. (2015). ShRNA-mediated silencing of the ubiquitin-specific protease 22 gene restrained cell progression and affected the Akt pathway in nasopharyngeal carcinoma. Cancer Biol Ther 16, 88–96.

Zito, E. (2013). PRDX4, an Endoplasmic Reticulum-Localized Peroxiredoxin at the Crossroads Between Enzymatic Oxidative Protein Folding and Nonenzymatic Protein Oxidation. 18, 1666–1674.

