## Supplementary Figures for "Single-cell based elucidation of molecularly-distinct glioblastoma states and drug sensitivity"

### Fig1SupFig: General workflow

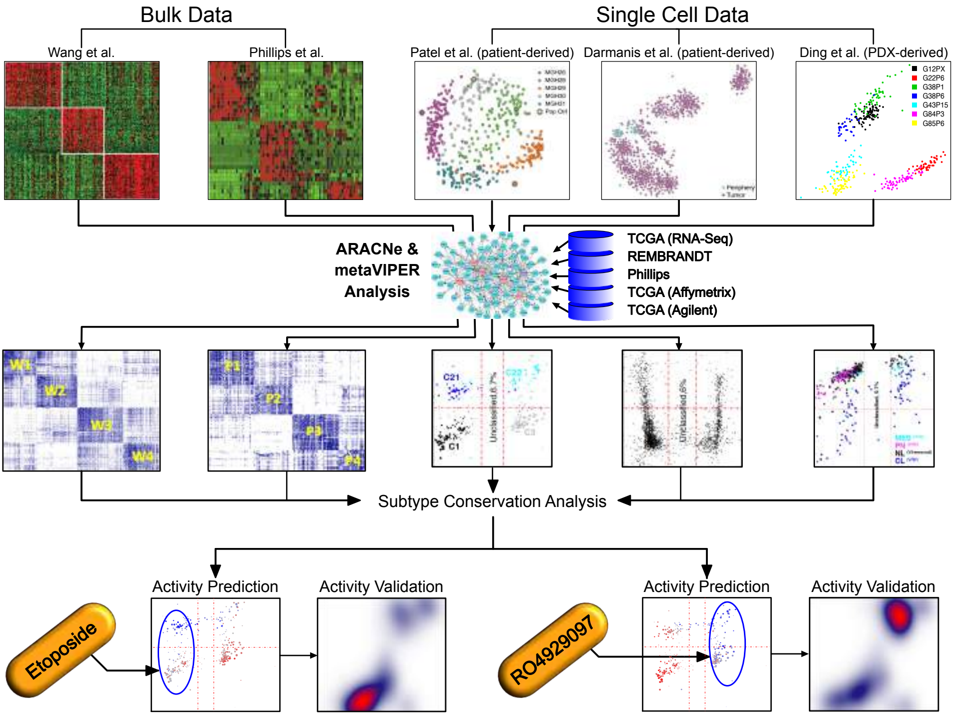

#### Fig2SupFig A: Silhouette score protein activity Phillips

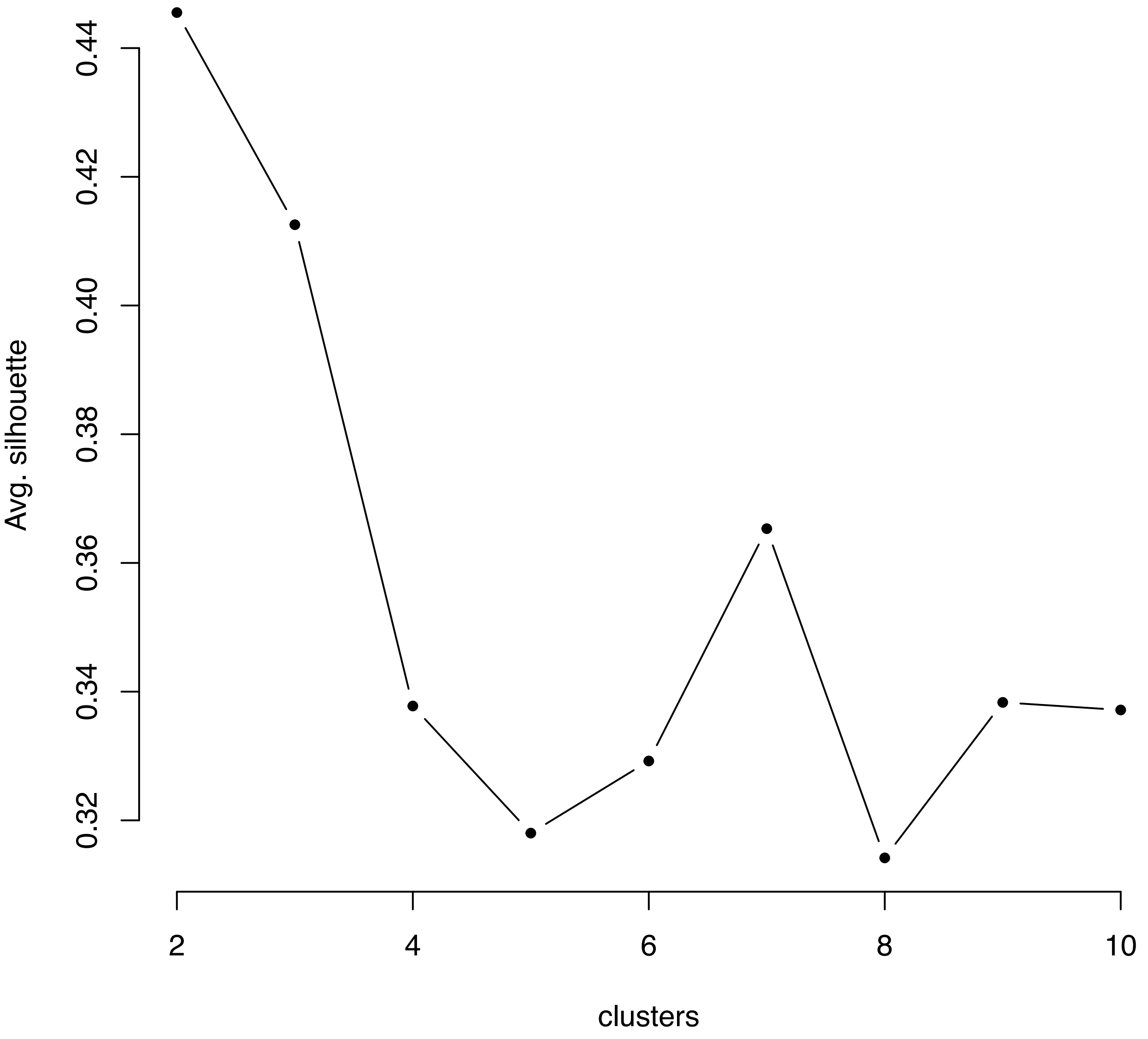

#### Fig2SupFig B: Silhouette score protein activity Wang

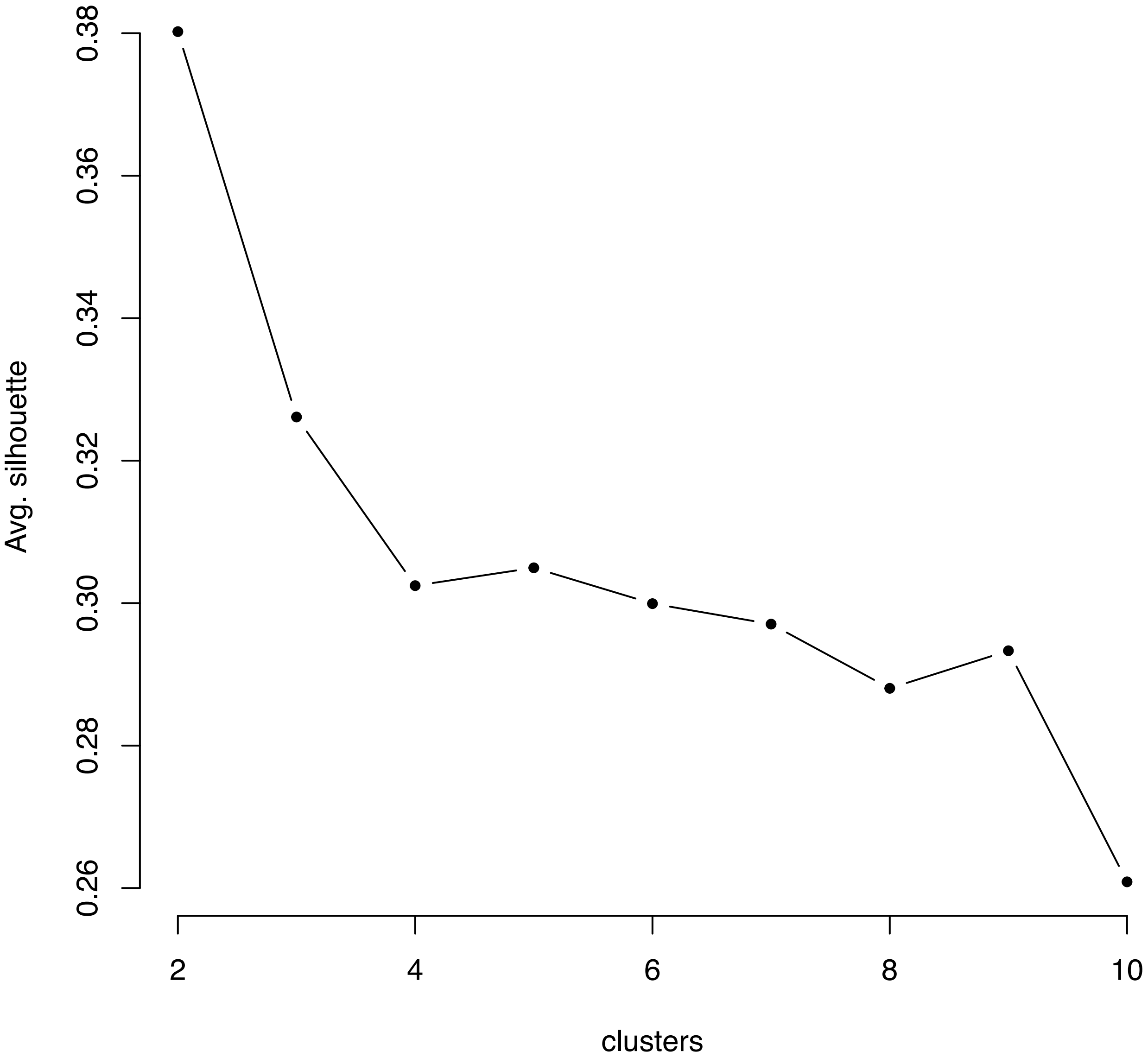

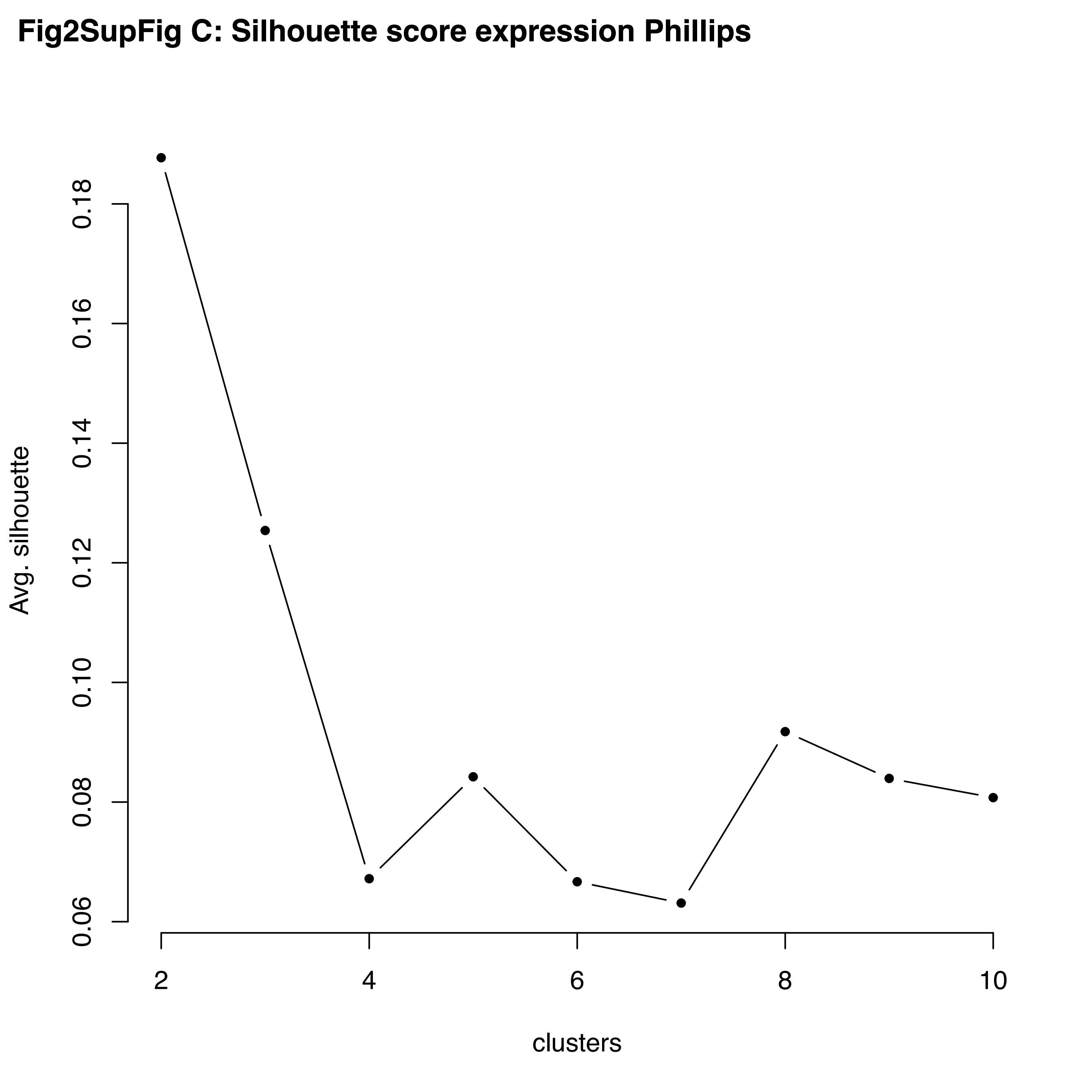
Fig2SupFig C: Silhouette score expression Phillips
Avg. silhouette
0.06
0.08
0.10
0.12
0.14
0.16
0.18
2
4
6
8
10
clusters

**Fig2SupFig D: Silhouette score expression Wang**

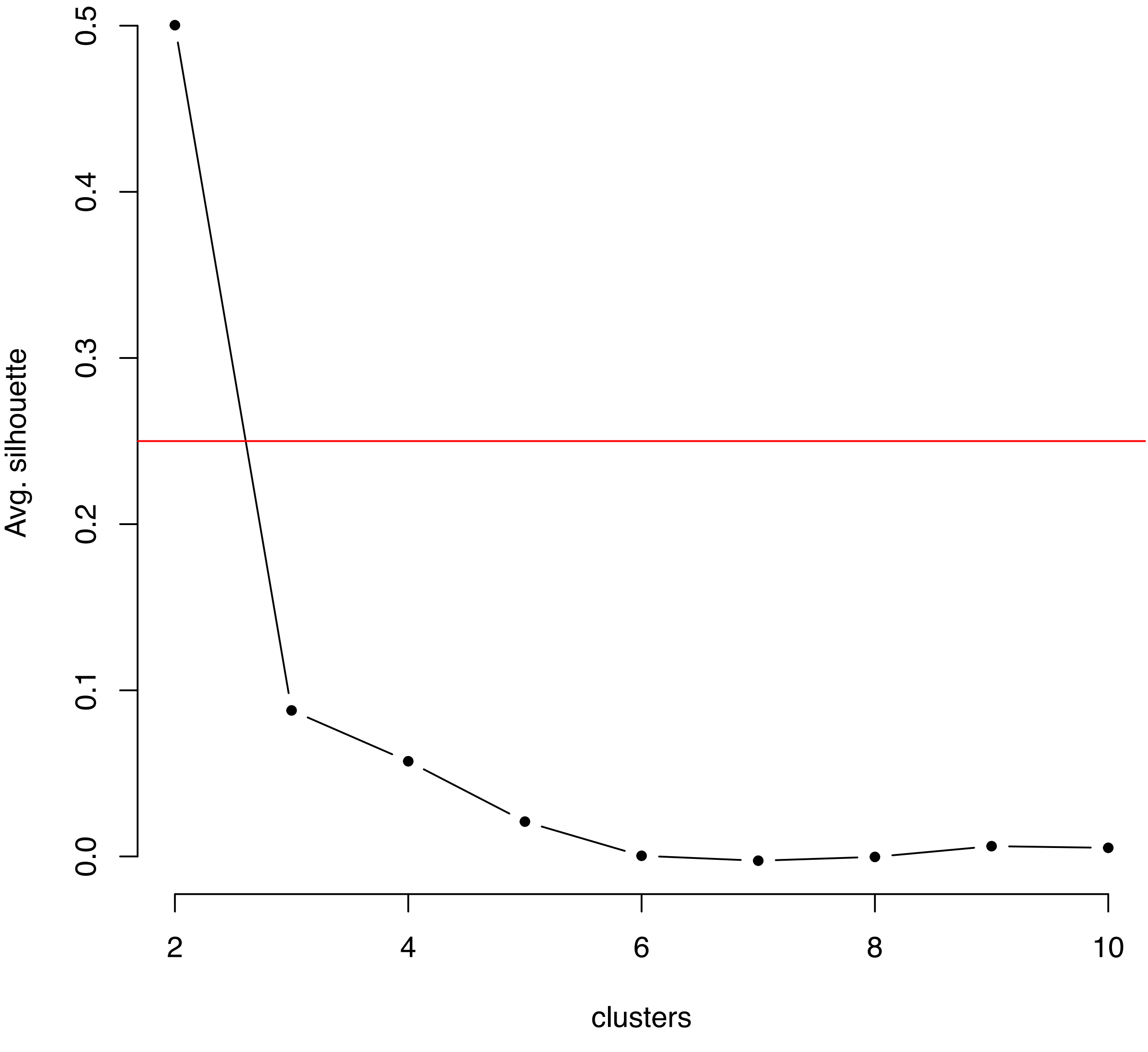

Fig2SupFig E: Phillips Dataset: Protein Activity based clustering, K = 2)

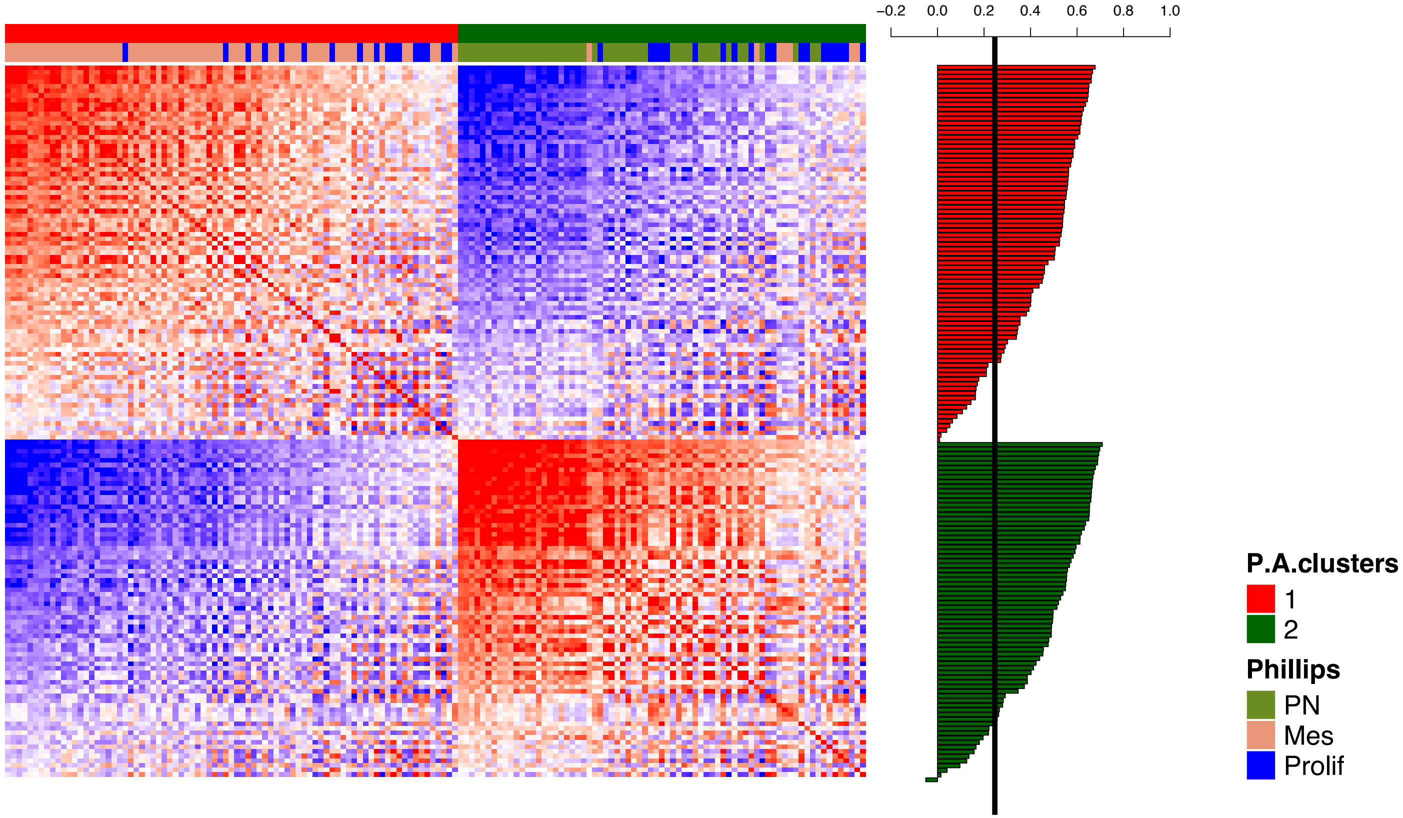

### Fig2SupFig F: Wang Dataset: Protein Activity based clustering, K = 2

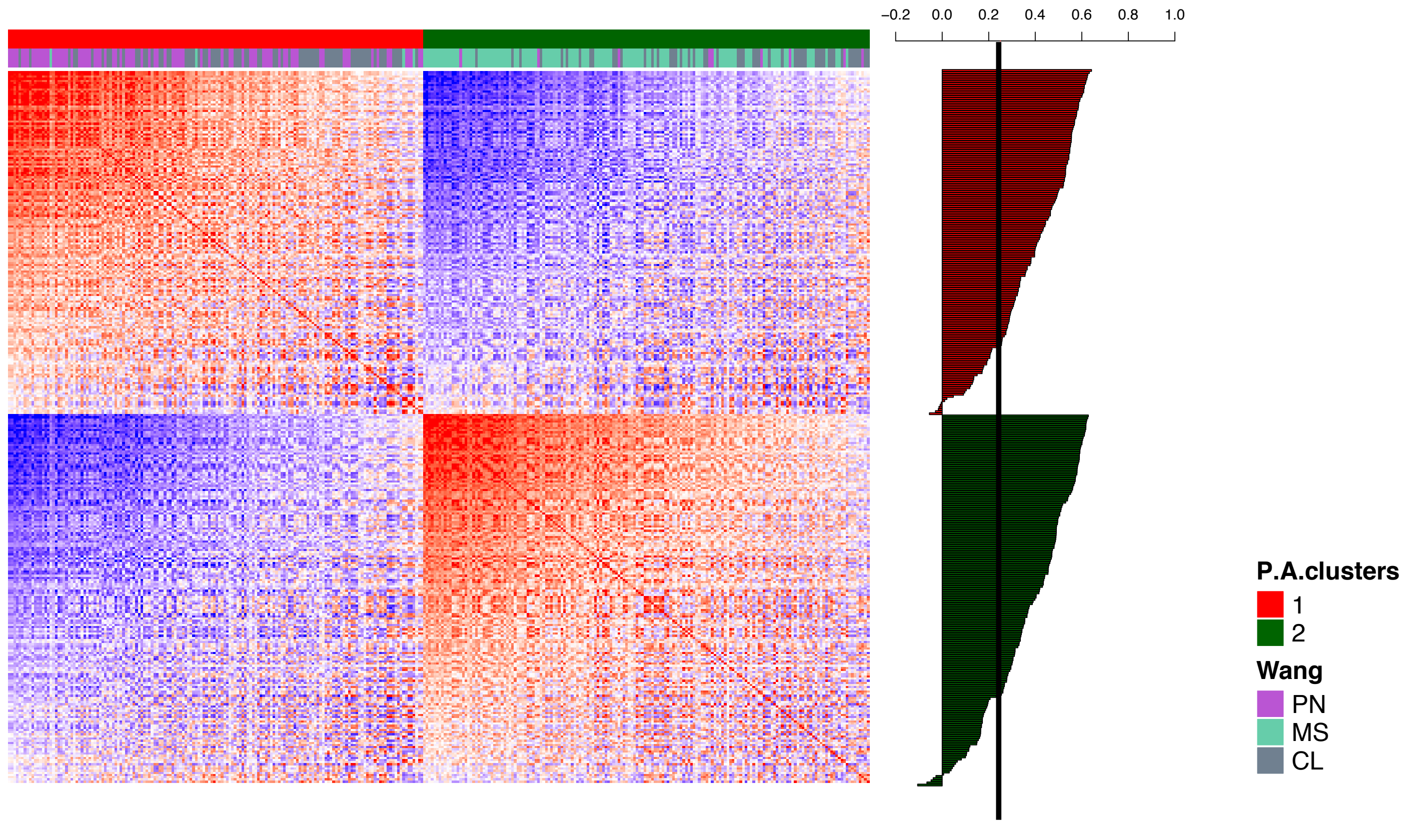

Fig2SupFig G: Phillips Dataset: Gene expression based clustering, K = 2

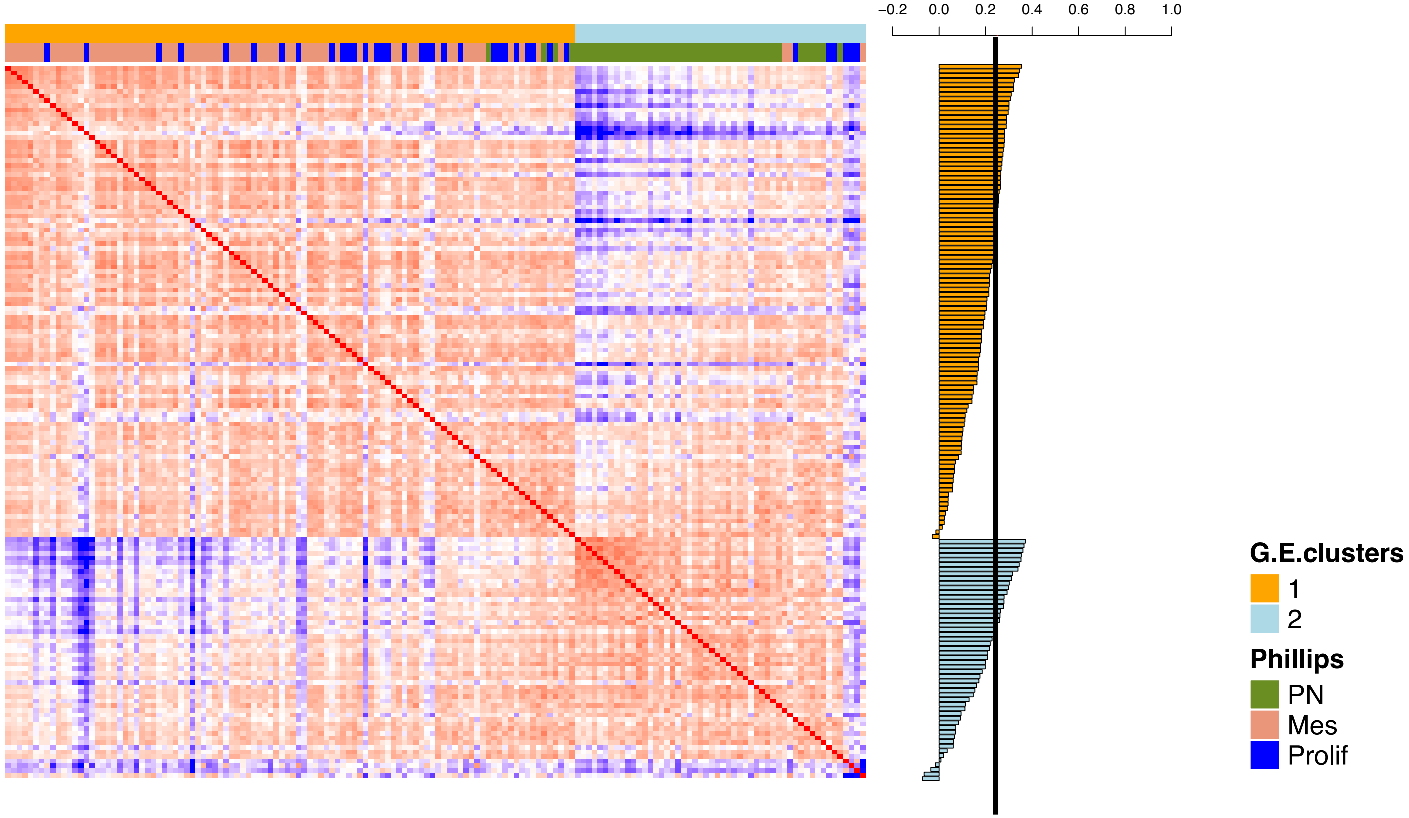

### Fig2SupFig H: Wang Dataset: Gene expression based clustering, K = 2

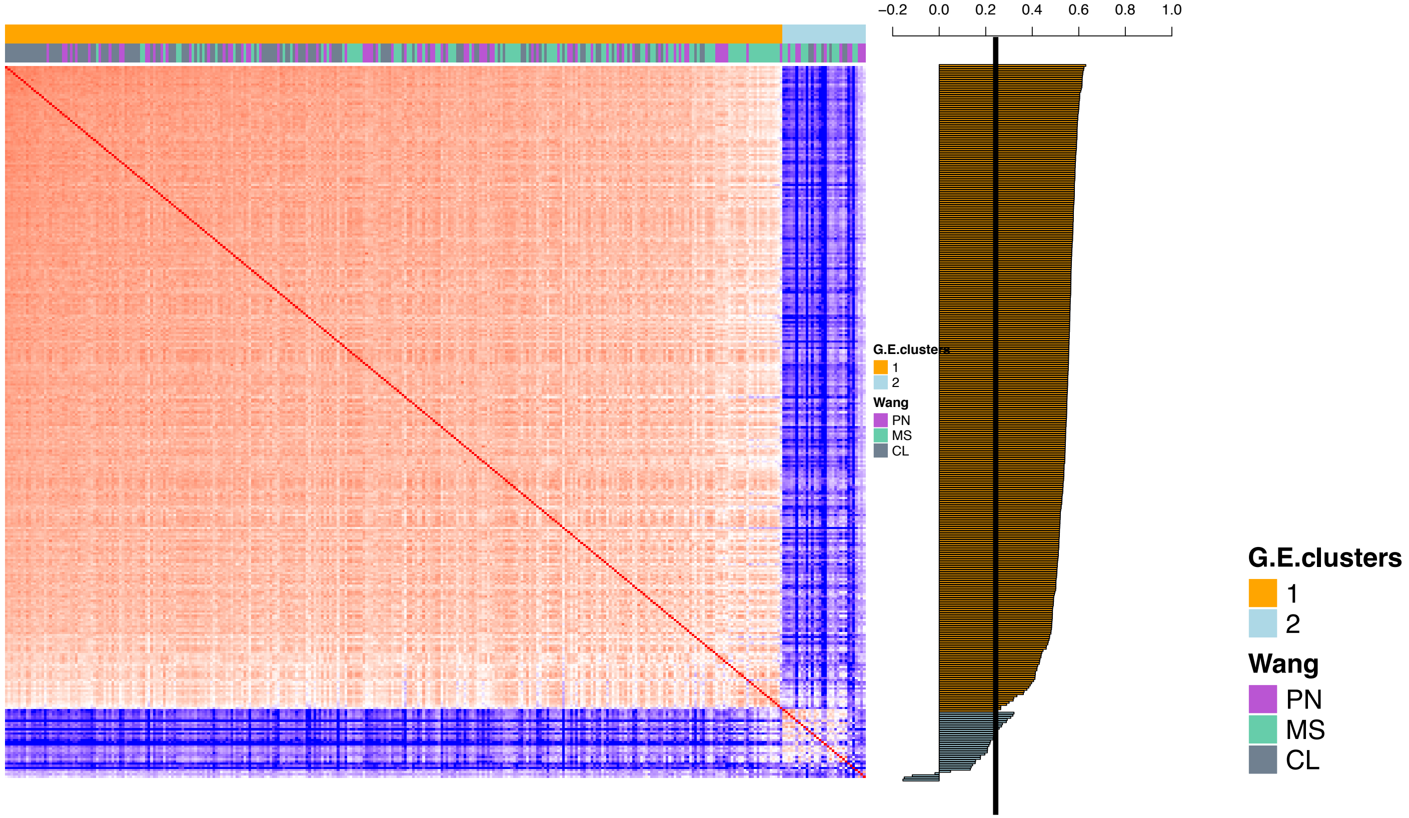

**Fig2SupFig I**

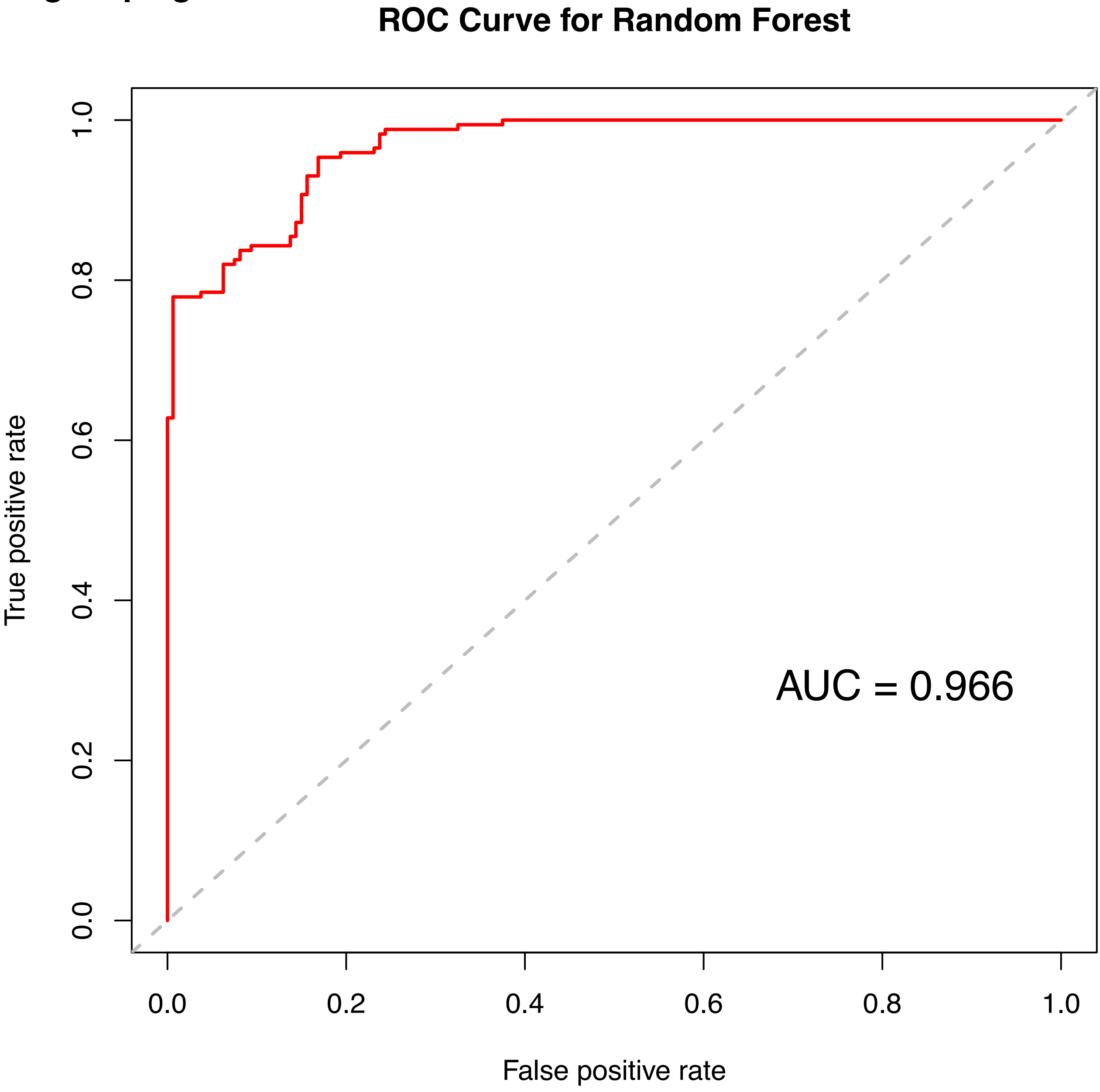

**Fig2SupFig J**

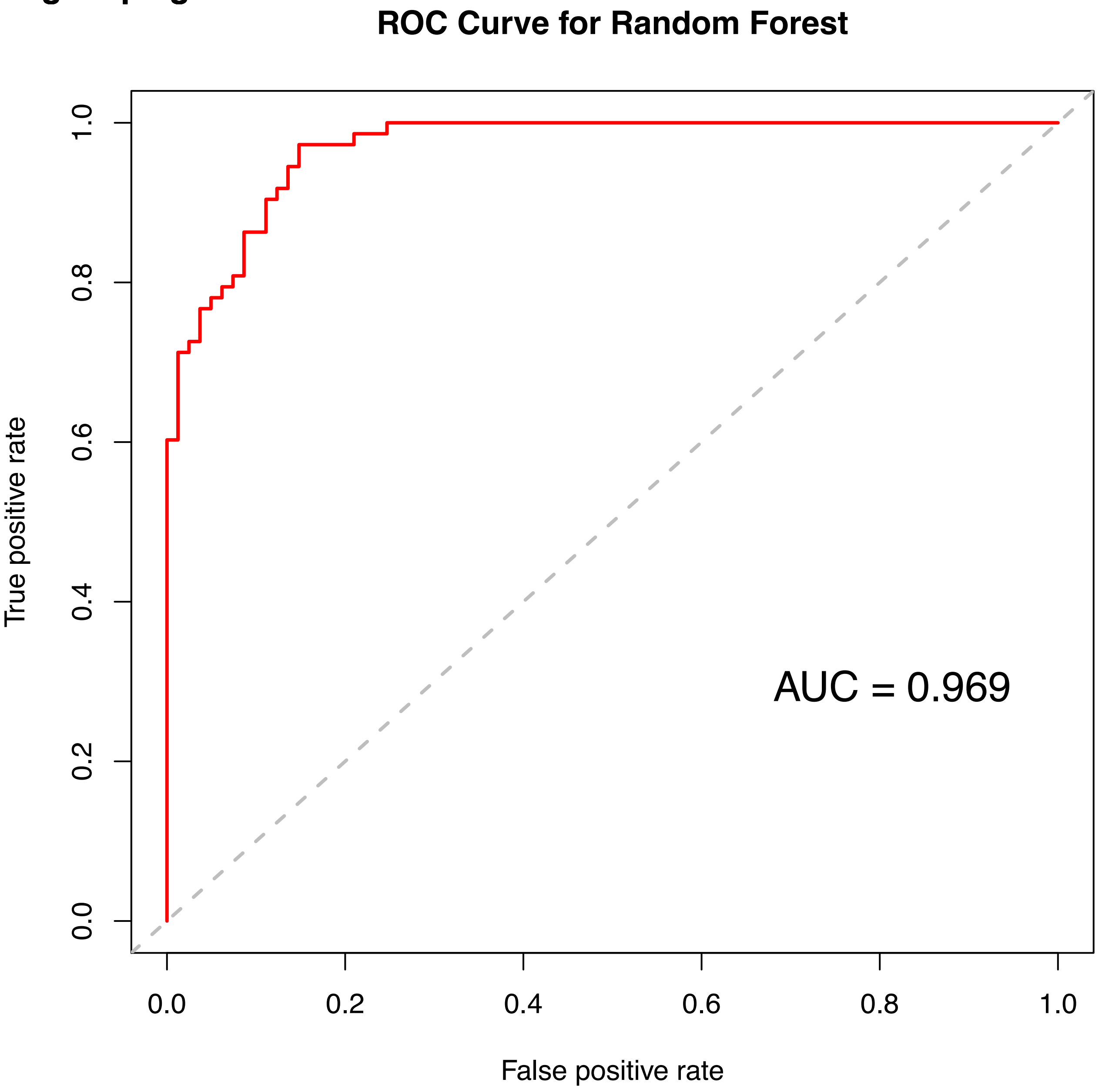

Fig2SupFig K: Differentially activated proteins: Phillips (K = 2)

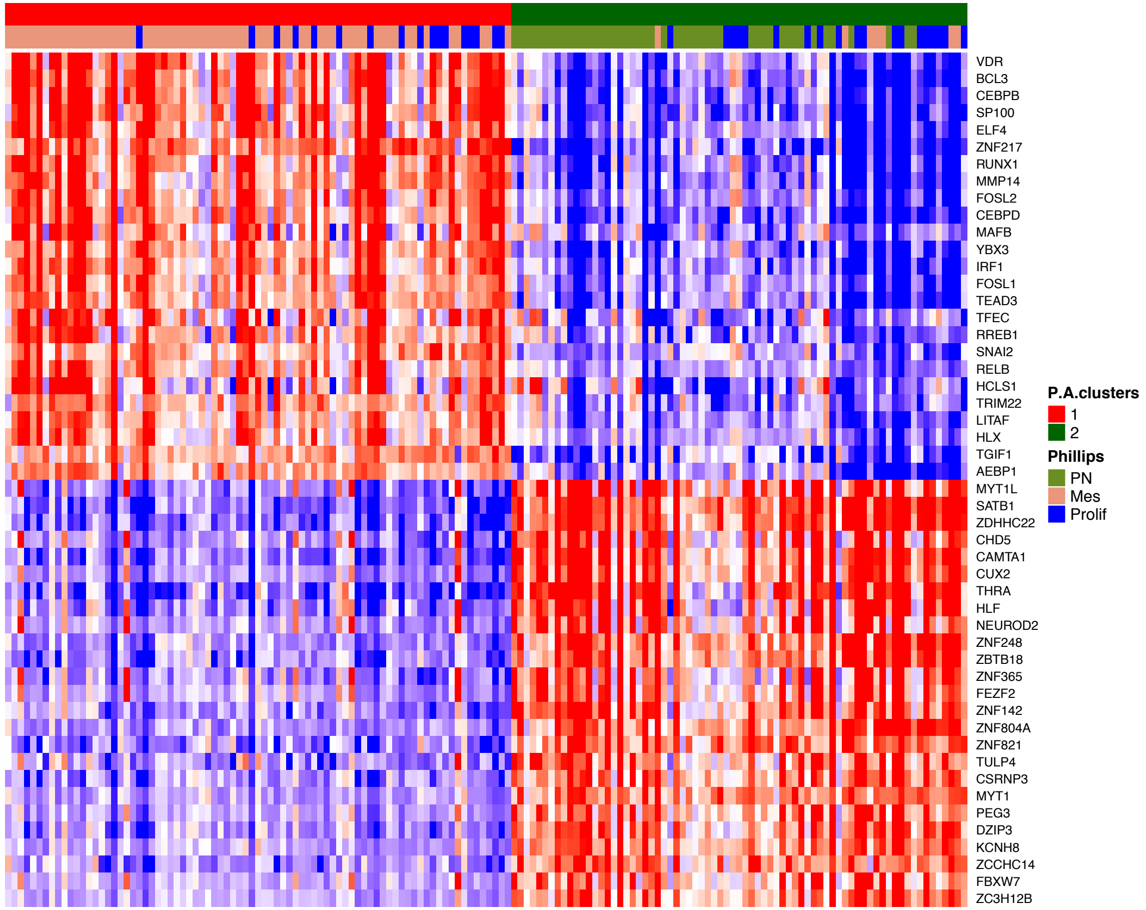


Fig2SupFig L: Differentially activated proteins: Wang (K = 2)

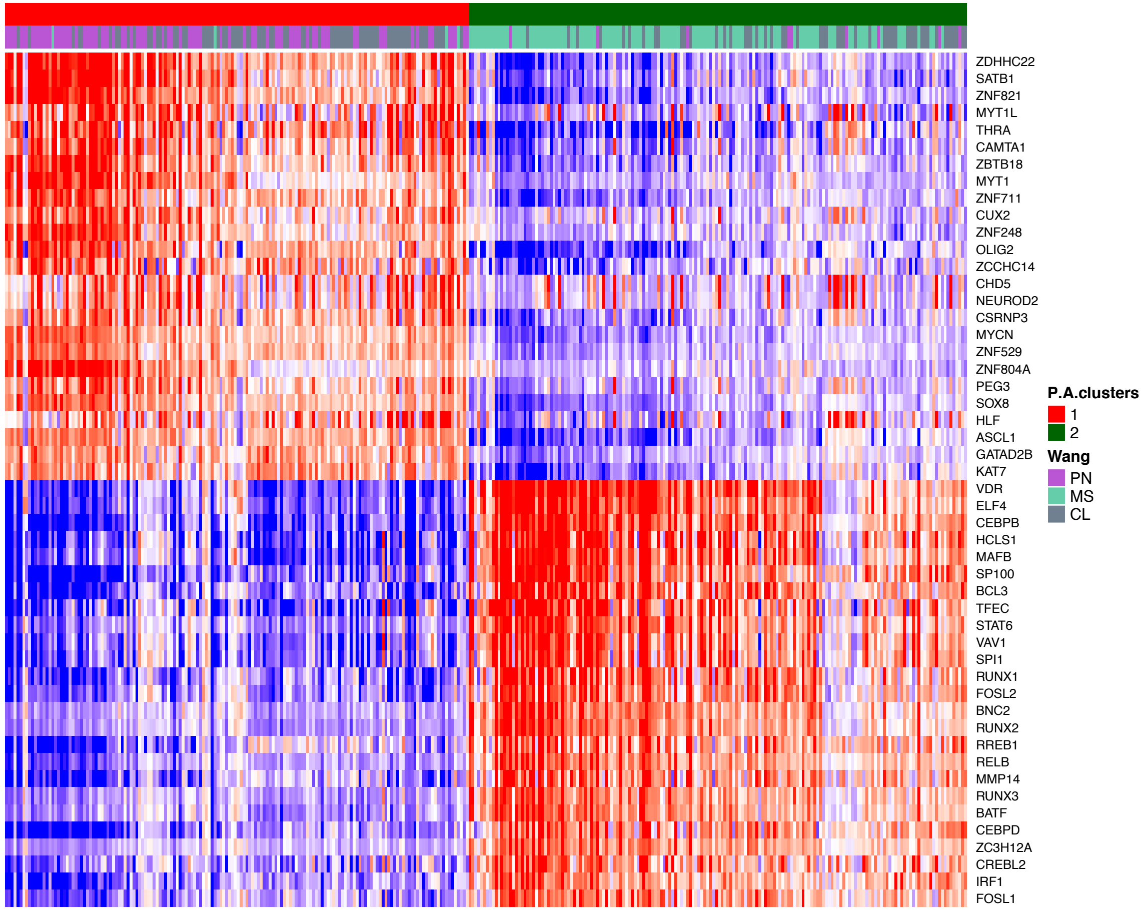

### Fig2SupFig M: GBM analysis  (Phillips "iterative clustering" analysis)

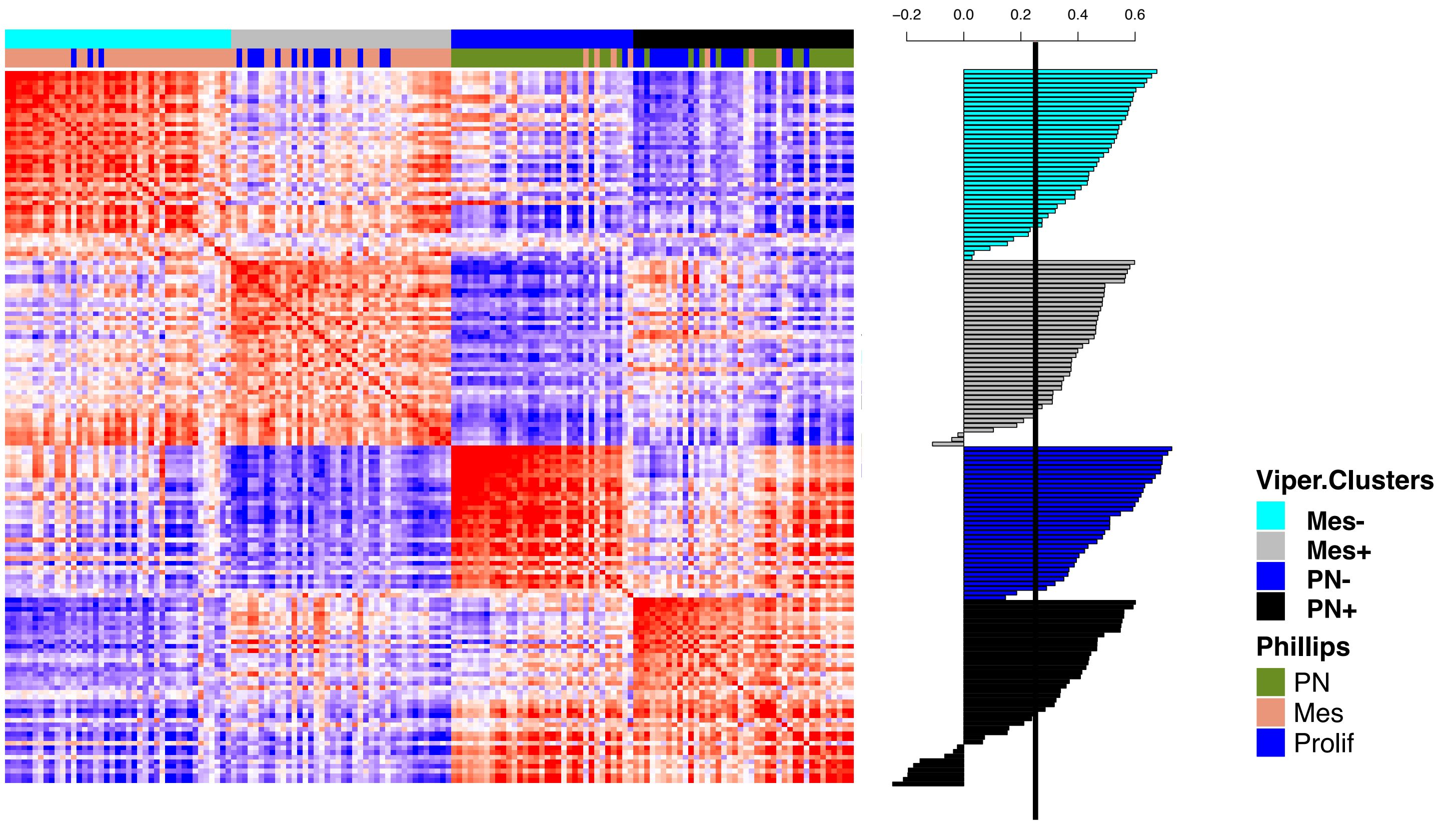

Fig2SupFig N: GBM analysis  (Wang “iterative clustering” analysis)

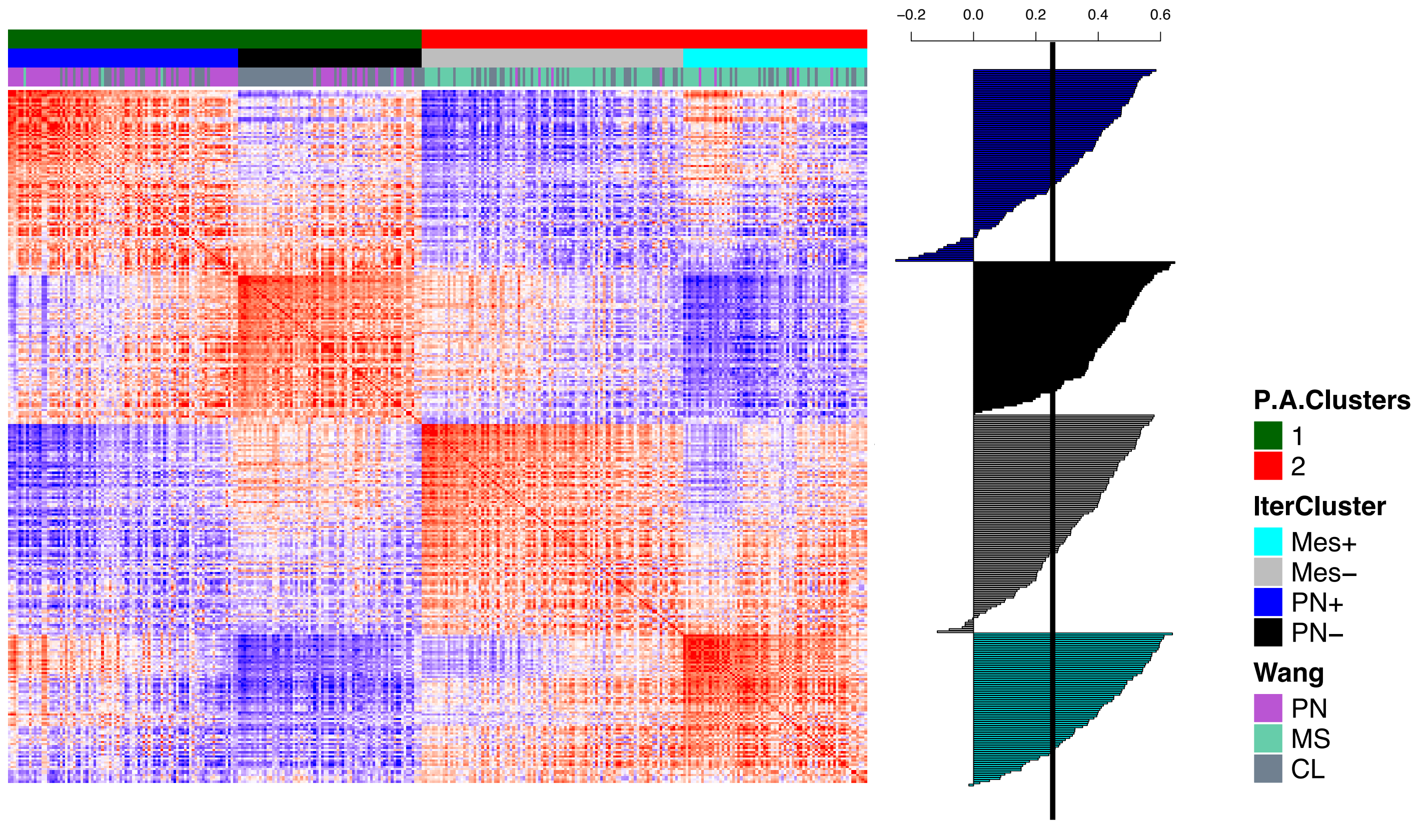

### Fig3SupFig

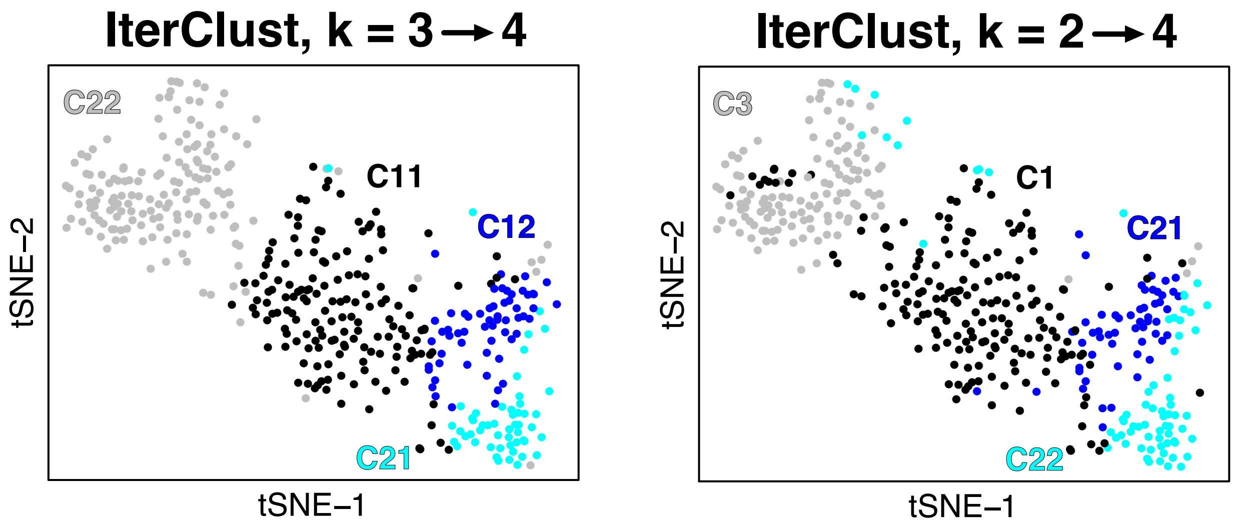

IterClust, k = 3 → 4
C22
C11
C12
C21
tSNE−2
tSNE−1
IterClust, k = 2 → 4
C3
C1
C21
C22
tSNE−2
tSNE−1

### Fig4SupFig-1

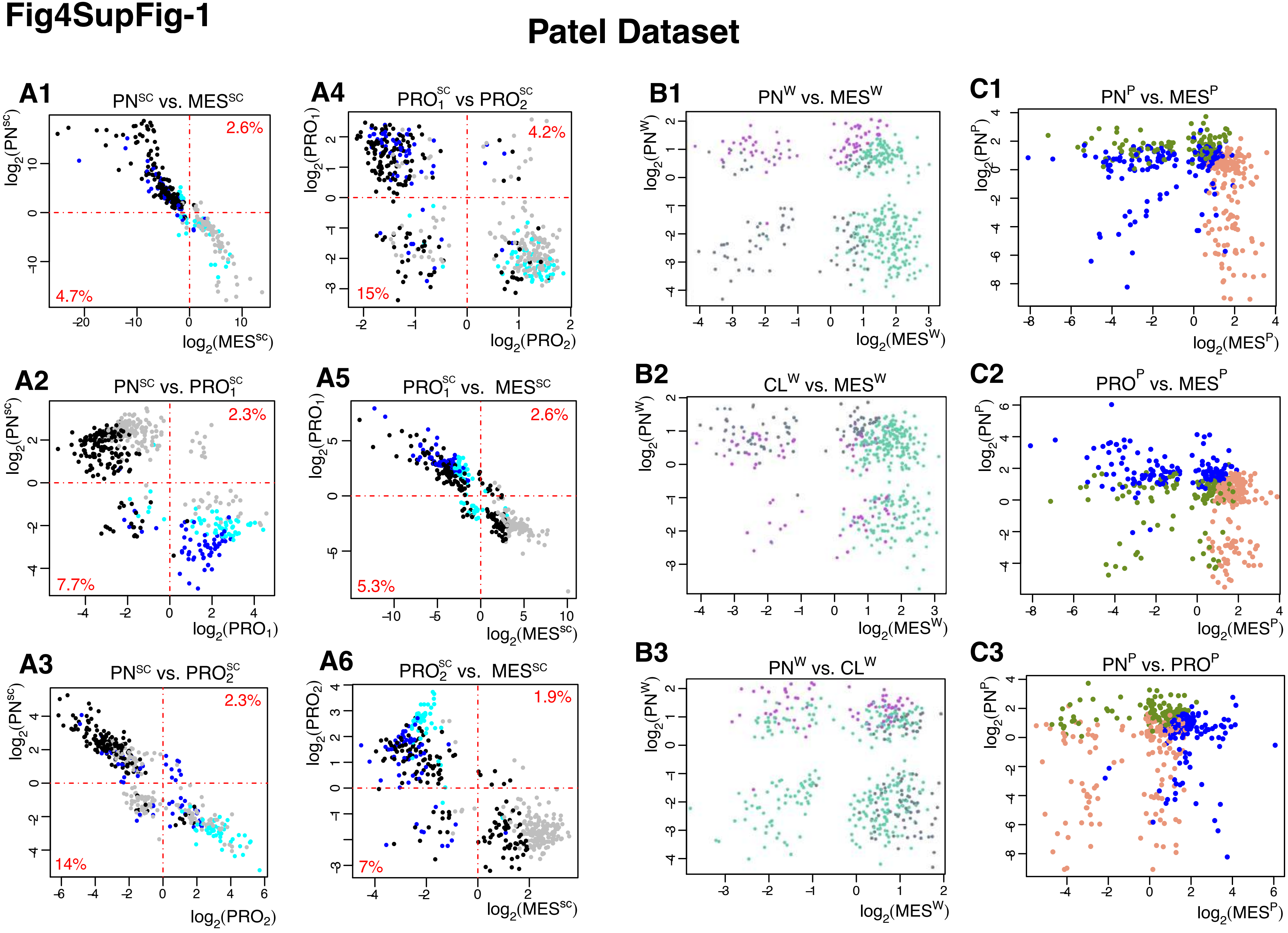

**Fig4upFig-2**

### Darmanis Dataset

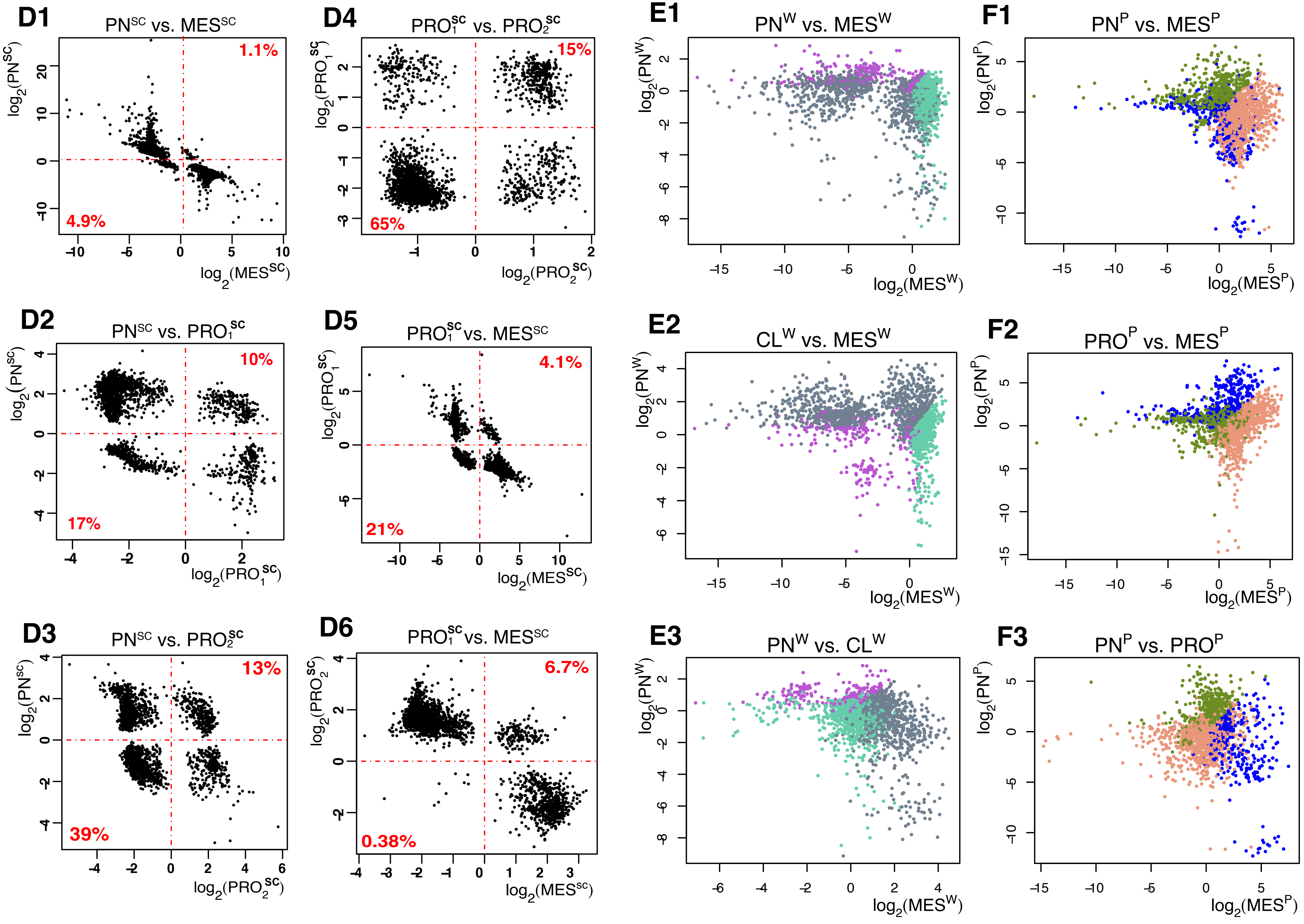

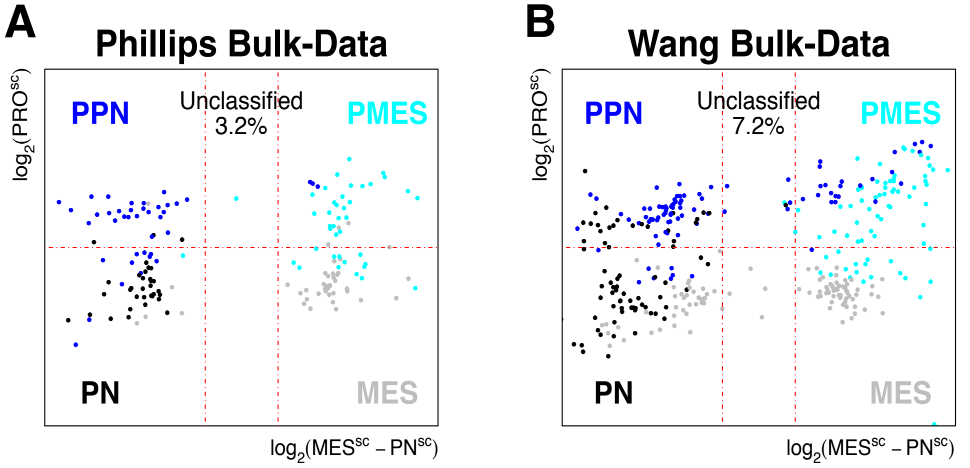


**Fig4SupFig-3**

### Fig5SupFig-1

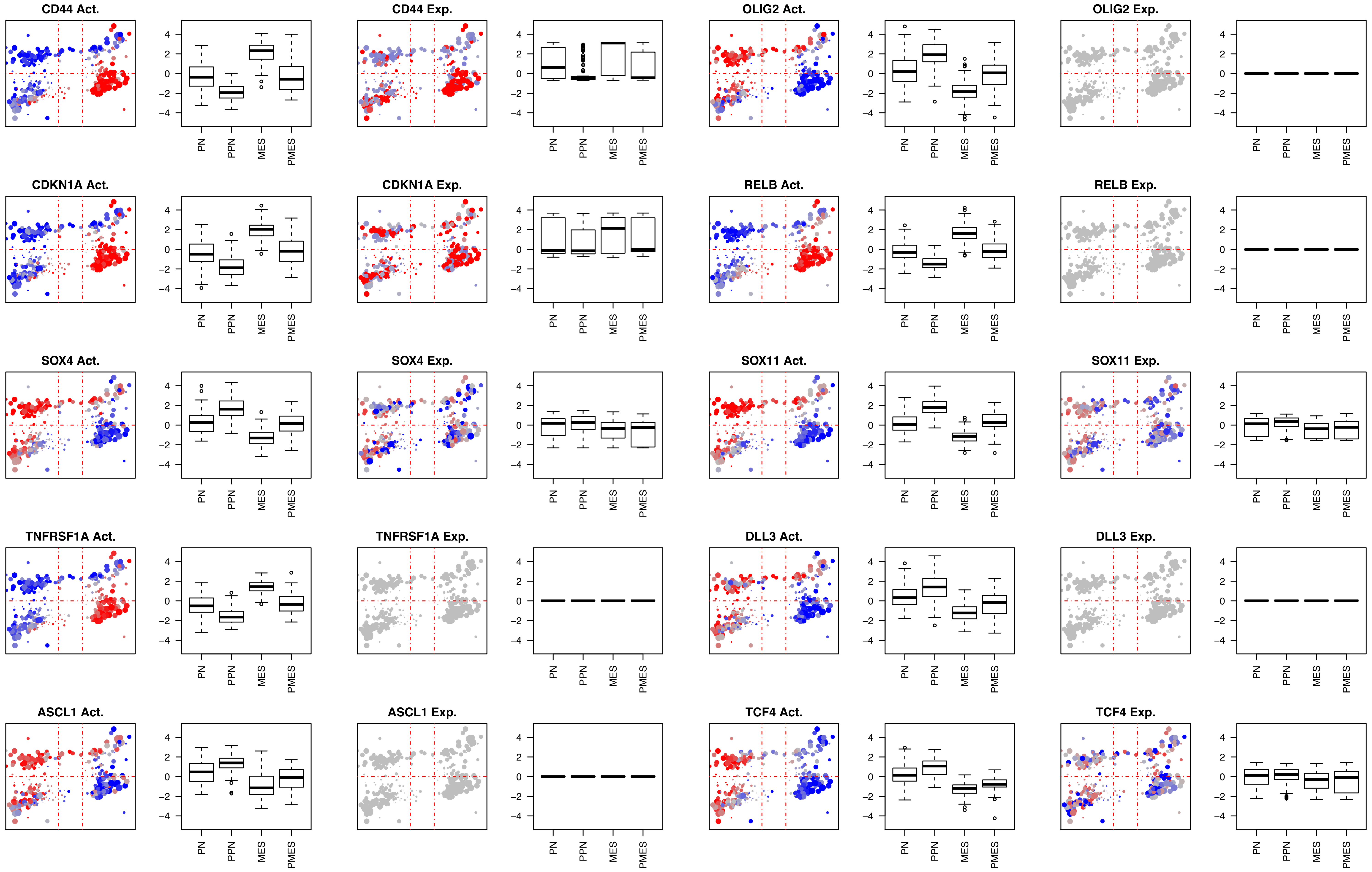

CD44 Act.
CD44 Exp.
OLIG2 Act.
OLIG2 Exp.
CDKN1A Act.
CDKN1A Exp.
RELB Act.
RELB Exp.
SOX4 Act.
SOX4 Exp.
SOX11 Act.
SOX11 Exp.
TNFRSF1A Act.
TNFRSF1A Exp.
DLL3 Act.
DLL3 Exp.
ASCL1 Act.
ASCL1 Exp.
TCF4 Act.
TCF4 Exp.
4
2
0
-2
-4
PN
PPN
MES
PMES

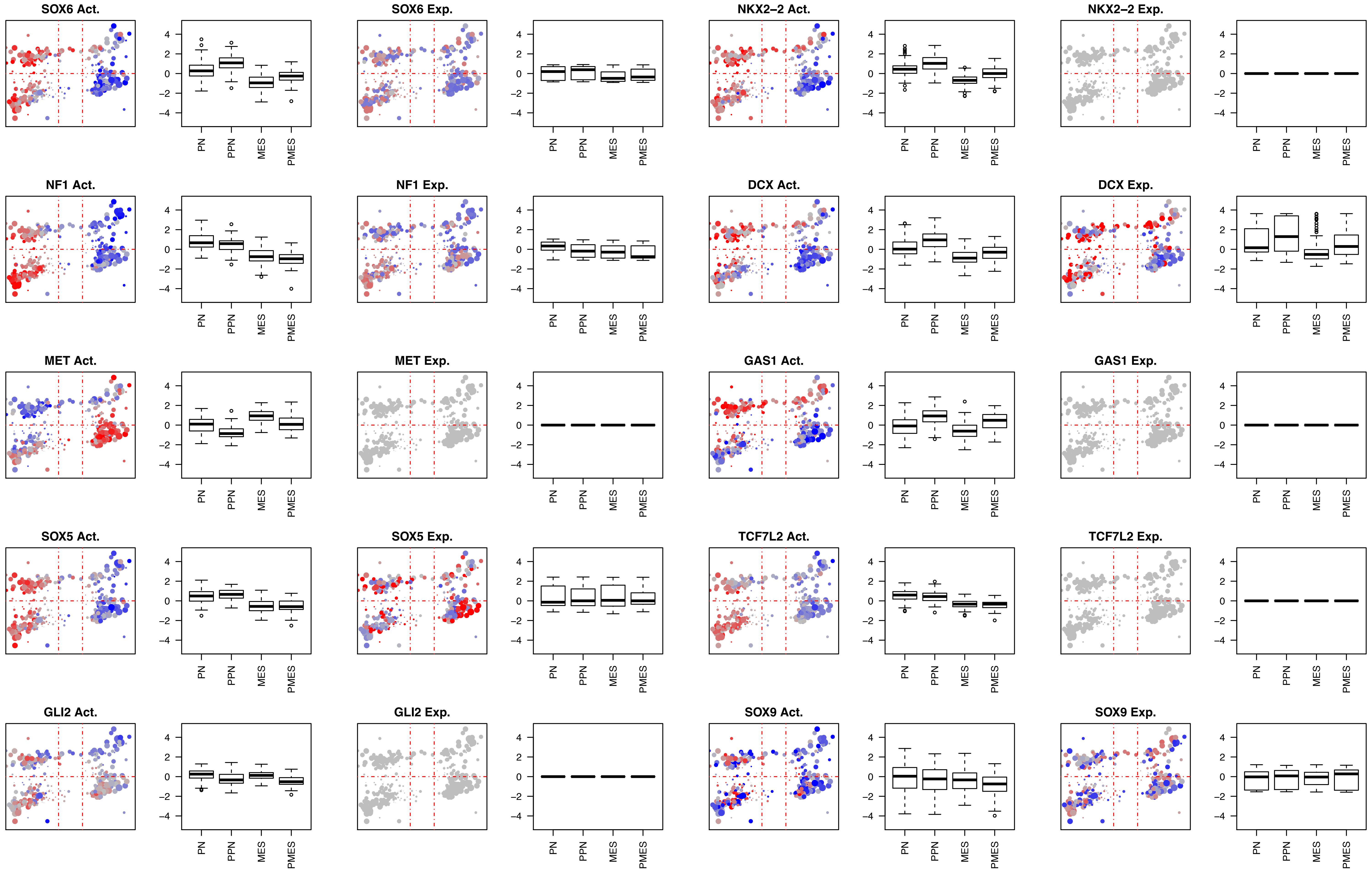
SOX6 Act.
SOX6 Exp.
NKX2–2 Act.
NKX2–2 Exp.
NF1 Act.
NF1 Exp.
DCX Act.
DCX Exp.
MET Act.
MET Exp.
GAS1 Act.
GAS1 Exp.
SOX5 Act.
SOX5 Exp.
TCF7L2 Act.
TCF7L2 Exp.
GLI2 Act.
GLI2 Exp.
SOX9 Act.
SOX9 Exp.
4
2
0
−2
−4
PN
PPN
MES
PMES

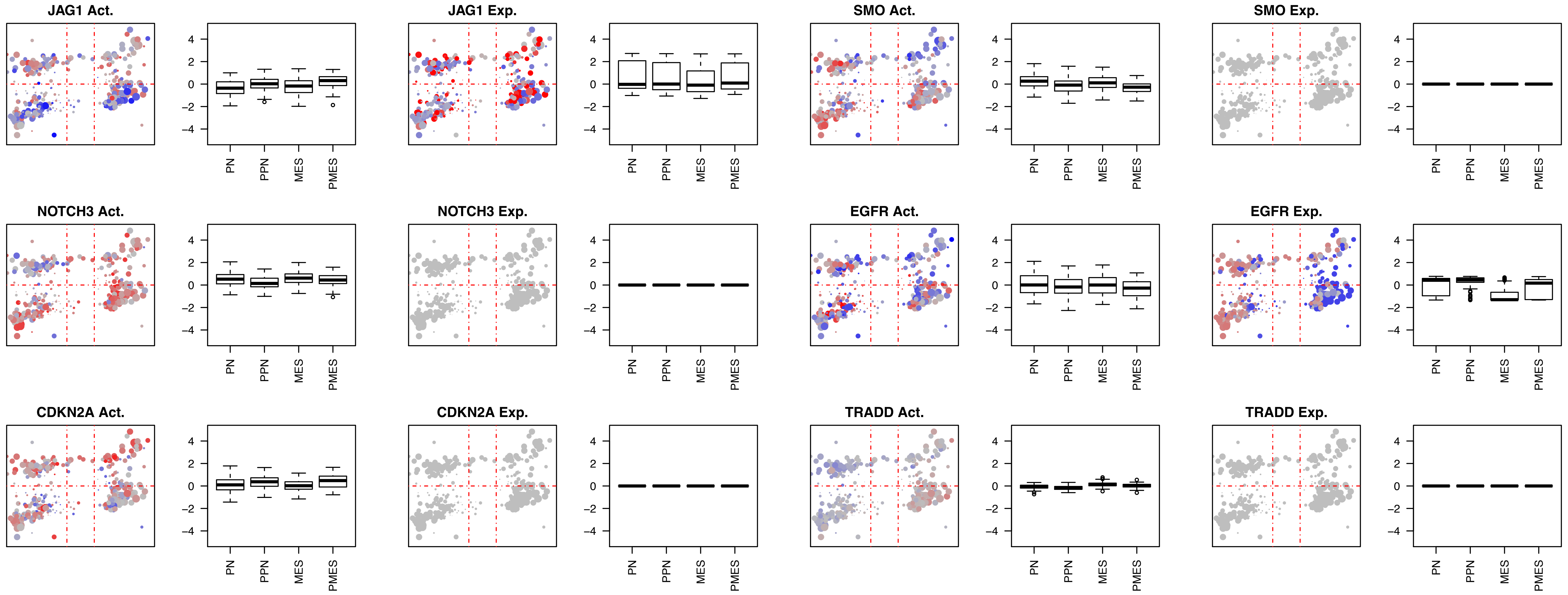

JAG1 Act.
JAG1 Exp.
SMO Act.
SMO Exp.
NOTCH3 Act.
NOTCH3 Exp.
EGFR Act.
EGFR Exp.
CDKN2A Act.
CDKN2A Exp.
TRADD Act.
TRADD Exp.
4
2
0
−2
−4
PN
PPN
MES
PMES

Fig5SupFig-2

**Proteins aberrantly activated in the PN quadrant.**

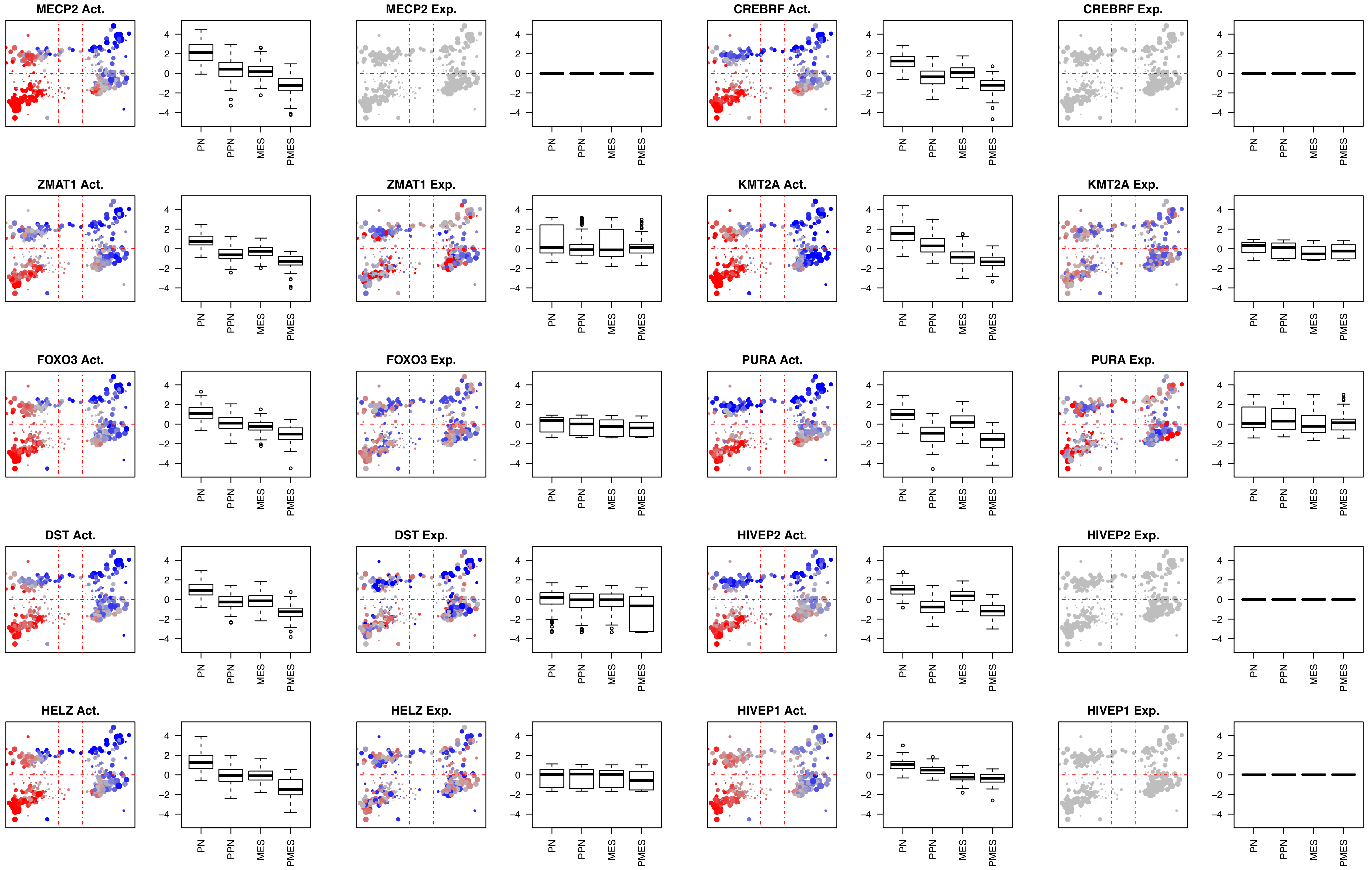
MECP2 Act.
MECP2 Exp.
CREBRF Act.
CREBRF Exp.
ZMAT1 Act.
ZMAT1 Exp.
KMT2A Act.
KMT2A Exp.
FOXO3 Act.
FOXO3 Exp.
PURA Act.
PURA Exp.
DST Act.
DST Exp.
HIVEP2 Act.
HIVEP2 Exp.
HELZ Act.
HELZ Exp.
HIVEP1 Act.
HIVEP1 Exp.
4
2
0
−2
−4
PN
PPN
MES
PMES

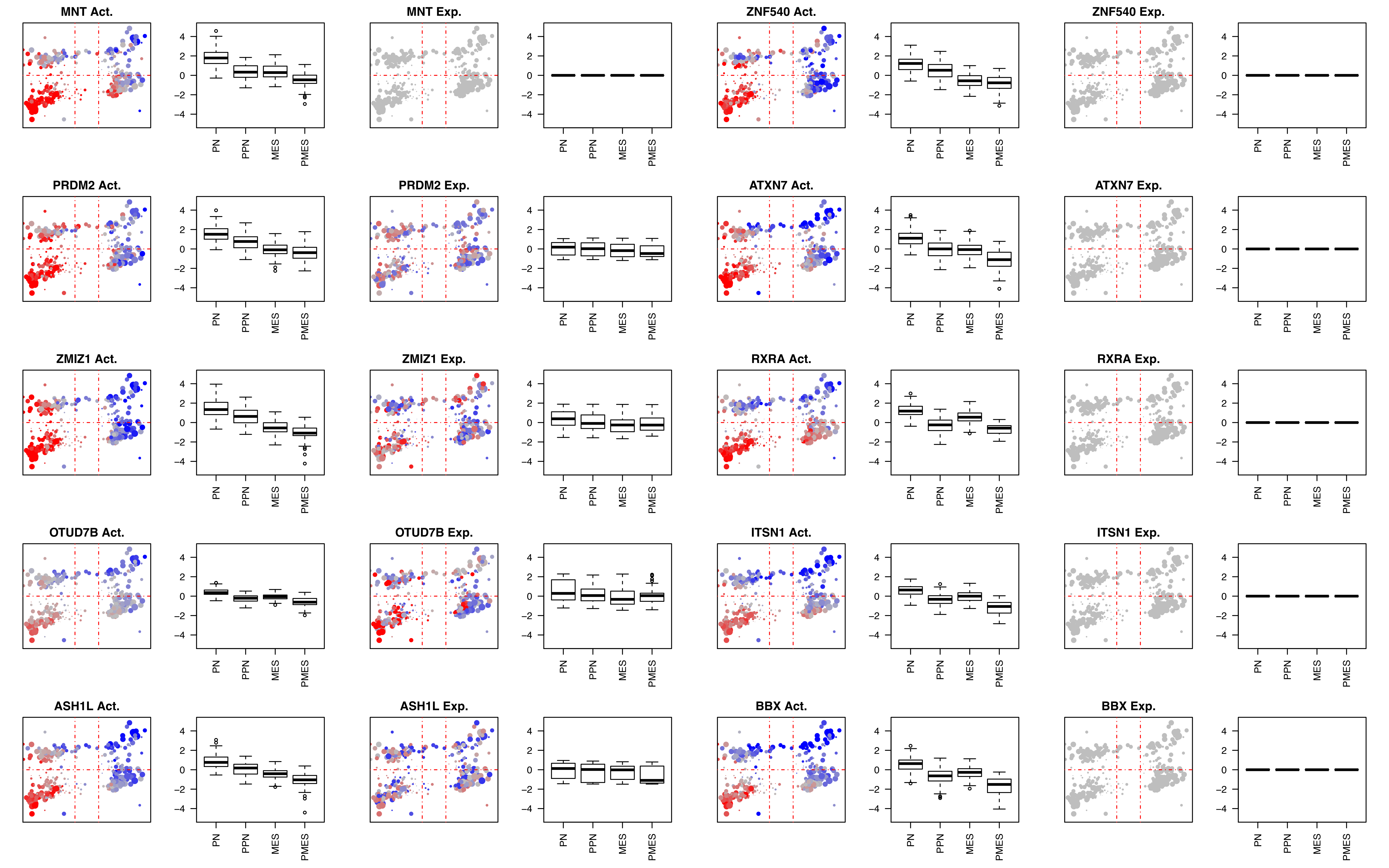

MNT Act.
MNT Exp.
ZNF540 Act.
ZNF540 Exp.
PRDM2 Act.
PRDM2 Exp.
ATXN7 Act.
ATXN7 Exp.
ZMIZ1 Act.
ZMIZ1 Exp.
RXRA Act.
RXRA Exp.
OTUD7B Act.
OTUD7B Exp.
ITSN1 Act.
ITSN1 Exp.
ASH1L Act.
ASH1L Exp.
BBX Act.
BBX Exp.
4
2
0
-2
-4
PN
PPN
MES
PMES

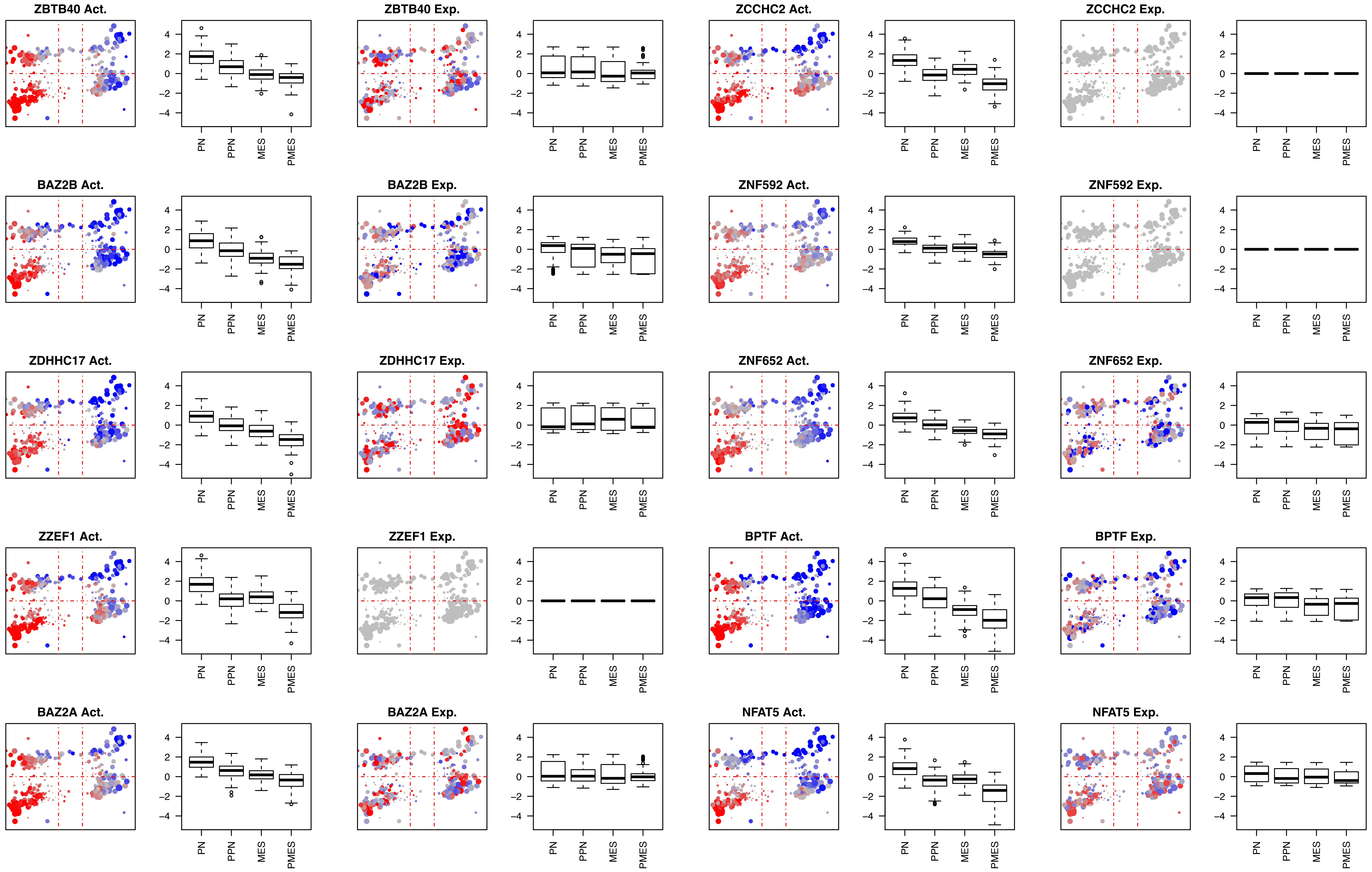

ZBTB40 Act.
ZBTB40 Exp.
ZCCHC2 Act.
ZCCHC2 Exp.
BAZ2B Act.
BAZ2B Exp.
ZNF592 Act.
ZNF592 Exp.
ZDHHC17 Act.
ZDHHC17 Exp.
ZNF652 Act.
ZNF652 Exp.
ZZEF1 Act.
ZZEF1 Exp.
BPTF Act.
BPTF Exp.
BAZ2A Act.
BAZ2A Exp.
NFAT5 Act.
NFAT5 Exp.
4
2
0
−2
−4
PN
PPN
MES
PMES

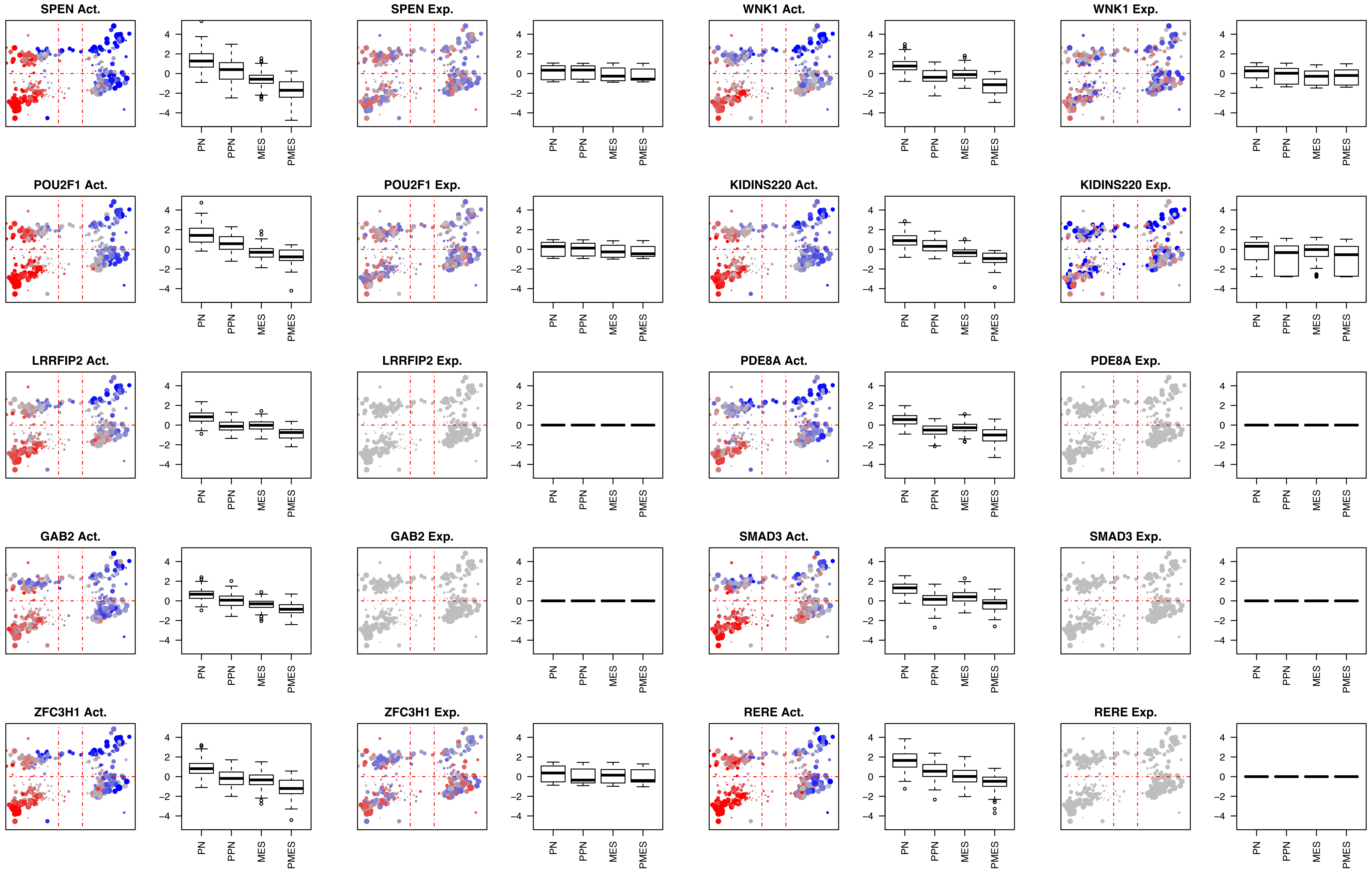
SPEN Act.
SPEN Exp.
WNK1 Act.
WNK1 Exp.
POU2F1 Act.
POU2F1 Exp.
KIDINS220 Act.
KIDINS220 Exp.
LRRFIP2 Act.
LRRFIP2 Exp.
PDE8A Act.
PDE8A Exp.
GAB2 Act.
GAB2 Exp.
SMAD3 Act.
SMAD3 Exp.
ZFC3H1 Act.
ZFC3H1 Exp.
RERE Act.
RERE Exp.
4
2
0
−2
−4
PN
PPN
MES
PMES

**Proteins aberrantly activated in the PPN quadrant.**

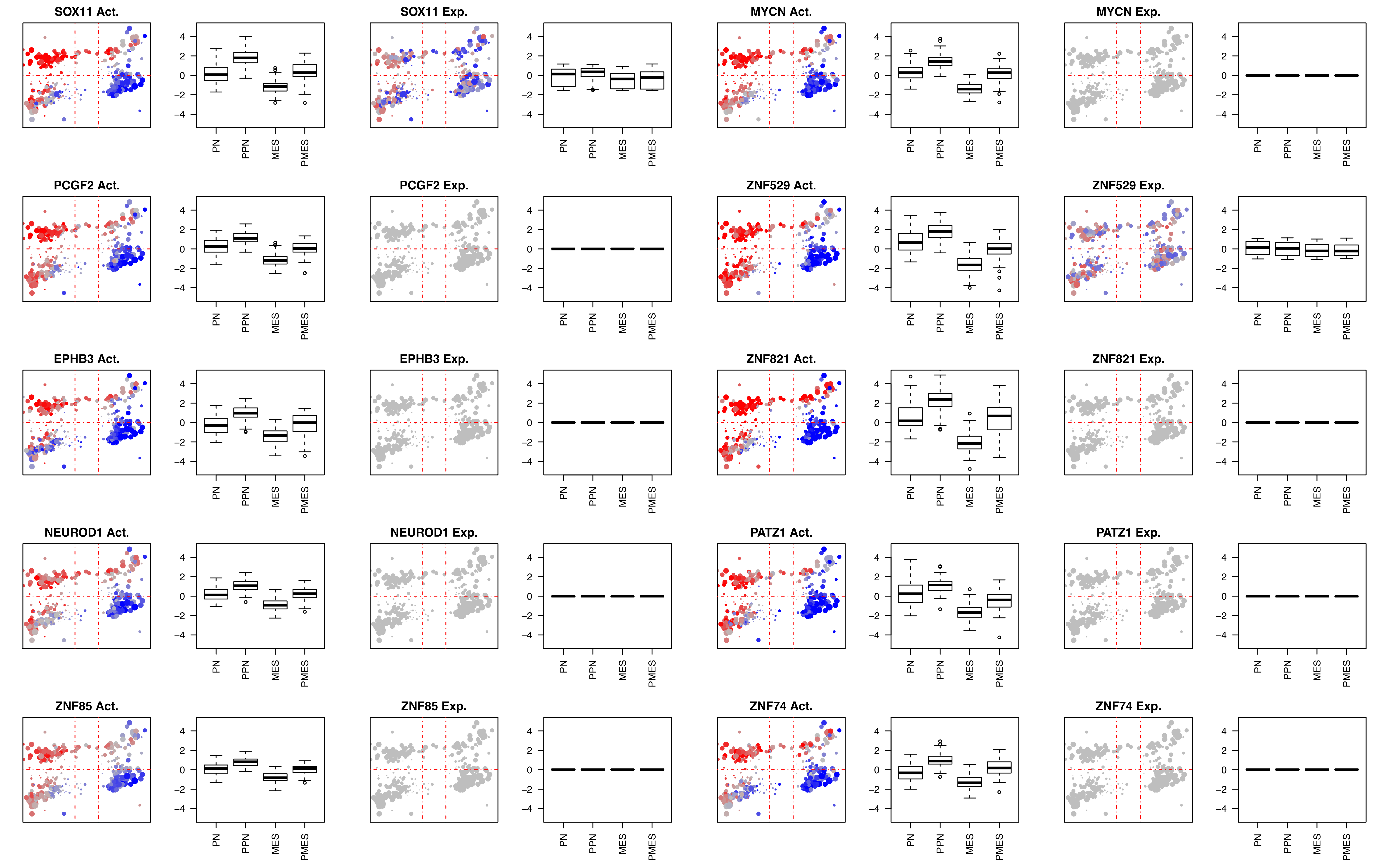
SOX11 Act.
SOX11 Exp.
MYCN Act.
MYCN Exp.
PCGF2 Act.
PCGF2 Exp.
ZNF529 Act.
ZNF529 Exp.
EPHB3 Act.
EPHB3 Exp.
ZNF821 Act.
ZNF821 Exp.
NEUROD1 Act.
NEUROD1 Exp.
PATZ1 Act.
PATZ1 Exp.
ZNF85 Act.
ZNF85 Exp.
ZNF74 Act.
ZNF74 Exp.
4
2
0
−2
−4
PN
PPN
MES
PMES

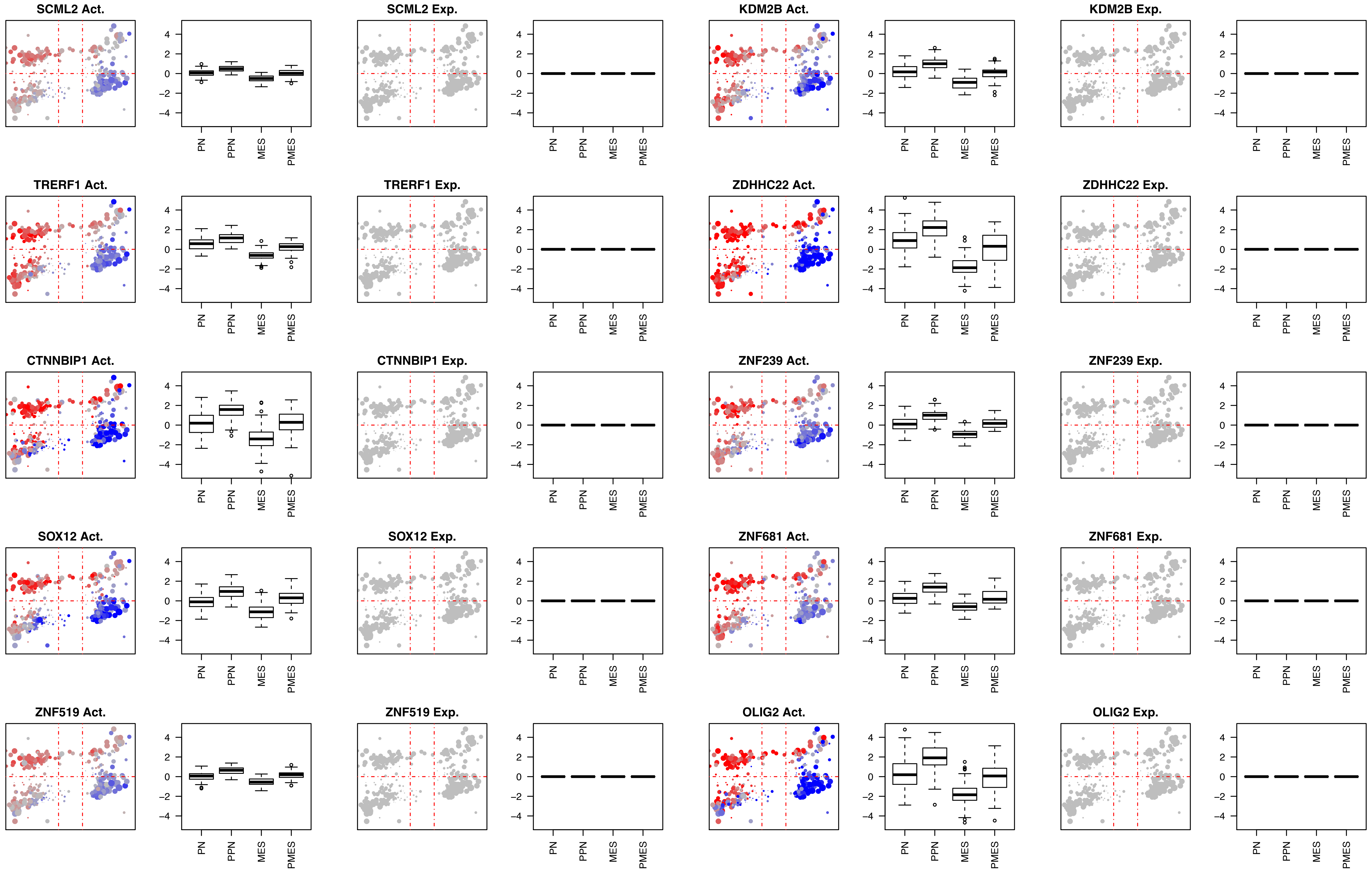
SCML2 Act.
SCML2 Exp.
KDM2B Act.
KDM2B Exp.
TRERF1 Act.
TRERF1 Exp.
ZDHHC22 Act.
ZDHHC22 Exp.
CTNNBIP1 Act.
CTNNBIP1 Exp.
ZNF239 Act.
ZNF239 Exp.
SOX12 Act.
SOX12 Exp.
ZNF681 Act.
ZNF681 Exp.
ZNF519 Act.
ZNF519 Exp.
OLIG2 Act.
OLIG2 Exp.
4
2
0
−2
−4
PN
PPN
MES
PMES

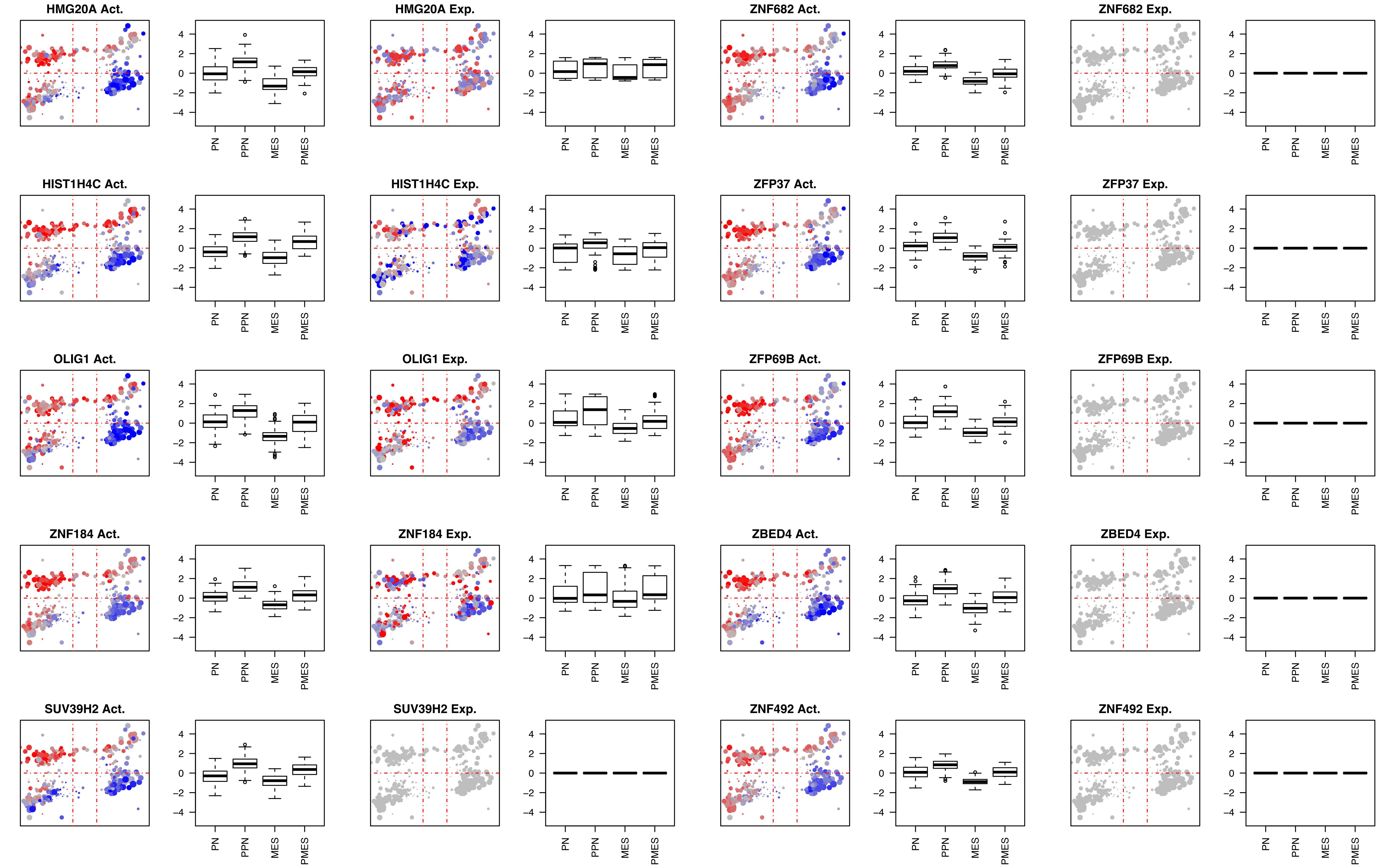
HMG20A Act.
HMG20A Exp.
ZNF682 Act.
ZNF682 Exp.
HIST1H4C Act.
HIST1H4C Exp.
ZFP37 Act.
ZFP37 Exp.
OLIG1 Act.
OLIG1 Exp.
ZFP69B Act.
ZFP69B Exp.
ZNF184 Act.
ZNF184 Exp.
ZBED4 Act.
ZBED4 Exp.
SUV39H2 Act.
SUV39H2 Exp.
ZNF492 Act.
ZNF492 Exp.
4
2
0
−2
−4
PN
PPN
MES
PMES

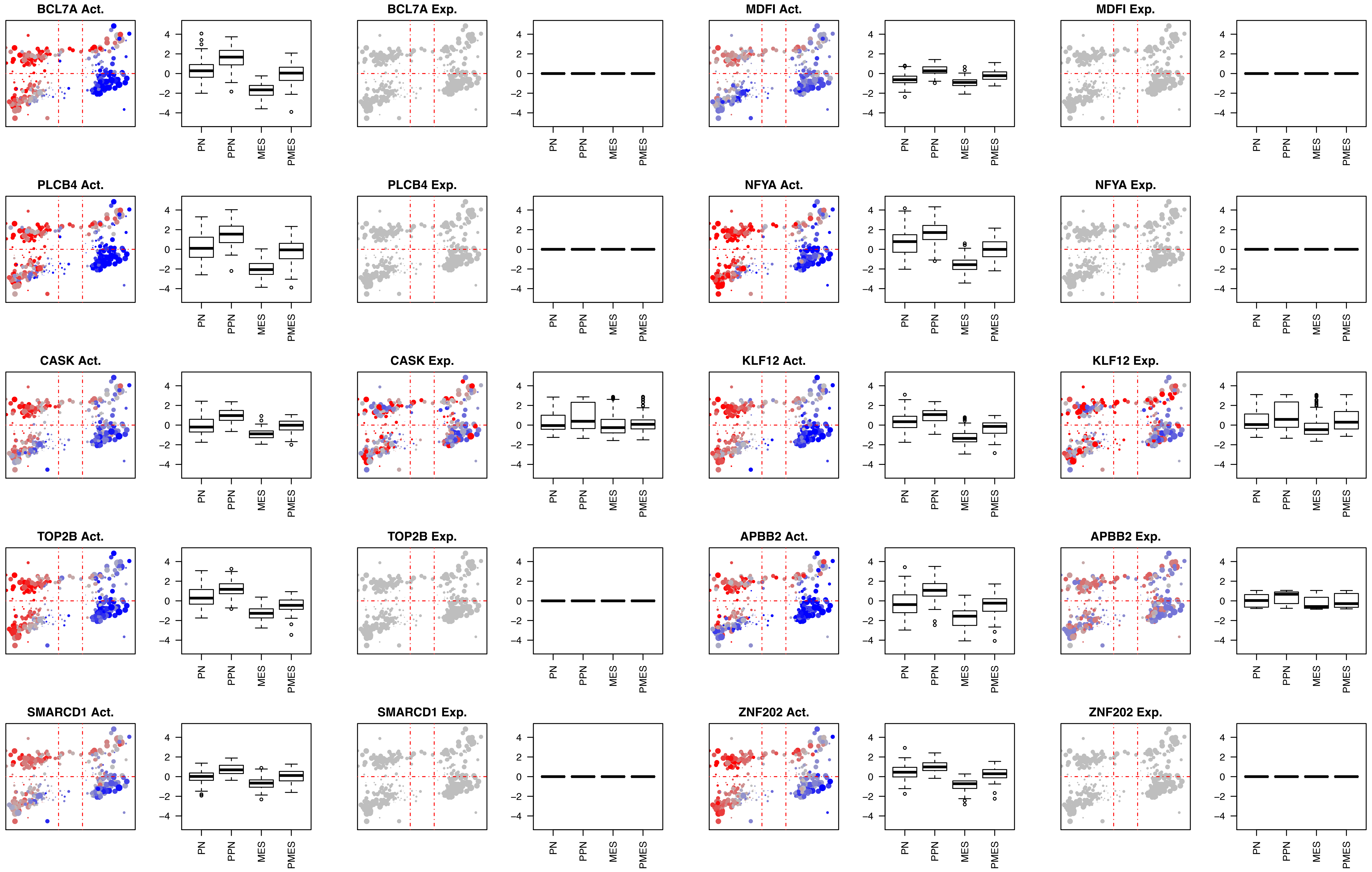
BCL7A Act.
BCL7A Exp.
MDFI Act.
MDFI Exp.
PLCB4 Act.
PLCB4 Exp.
NFYA Act.
NFYA Exp.
CASK Act.
CASK Exp.
KLF12 Act.
KLF12 Exp.
TOP2B Act.
TOP2B Exp.
APBB2 Act.
APBB2 Exp.
SMARCD1 Act.
SMARCD1 Exp.
ZNF202 Act.
ZNF202 Exp.
4
2
0
−2
−4
PN
PPN
MES
PMES

**Proteins aberrantly activated in the MES quadrant.**

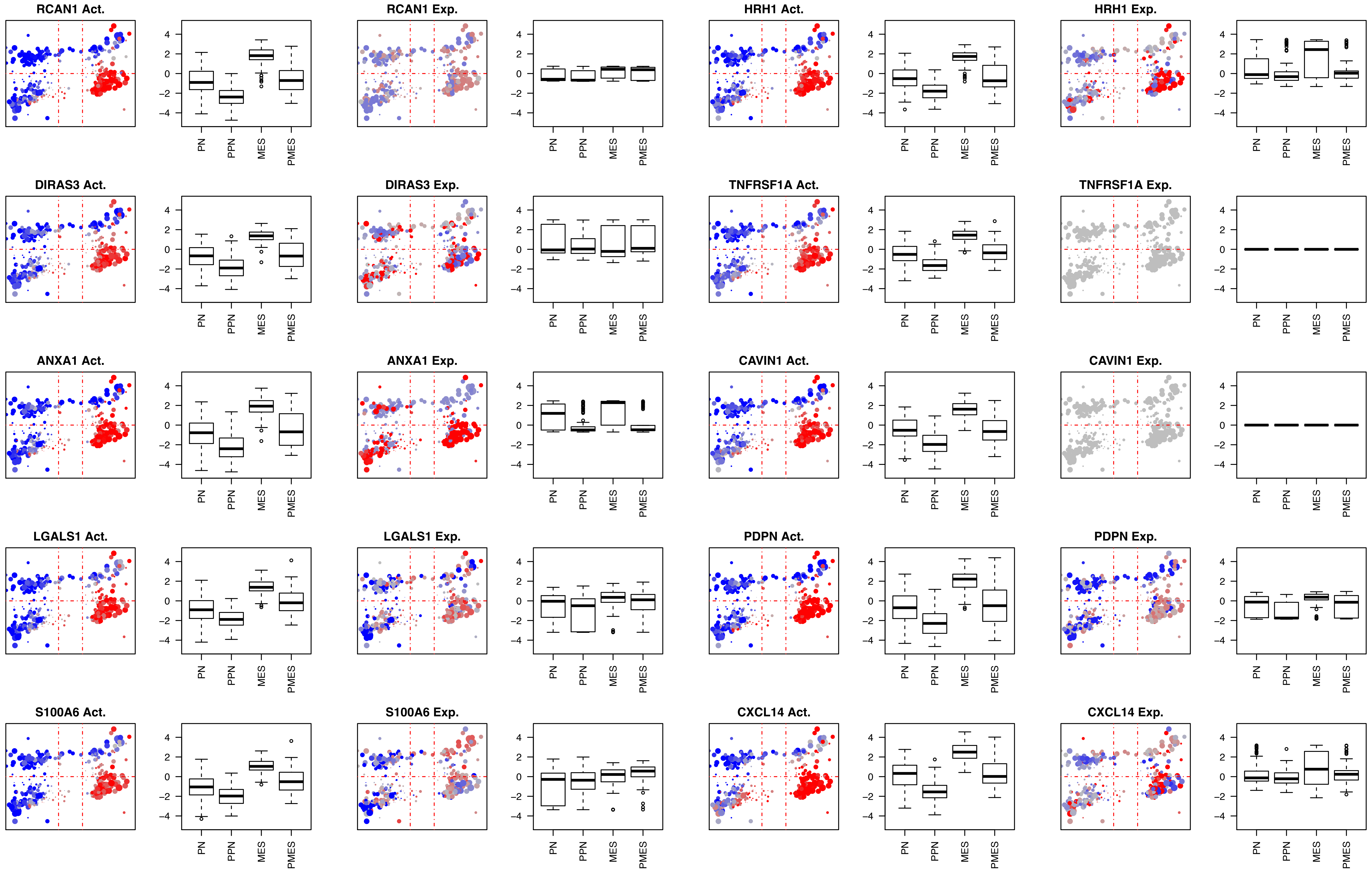
RCAN1 Act.
RCAN1 Exp.
HRH1 Act.
HRH1 Exp.
DIRAS3 Act.
DIRAS3 Exp.
TNFRSF1A Act.
TNFRSF1A Exp.
ANXA1 Act.
ANXA1 Exp.
CAVIN1 Act.
CAVIN1 Exp.
LGALS1 Act.
LGALS1 Exp.
PDPN Act.
PDPN Exp.
S100A6 Act.
S100A6 Exp.
CXCL14 Act.
CXCL14 Exp.
4
2
0
-2
-4
PN
PPN
MES
PMES

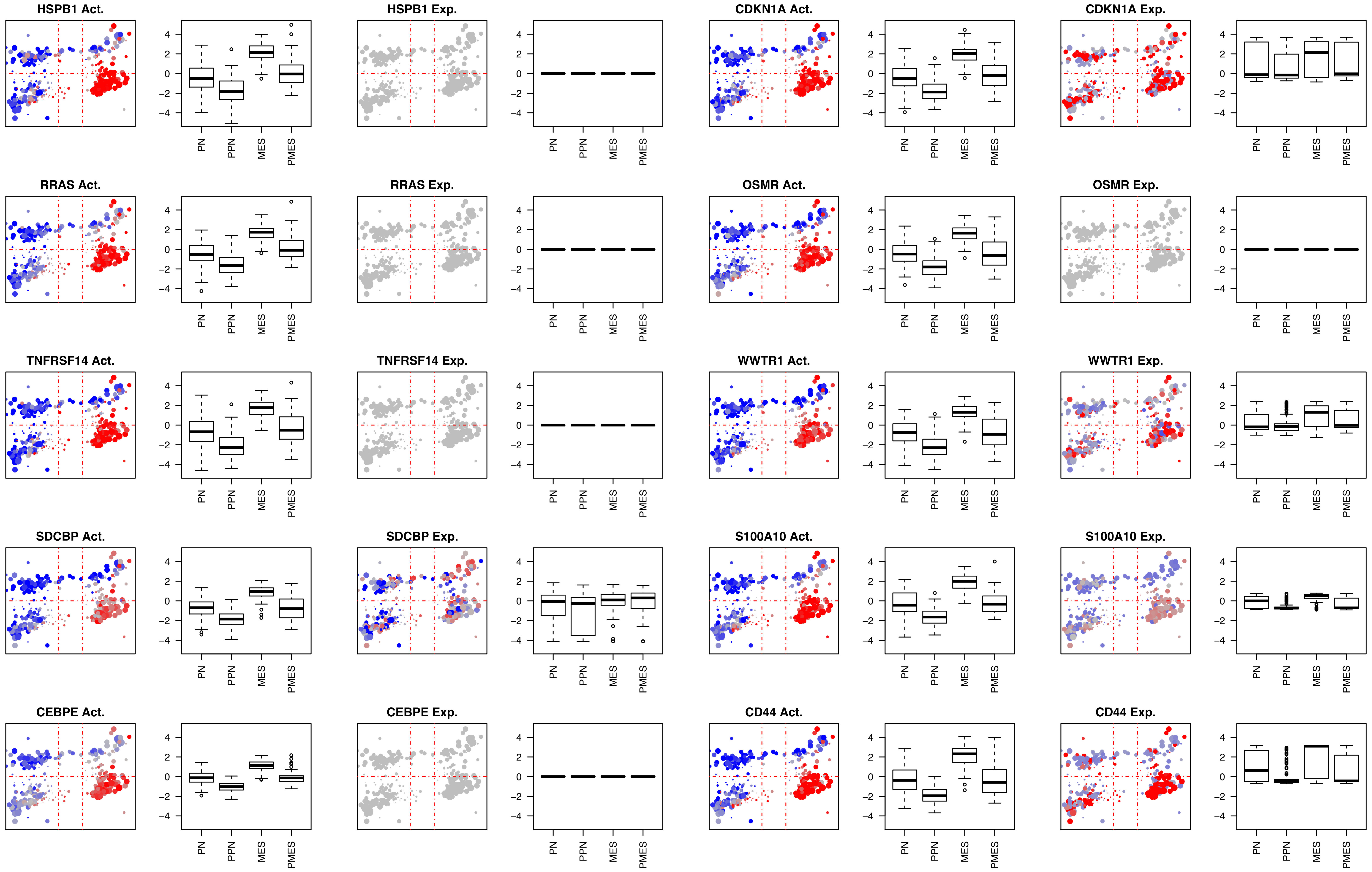
HSPB1 Act.
HSPB1 Exp.
CDKN1A Act.
CDKN1A Exp.
RRAS Act.
RRAS Exp.
OSMR Act.
OSMR Exp.
TNFRSF14 Act.
TNFRSF14 Exp.
WWTR1 Act.
WWTR1 Exp.
SDCBP Act.
SDCBP Exp.
S100A10 Act.
S100A10 Exp.
CEBPE Act.
CEBPE Exp.
CD44 Act.
CD44 Exp.
4
2
0
−2
−4
PN
PPN
MES
PMES

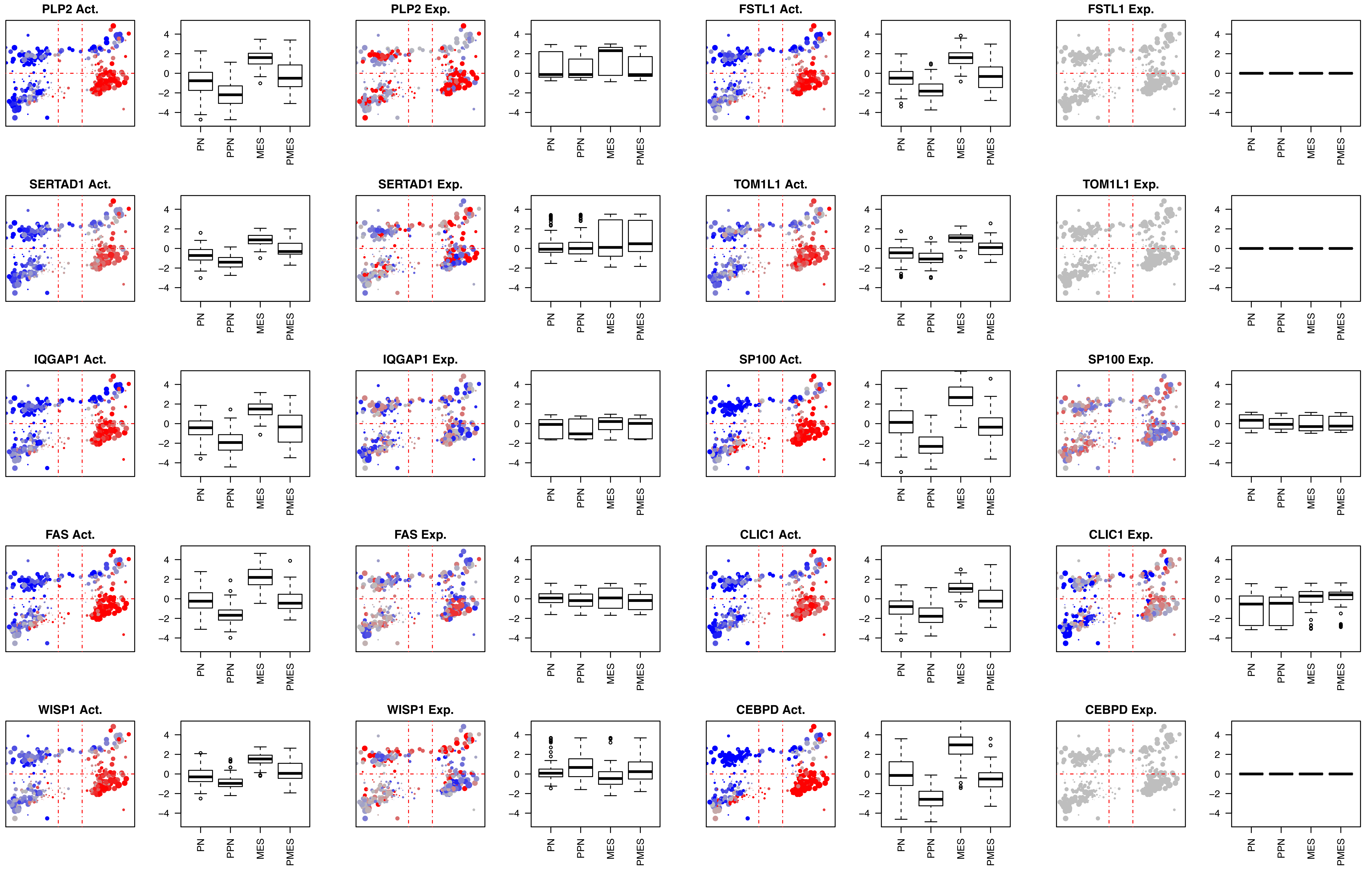
PLP2 Act.
PLP2 Exp.
FSTL1 Act.
FSTL1 Exp.
SERTAD1 Act.
SERTAD1 Exp.
TOM1L1 Act.
TOM1L1 Exp.
IQGAP1 Act.
IQGAP1 Exp.
SP100 Act.
SP100 Exp.
FAS Act.
FAS Exp.
CLIC1 Act.
CLIC1 Exp.
WISP1 Act.
WISP1 Exp.
CEBPD Act.
CEBPD Exp.
4
2
0
-2
-4
PN
PPN
MES
PMES

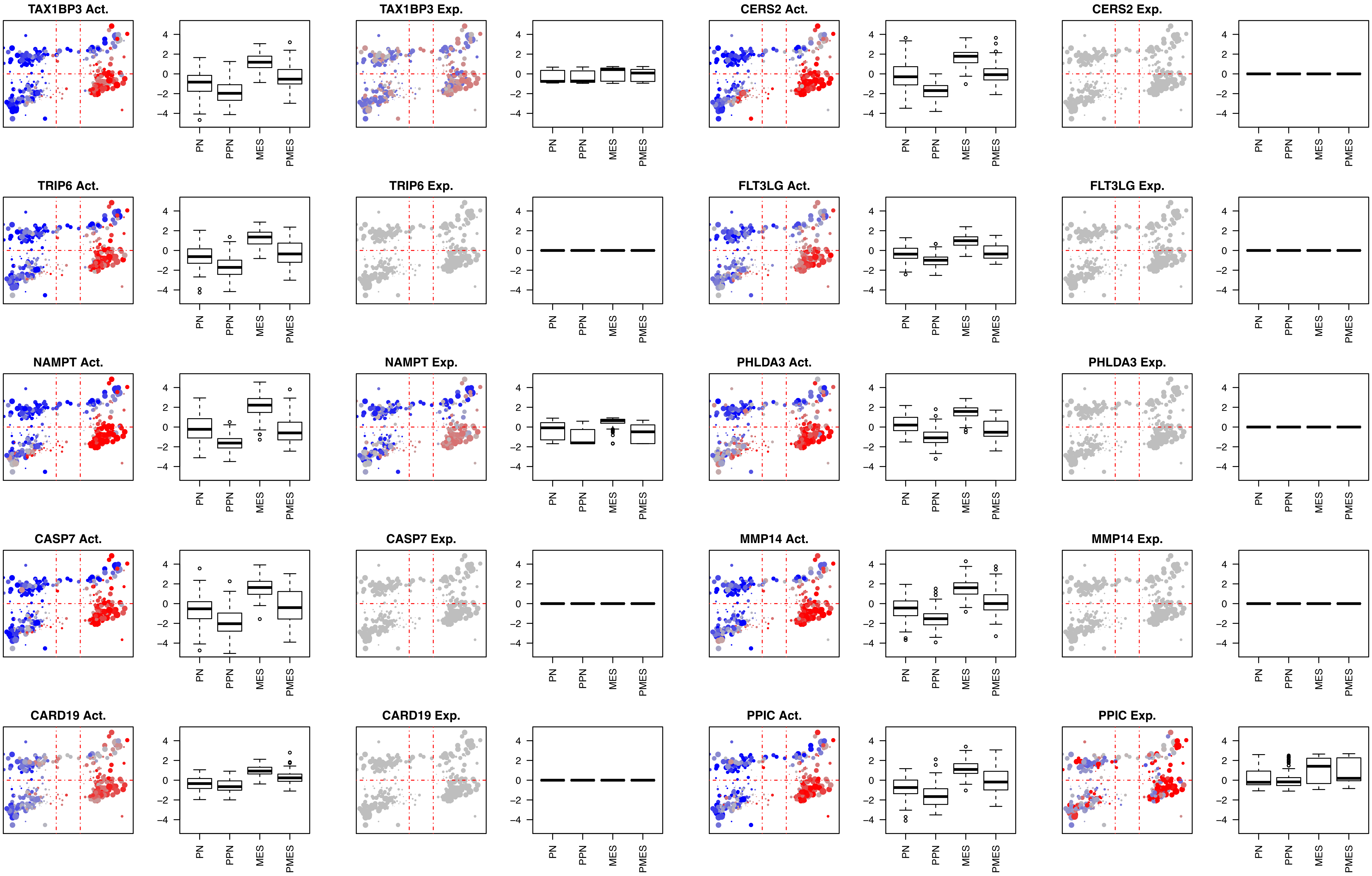

TAX1BP3 Act.
TAX1BP3 Exp.
CERS2 Act.
CERS2 Exp.
TRIP6 Act.
TRIP6 Exp.
FLT3LG Act.
FLT3LG Exp.
NAMPT Act.
NAMPT Exp.
PHLDA3 Act.
PHLDA3 Exp.
CASP7 Act.
CASP7 Exp.
MMP14 Act.
MMP14 Exp.
CARD19 Act.
CARD19 Exp.
PPIC Act.
PPIC Exp.
4
2
0
-2
-4
PN
PPN
MES
PMES

**Proteins aberrantly activated in the PMES quadrant.**

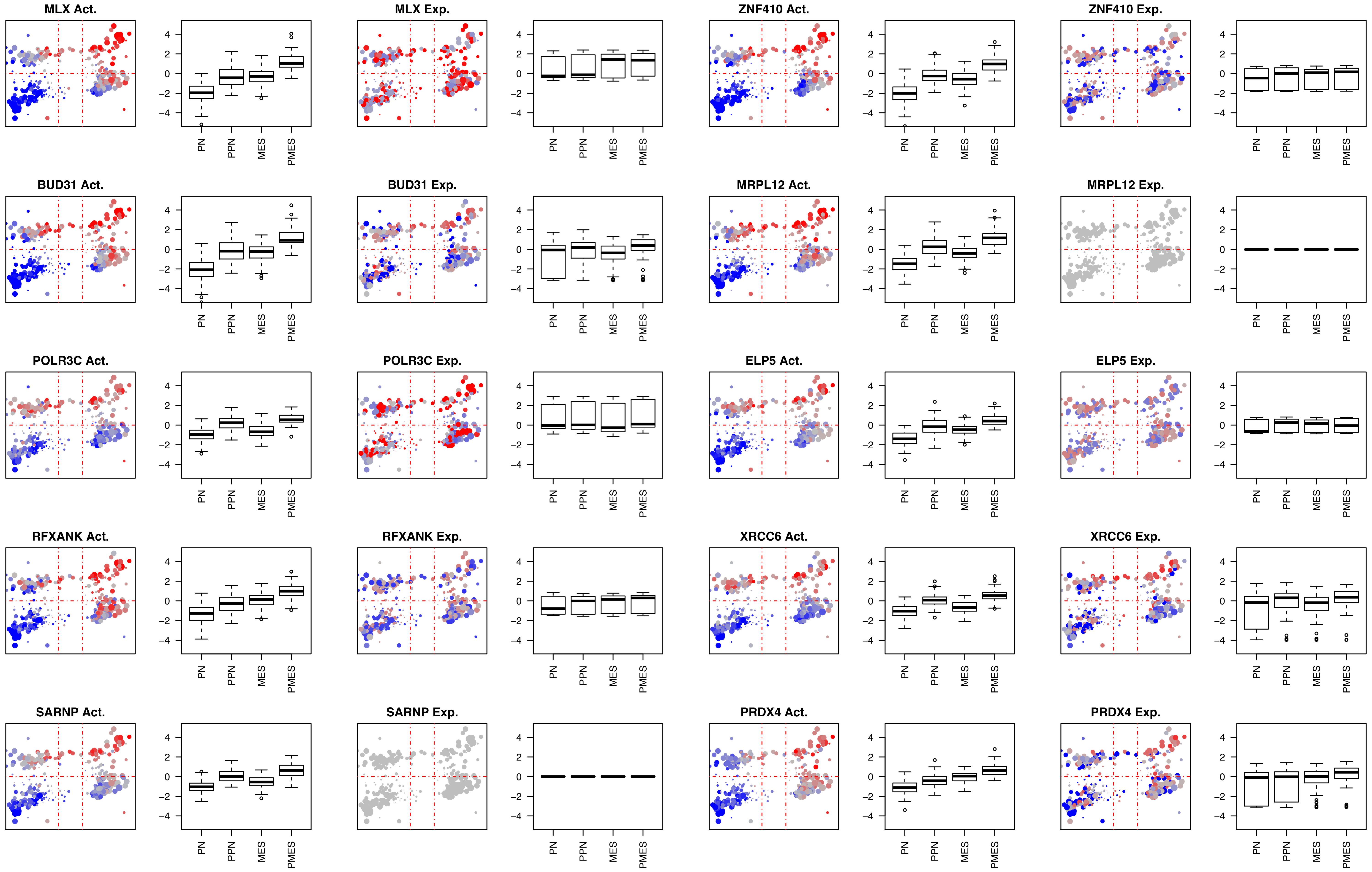
MLX Act.
4 2 0 −2 −4
PN PPN MES PMES
MLX Exp.
4 2 0 −2 −4
PN PPN MES PMES
ZNF410 Act.
4 2 0 −2 −4
PN PPN MES PMES
ZNF410 Exp.
4 2 0 −2 −4
PN PPN MES PMES
BUD31 Act.
4 2 0 −2 −4
PN PPN MES PMES
BUD31 Exp.
4 2 0 −2 −4
PN PPN MES PMES
MRPL12 Act.
4 2 0 −2 −4
PN PPN MES PMES
MRPL12 Exp.
4 2 0 −2 −4
PN PPN MES PMES
POLR3C Act.
4 2 0 −2 −4
PN PPN MES PMES
POLR3C Exp.
4 2 0 −2 −4
PN PPN MES PMES
ELP5 Act.
4 2 0 −2 −4
PN PPN MES PMES
ELP5 Exp.
4 2 0 −2 −4
PN PPN MES PMES
RFXANK Act.
4 2 0 −2 −4
PN PPN MES PMES
RFXANK Exp.
4 2 0 −2 −4
PN PPN MES PMES
XRCC6 Act.
4 2 0 −2 −4
PN PPN MES PMES
XRCC6 Exp.
4 2 0 −2 −4
PN PPN MES PMES
SARNP Act.
4 2 0 −2 −4
PN PPN MES PMES
SARNP Exp.
4 2 0 −2 −4
PN PPN MES PMES
PRDX4 Act.
4 2 0 −2 −4
PN PPN MES PMES
PRDX4 Exp.
4 2 0 −2 −4
PN PPN MES PMES

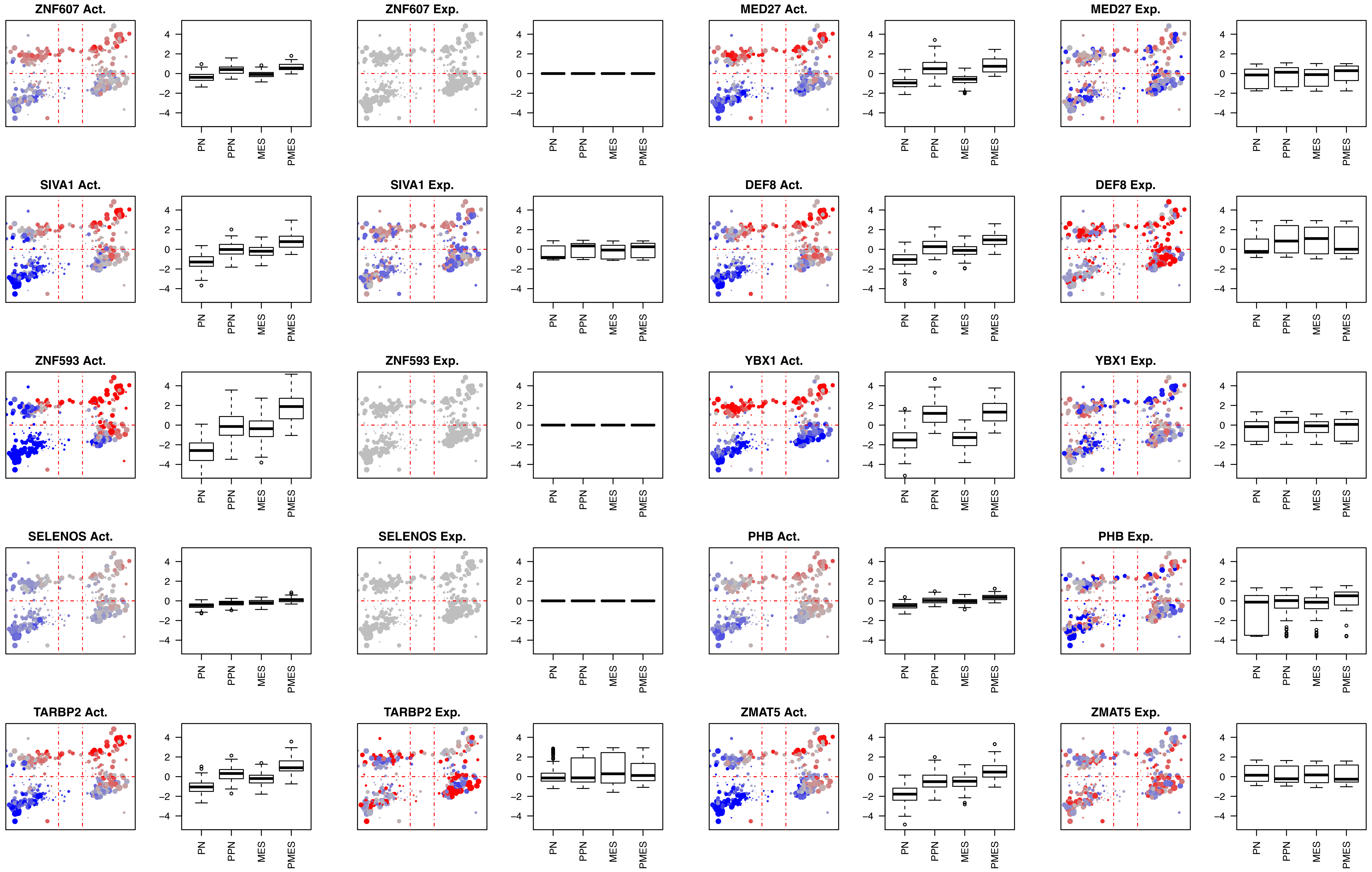

ZNF607 Act.
ZNF607 Exp.
MED27 Act.
MED27 Exp.
SIVA1 Act.
SIVA1 Exp.
DEF8 Act.
DEF8 Exp.
ZNF593 Act.
ZNF593 Exp.
YBX1 Act.
YBX1 Exp.
SELENOS Act.
SELENOS Exp.
PHB Act.
PHB Exp.
TARBP2 Act.
TARBP2 Exp.
ZMAT5 Act.
ZMAT5 Exp.
4
2
0
-2
-4
PN
PPN
MES
PMES

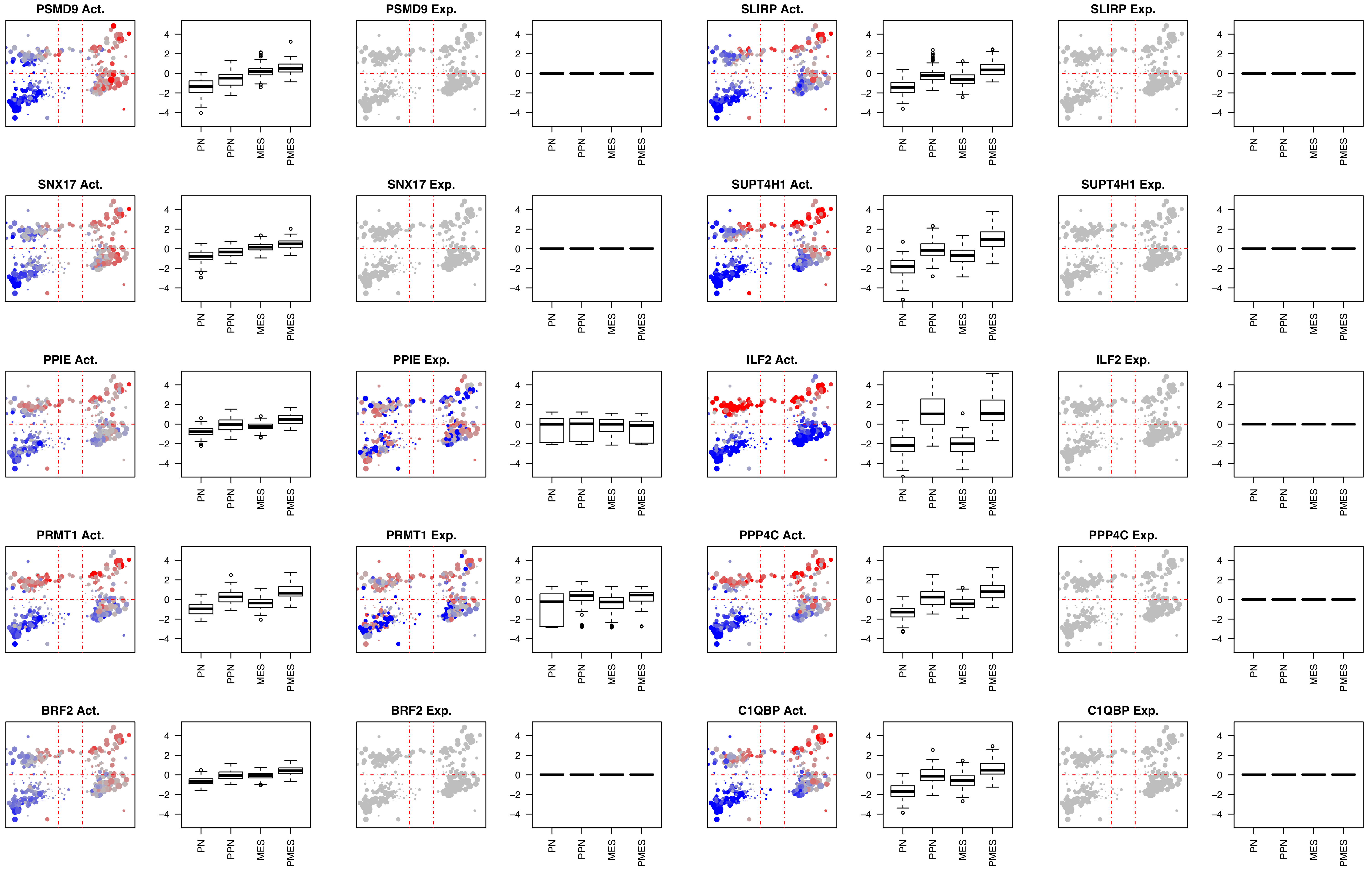
PSMD9 Act.
PSMD9 Exp.
SLIRP Act.
SLIRP Exp.
SNX17 Act.
SNX17 Exp.
SUPT4H1 Act.
SUPT4H1 Exp.
PPIE Act.
PPIE Exp.
ILF2 Act.
ILF2 Exp.
PRMT1 Act.
PRMT1 Exp.
PPP4C Act.
PPP4C Exp.
BRF2 Act.
BRF2 Exp.
C1QBP Act.
C1QBP Exp.
4
2
0
−2
−4
PN
PPN
MES
PMES

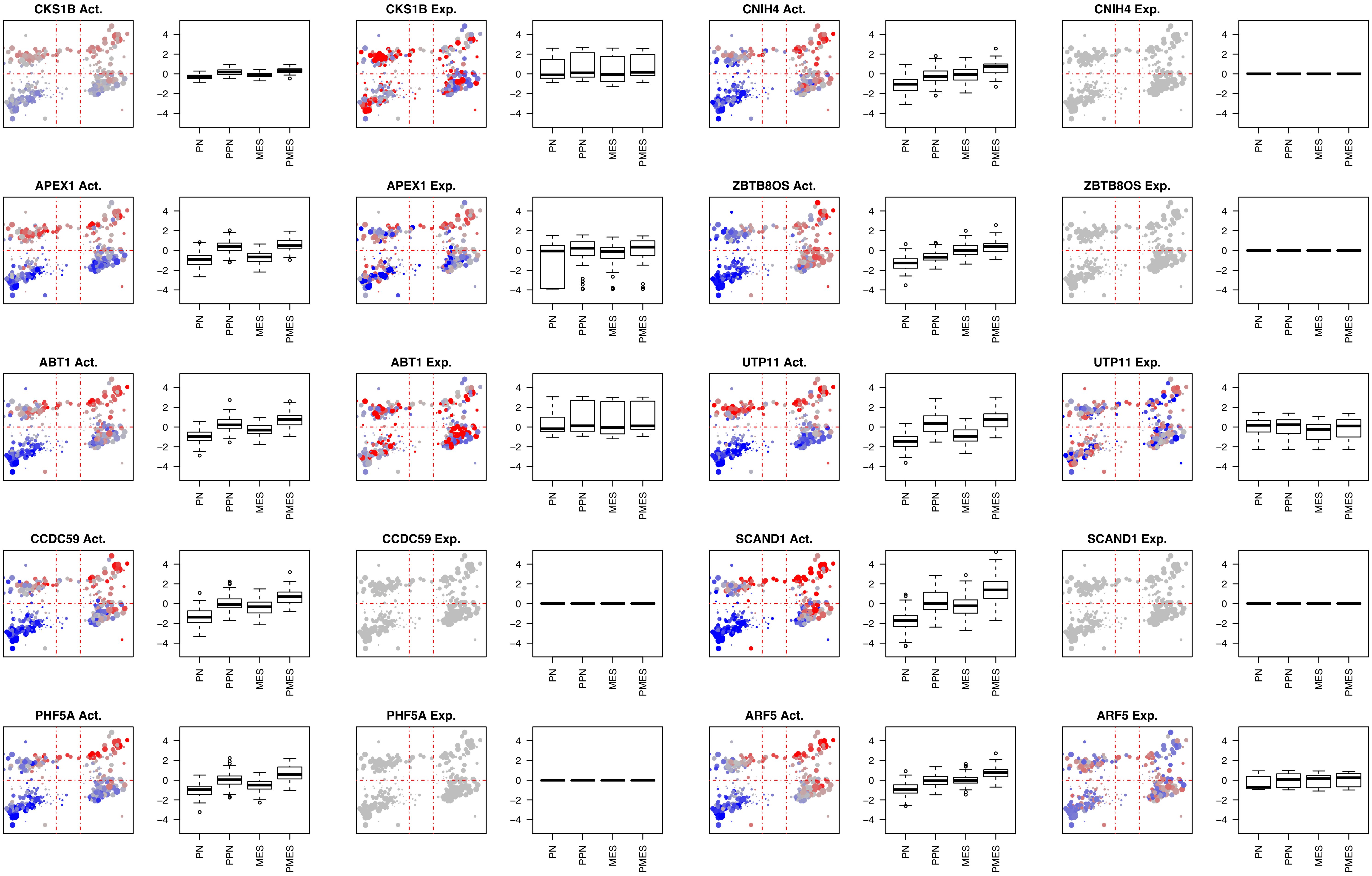
CKS1B Act.
CKS1B Exp.
CNIH4 Act.
CNIH4 Exp.
APEX1 Act.
APEX1 Exp.
ZBTB8OS Act.
ZBTB8OS Exp.
ABT1 Act.
ABT1 Exp.
UTP11 Act.
UTP11 Exp.
CCDC59 Act.
CCDC59 Exp.
SCAND1 Act.
SCAND1 Exp.
PHF5A Act.
PHF5A Exp.
ARF5 Act.
ARF5 Exp.
4
2
0
−2
−4
PN
PPN
MES
PMES

Fig5SupFig-3

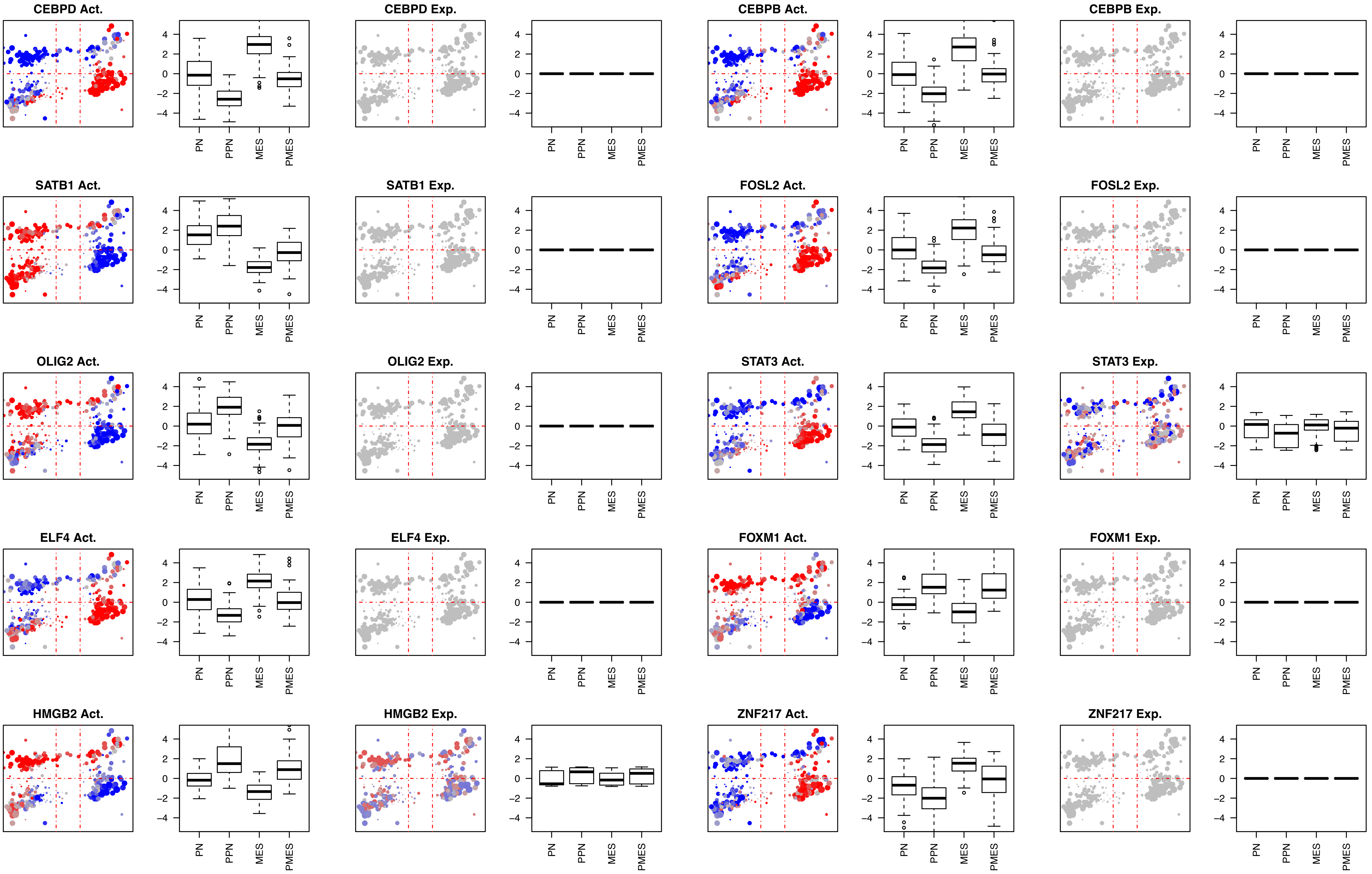
CEBPD Act.
CEBPD Exp.
CEBPB Act.
CEBPB Exp.
SATB1 Act.
SATB1 Exp.
FOSL2 Act.
FOSL2 Exp.
OLIG2 Act.
OLIG2 Exp.
STAT3 Act.
STAT3 Exp.
ELF4 Act.
ELF4 Exp.
FOXM1 Act.
FOXM1 Exp.
HMGB2 Act.
HMGB2 Exp.
ZNF217 Act.
ZNF217 Exp.
4
2
0
-2
-4
PN
PPN
MES
PMES

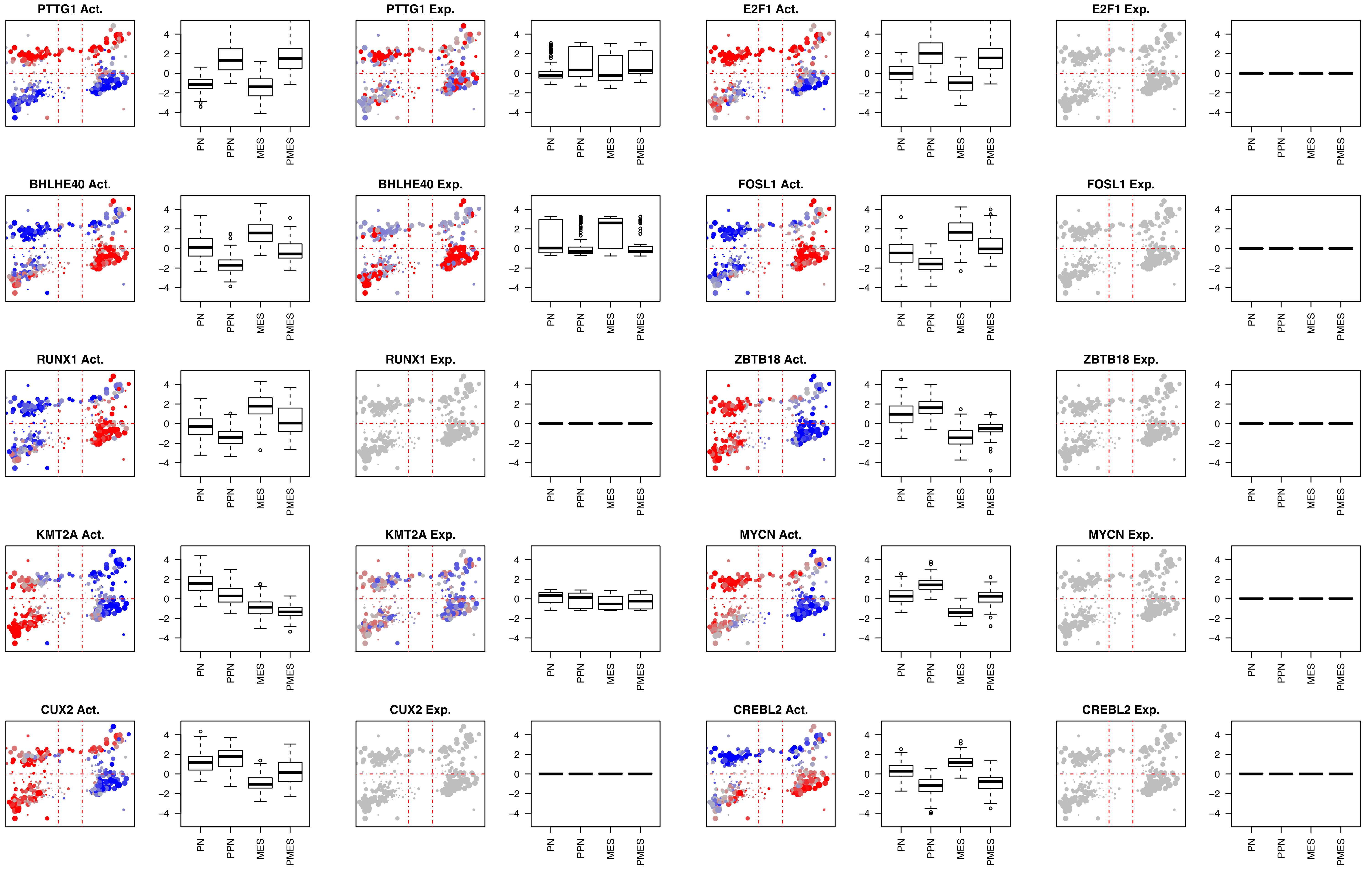
PTTG1 Act.
PTTG1 Exp.
E2F1 Act.
E2F1 Exp.
BHLHE40 Act.
BHLHE40 Exp.
FOSL1 Act.
FOSL1 Exp.
RUNX1 Act.
RUNX1 Exp.
ZBTB18 Act.
ZBTB18 Exp.
KMT2A Act.
KMT2A Exp.
MYCN Act.
MYCN Exp.
CUX2 Act.
CUX2 Exp.
CREBL2 Act.
CREBL2 Exp.
4
2
0
-2
-4
PN
PPN
MES
PMES

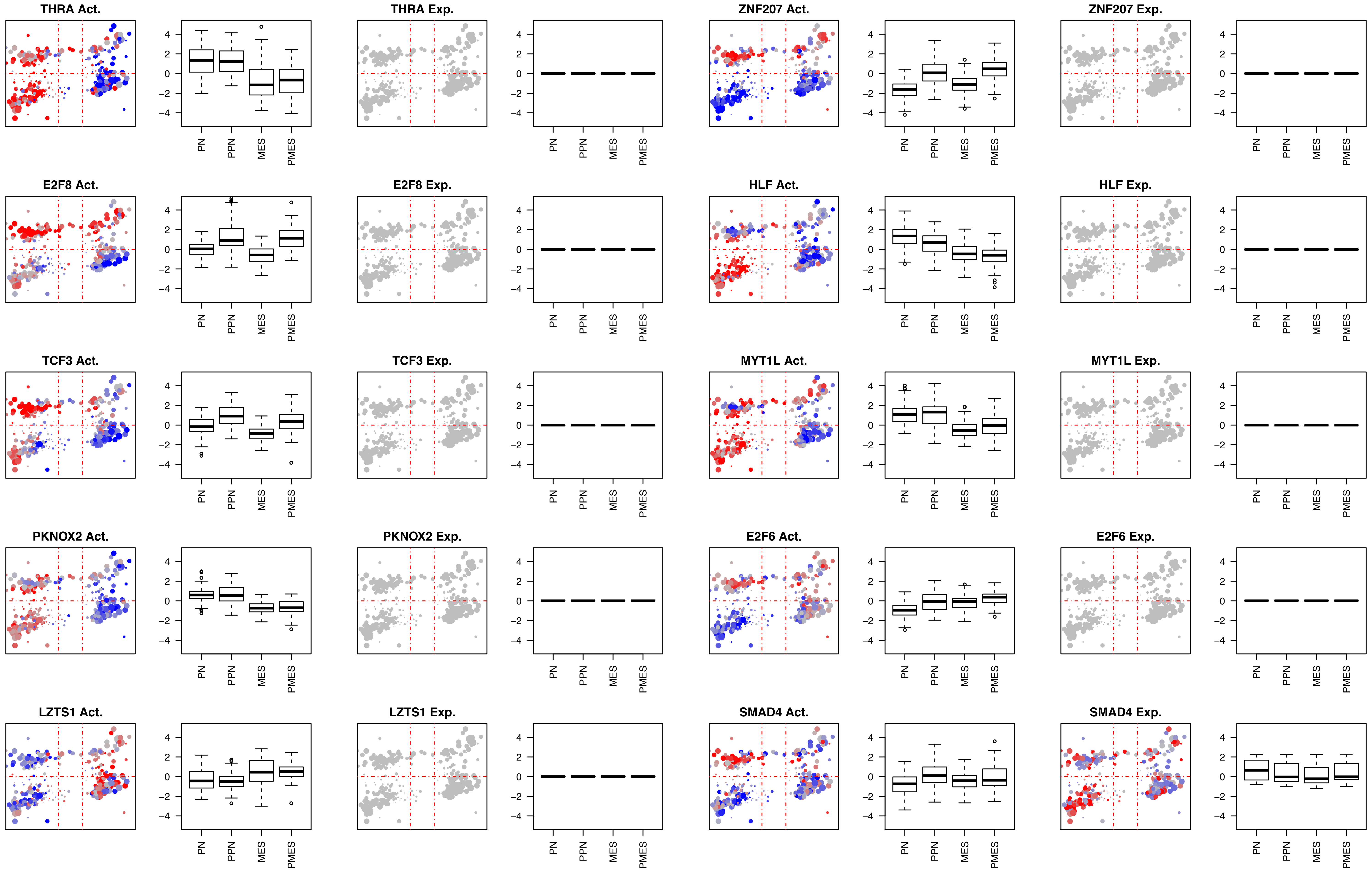
THRA Act.
THRA Exp.
ZNF207 Act.
ZNF207 Exp.
E2F8 Act.
E2F8 Exp.
HLF Act.
HLF Exp.
TCF3 Act.
TCF3 Exp.
MYT1L Act.
MYT1L Exp.
PKNOX2 Act.
PKNOX2 Exp.
E2F6 Act.
E2F6 Exp.
LZTS1 Act.
LZTS1 Exp.
SMAD4 Act.
SMAD4 Exp.
4
2
0
−2
−4
PN
PPN
MES
PMES

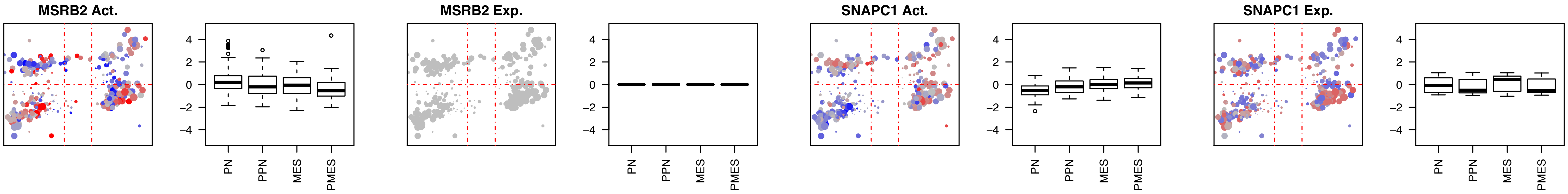
MSRB2 Act.
MSRB2 Exp.
SNAPC1 Act.
SNAPC1 Exp.
4
2
0
−2
−4
PN
PPN
MES
PMES

Fig5SupFig-4

#### Proteins aberrantly activated in the Proneural quadrants (PN + PPN)

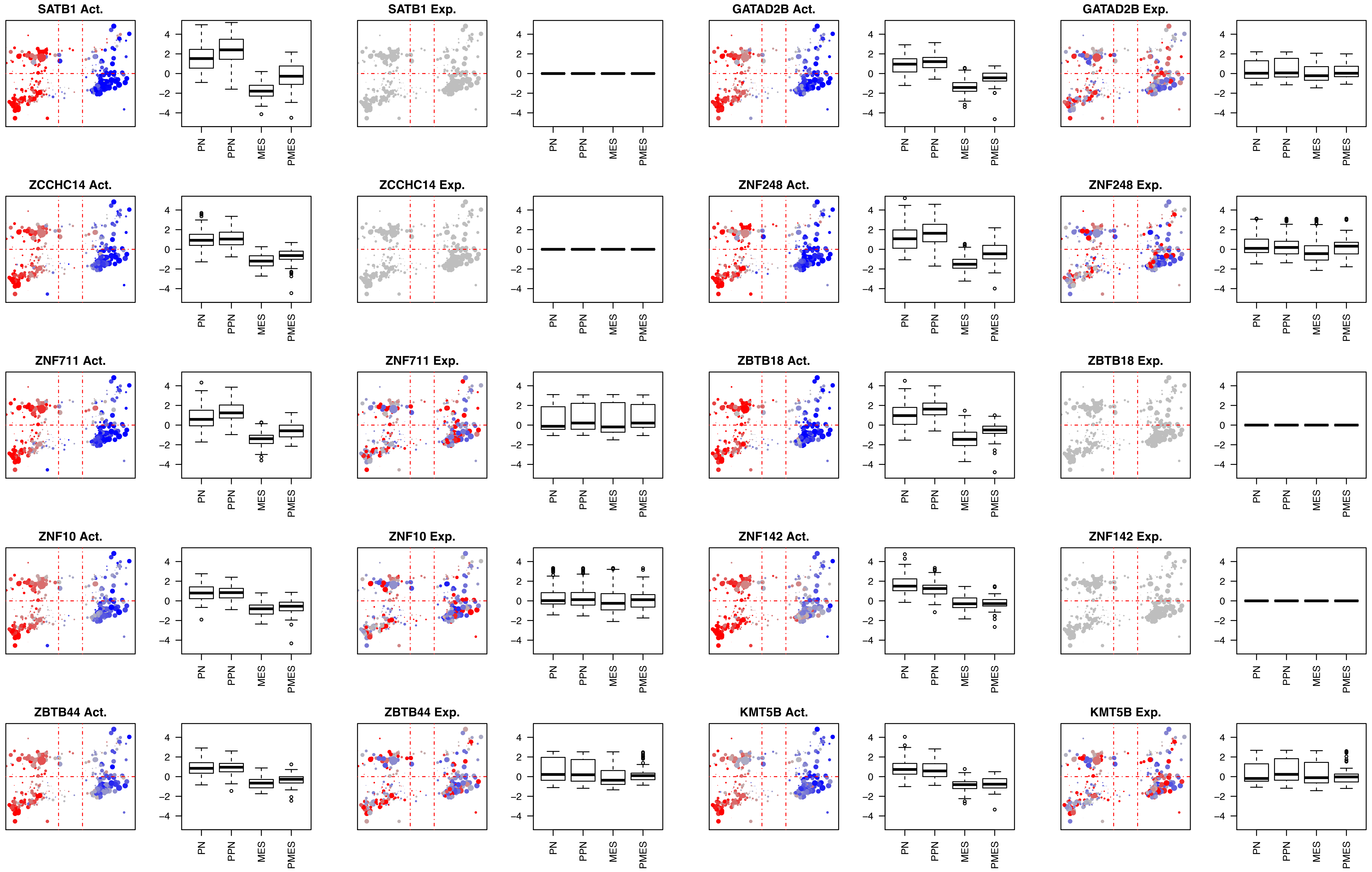
SATB1 Act.
SATB1 Exp.
GATAD2B Act.
GATAD2B Exp.
ZCCHC14 Act.
ZCCHC14 Exp.
ZNF248 Act.
ZNF248 Exp.
ZNF711 Act.
ZNF711 Exp.
ZBTB18 Act.
ZBTB18 Exp.
ZNF10 Act.
ZNF10 Exp.
ZNF142 Act.
ZNF142 Exp.
ZBTB44 Act.
ZBTB44 Exp.
KMT5B Act.
KMT5B Exp.
4
2
0
-2
-4
PN
PPN
MES
PMES

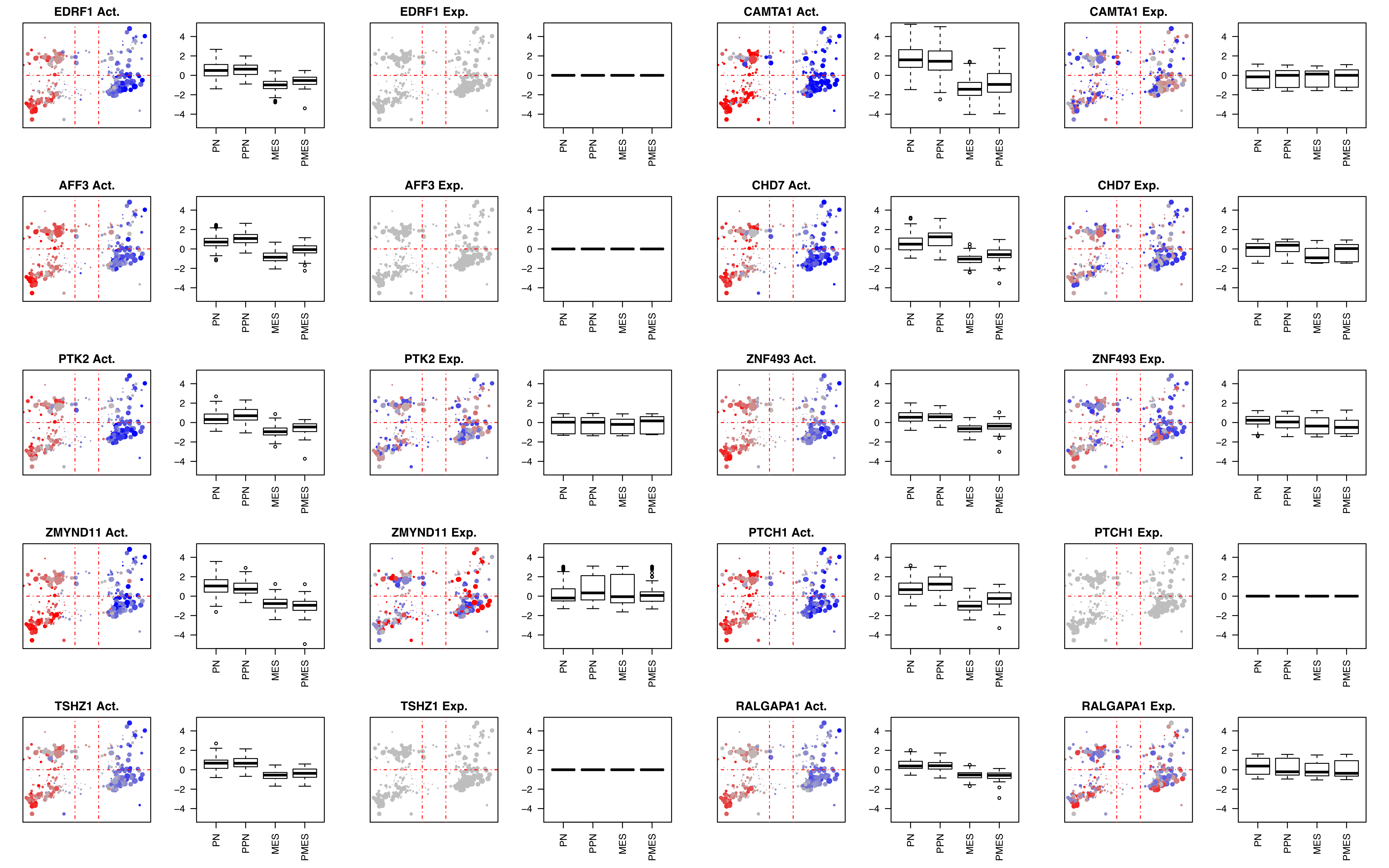
EDRF1 Act.
EDRF1 Exp.
CAMTA1 Act.
CAMTA1 Exp.
AFF3 Act.
AFF3 Exp.
CHD7 Act.
CHD7 Exp.
PTK2 Act.
PTK2 Exp.
ZNF493 Act.
ZNF493 Exp.
ZMYND11 Act.
ZMYND11 Exp.
PTCH1 Act.
PTCH1 Exp.
TSHZ1 Act.
TSHZ1 Exp.
RALGAPA1 Act.
RALGAPA1 Exp.
4
2
0
−2
−4
PN
PPN
MES
PMES

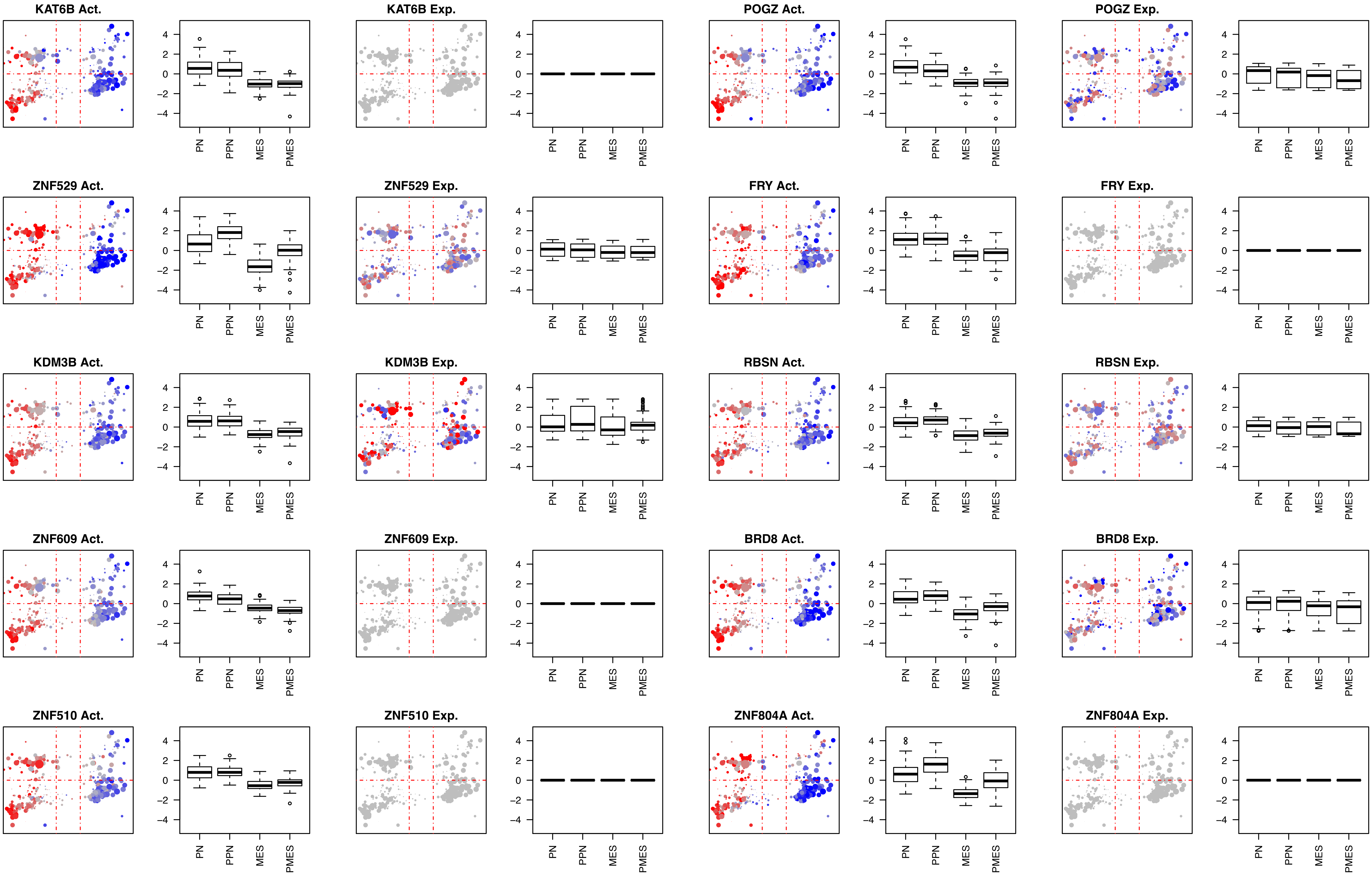
KAT6B Act.
KAT6B Exp.
POGZ Act.
POGZ Exp.
ZNF529 Act.
ZNF529 Exp.
FRY Act.
FRY Exp.
KDM3B Act.
KDM3B Exp.
RBSN Act.
RBSN Exp.
ZNF609 Act.
ZNF609 Exp.
BRD8 Act.
BRD8 Exp.
ZNF510 Act.
ZNF510 Exp.
ZNF804A Act.
ZNF804A Exp.
4
2
0
−2
−4
PN
PPN
MES
PMES

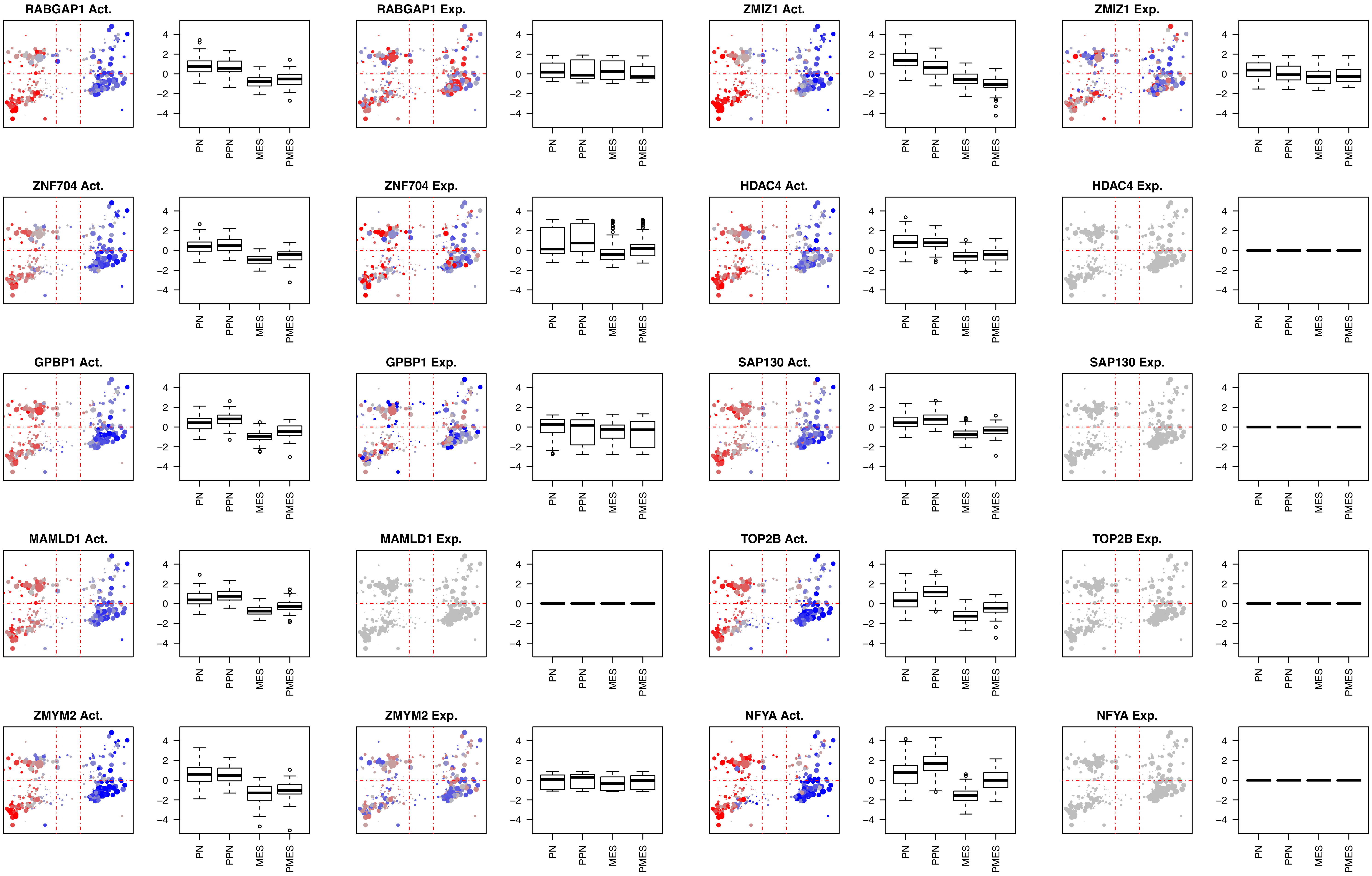
RABGAP1 Act.
RABGAP1 Exp.
ZMIZ1 Act.
ZMIZ1 Exp.
ZNF704 Act.
ZNF704 Exp.
HDAC4 Act.
HDAC4 Exp.
GPBP1 Act.
GPBP1 Exp.
SAP130 Act.
SAP130 Exp.
MAMLD1 Act.
MAMLD1 Exp.
TOP2B Act.
TOP2B Exp.
ZMYM2 Act.
ZMYM2 Exp.
NFYA Act.
NFYA Exp.
4
2
0
-2
-4
PN
PPN
MES
PMES

### Proteins aberrantly activated in the Mesenchymal quadrants (MES + PMES)

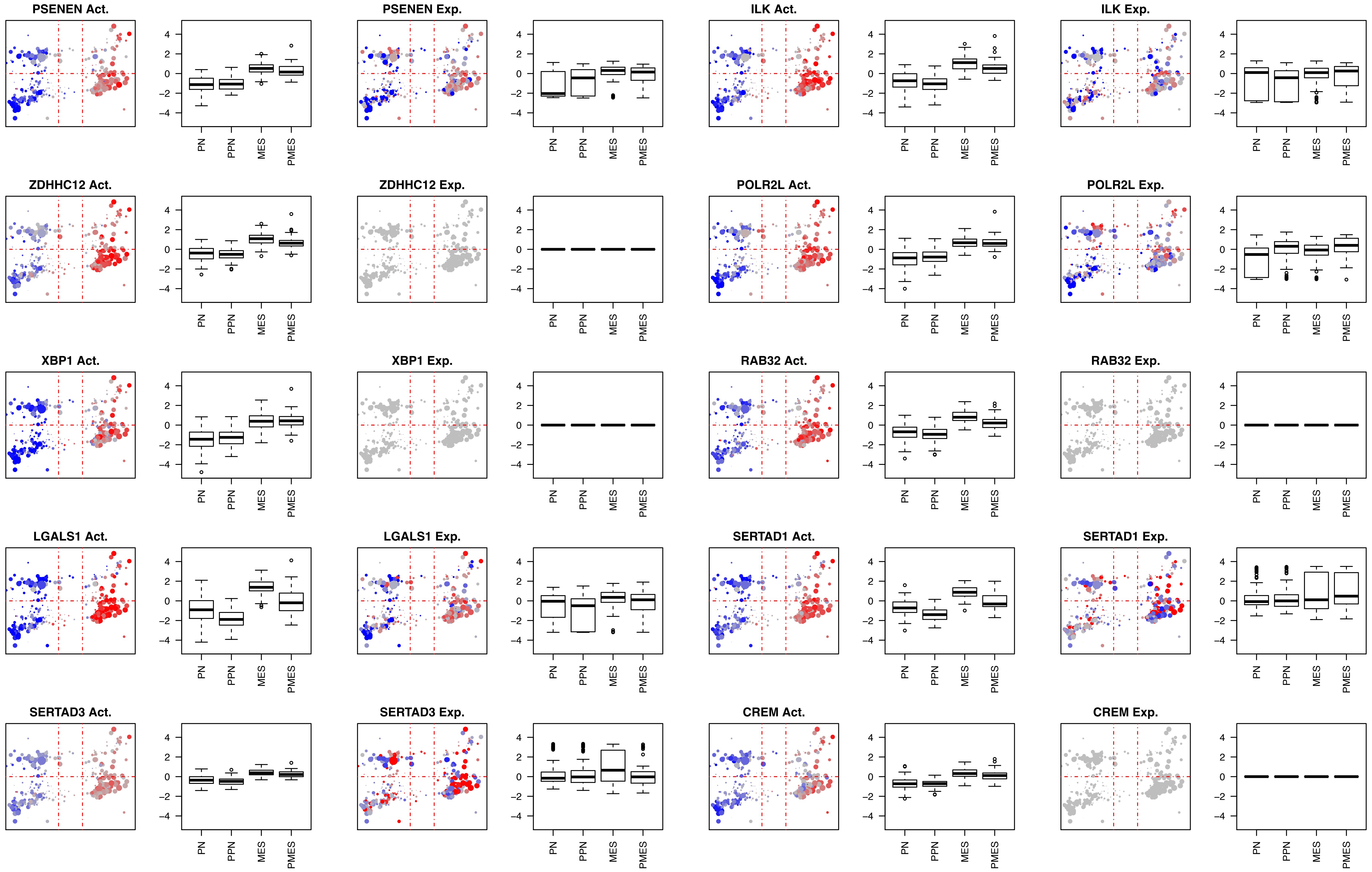
PSENEN Act.
PSENEN Exp.
ILK Act.
ILK Exp.
ZDHHC12 Act.
ZDHHC12 Exp.
POLR2L Act.
POLR2L Exp.
XBP1 Act.
XBP1 Exp.
RAB32 Act.
RAB32 Exp.
LGALS1 Act.
LGALS1 Exp.
SERTAD1 Act.
SERTAD1 Exp.
SERTAD3 Act.
SERTAD3 Exp.
CREM Act.
CREM Exp.
4
2
0
−2
−4
PN
PPN
MES
PMES

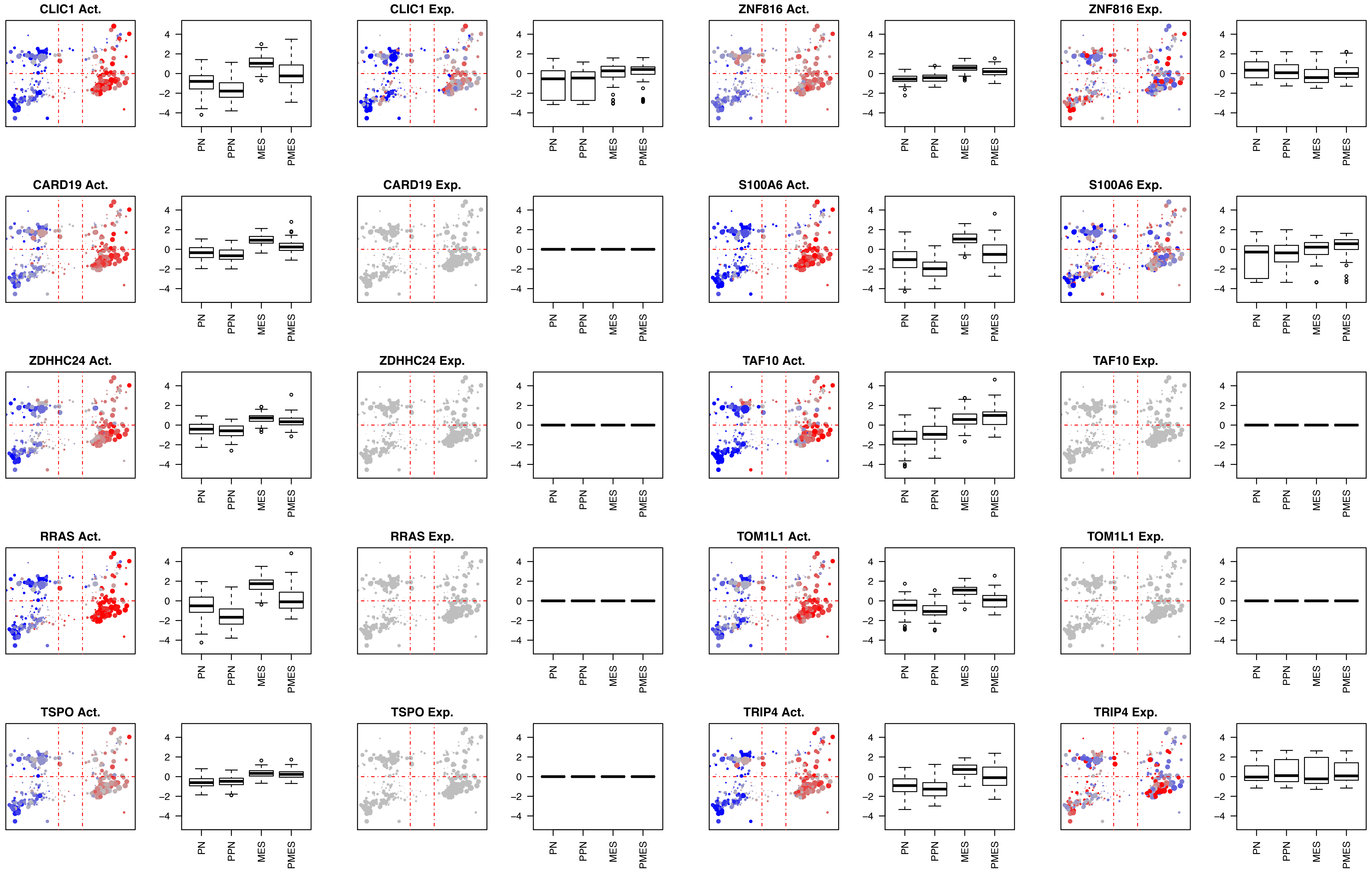
CLIC1 Act.
CLIC1 Exp.
ZNF816 Act.
ZNF816 Exp.
CARD19 Act.
CARD19 Exp.
S100A6 Act.
S100A6 Exp.
ZDHHC24 Act.
ZDHHC24 Exp.
TAF10 Act.
TAF10 Exp.
RRAS Act.
RRAS Exp.
TOM1L1 Act.
TOM1L1 Exp.
TSPO Act.
TSPO Exp.
TRIP4 Act.
TRIP4 Exp.
4
2
0
-2
-4
PN
PPN
MES
PMES

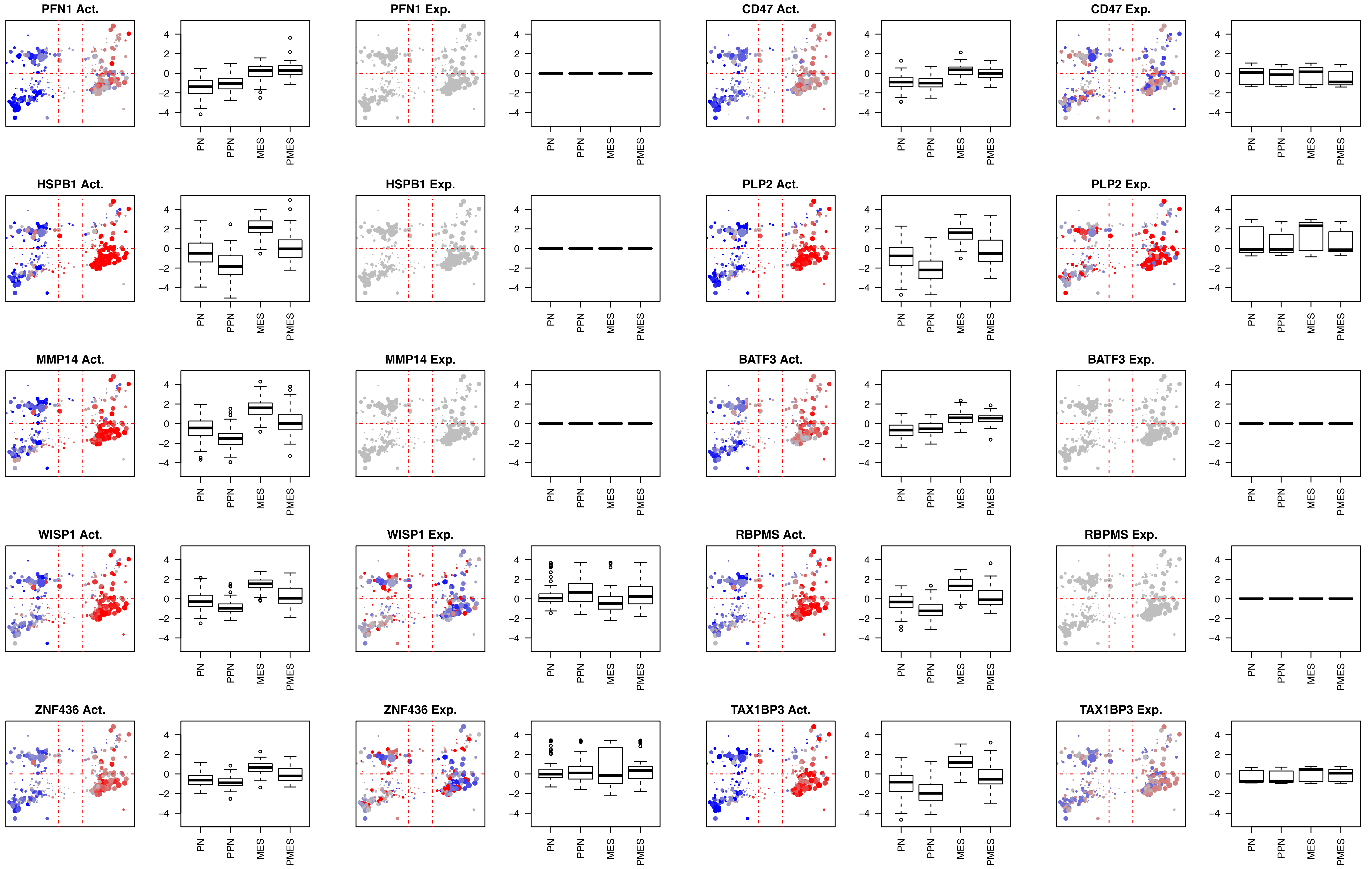
PFN1 Act.
4
2
0
-2
-4
PN
PPN
MES
PMES
PFN1 Exp.
CD47 Act.
CD47 Exp.
HSPB1 Act.
HSPB1 Exp.
PLP2 Act.
PLP2 Exp.
MMP14 Act.
MMP14 Exp.
BATF3 Act.
BATF3 Exp.
WISP1 Act.
WISP1 Exp.
RBPMS Act.
RBPMS Exp.
ZNF436 Act.
ZNF436 Exp.
TAX1BP3 Act.
TAX1BP3 Exp.

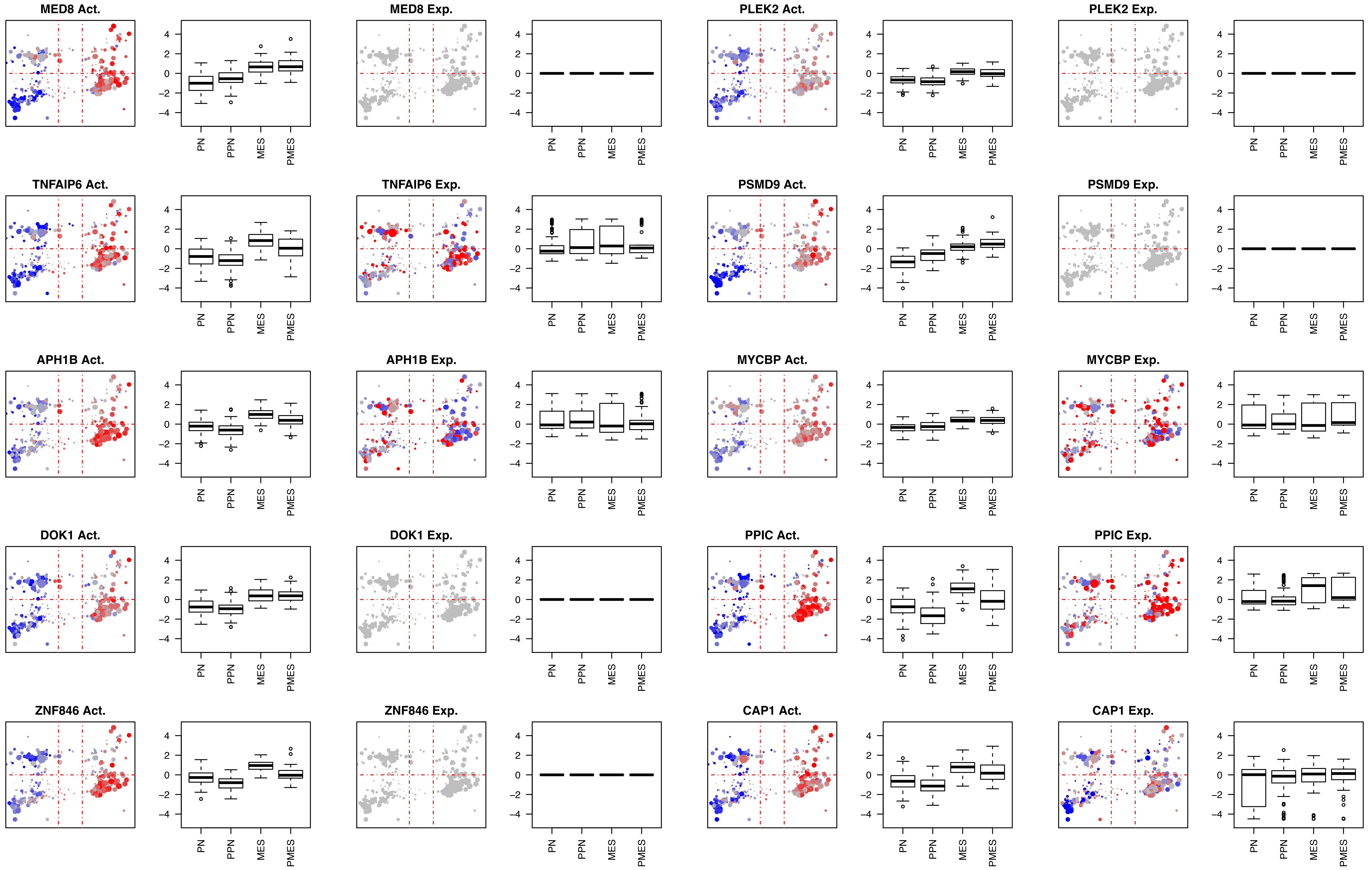
MED8 Act.
MED8 Exp.
PLEK2 Act.
PLEK2 Exp.
TNFAIP6 Act.
TNFAIP6 Exp.
PSMD9 Act.
PSMD9 Exp.
APH1B Act.
APH1B Exp.
MYCBP Act.
MYCBP Exp.
DOK1 Act.
DOK1 Exp.
PPIC Act.
PPIC Exp.
ZNF846 Act.
ZNF846 Exp.
CAP1 Act.
CAP1 Exp.
4
2
0
−2
−4
PN
PPN
MES
PMES

Fig5SupFig-5

**Proteins aberrantly activated in the PRO+ quadrants.**

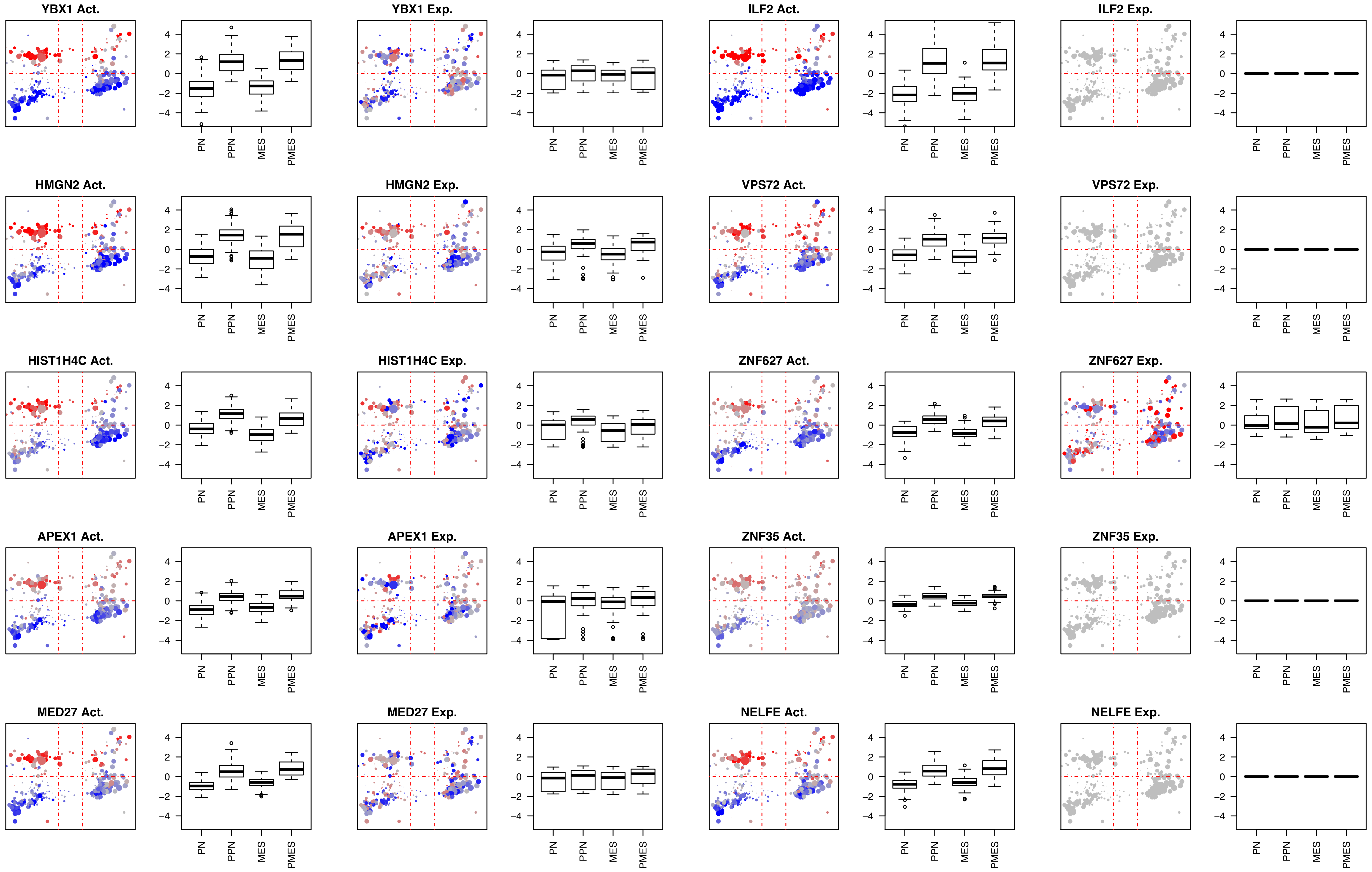
YBX1 Act.
YBX1 Exp.
ILF2 Act.
ILF2 Exp.
HMGN2 Act.
HMGN2 Exp.
VPS72 Act.
VPS72 Exp.
HIST1H4C Act.
HIST1H4C Exp.
ZNF627 Act.
ZNF627 Exp.
APEX1 Act.
APEX1 Exp.
ZNF35 Act.
ZNF35 Exp.
MED27 Act.
MED27 Exp.
NELFE Act.
NELFE Exp.
4
2
0
-2
-4
PN
PPN
MES
PMES

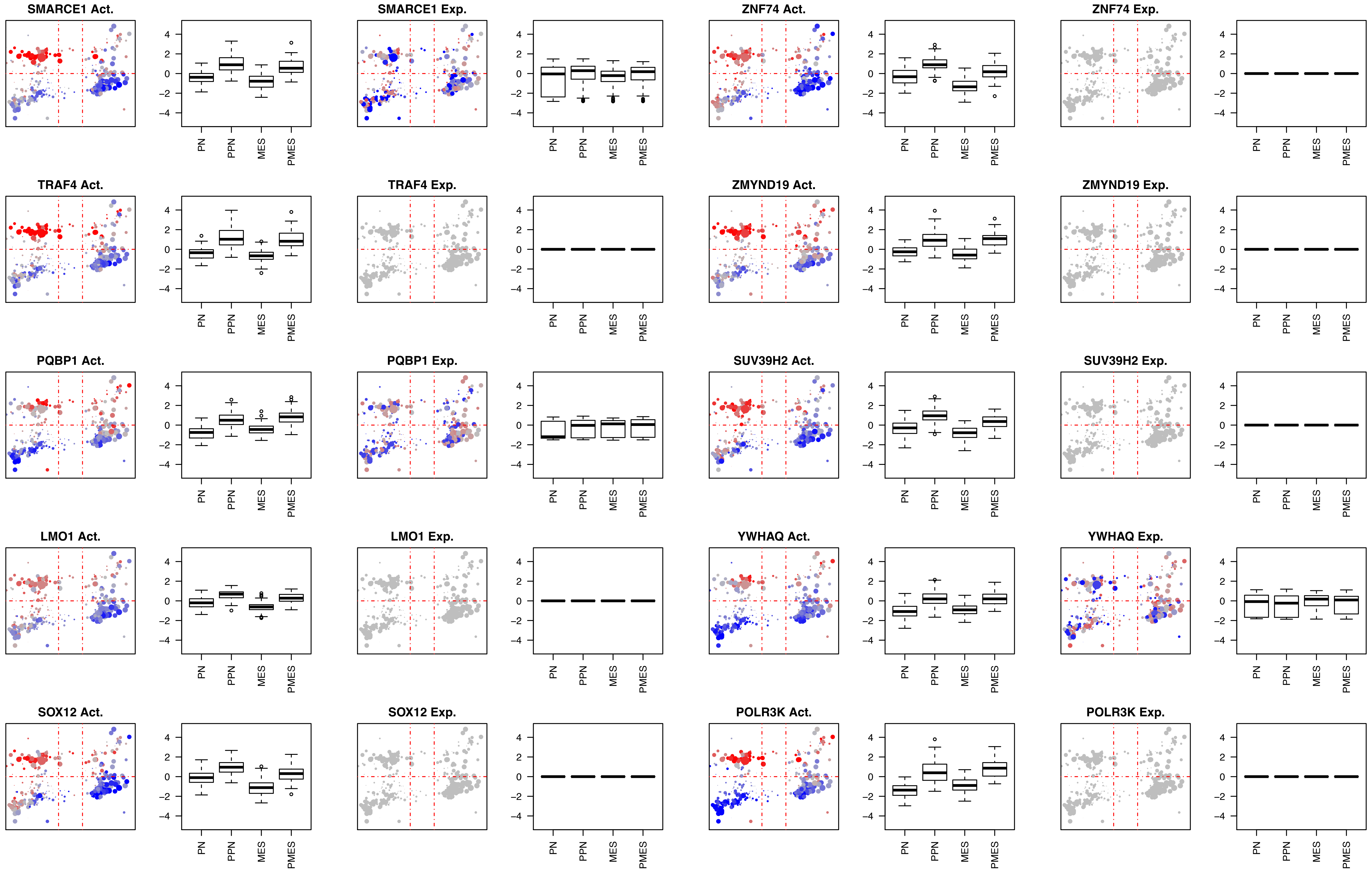
SMARCE1 Act.
SMARCE1 Exp.
ZNF74 Act.
ZNF74 Exp.
TRAF4 Act.
TRAF4 Exp.
ZMYND19 Act.
ZMYND19 Exp.
PQBP1 Act.
PQBP1 Exp.
SUV39H2 Act.
SUV39H2 Exp.
LMO1 Act.
LMO1 Exp.
YWHAQ Act.
YWHAQ Exp.
SOX12 Act.
SOX12 Exp.
POLR3K Act.
POLR3K Exp.
4
2
0
−2
−4
PN
PPN
MES
PMES

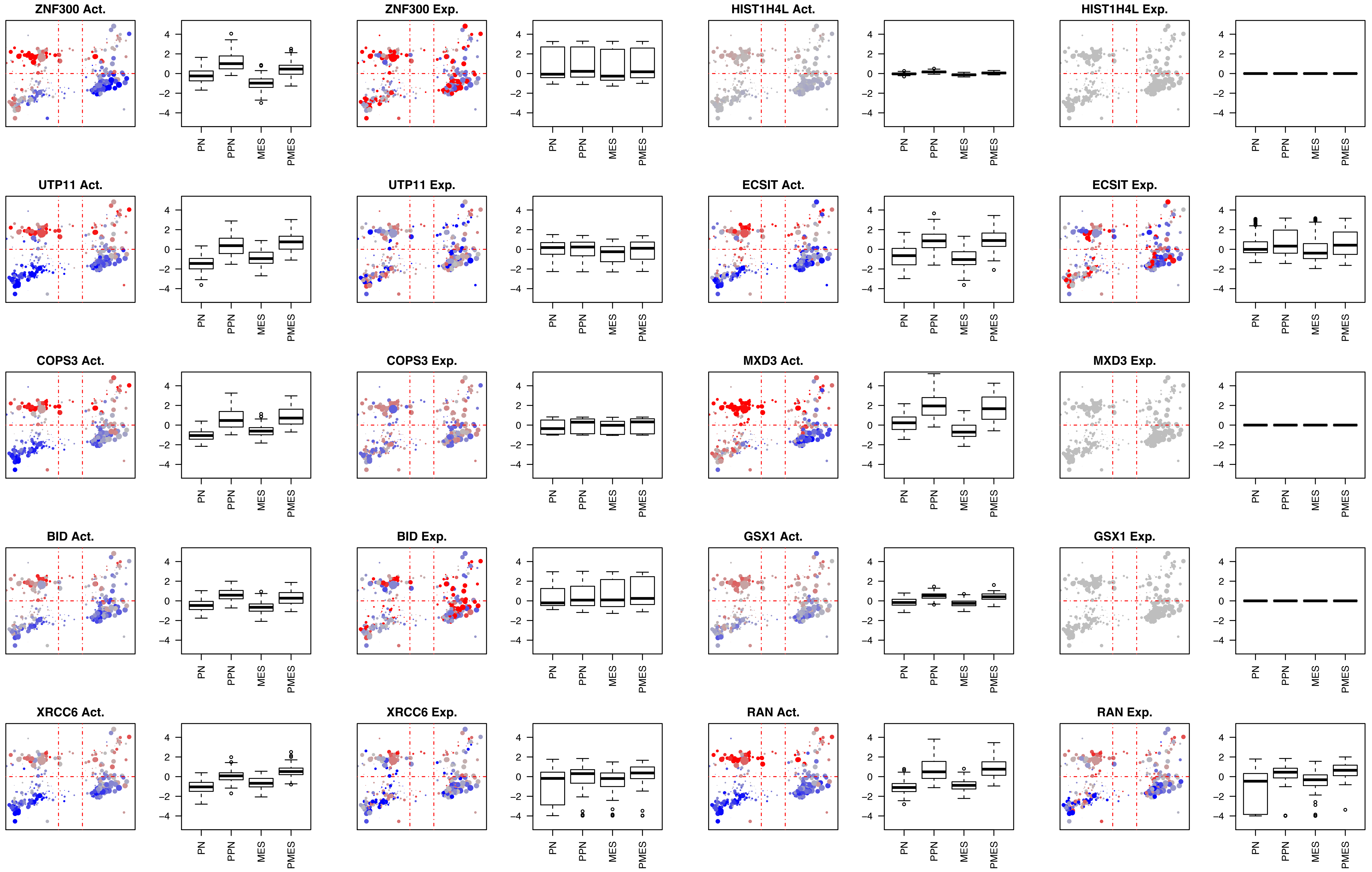
ZNF300 Act.
ZNF300 Exp.
HIST1H4L Act.
HIST1H4L Exp.
UTP11 Act.
UTP11 Exp.
ECSIT Act.
ECSIT Exp.
COPS3 Act.
COPS3 Exp.
MXD3 Act.
MXD3 Exp.
BID Act.
BID Exp.
GSX1 Act.
GSX1 Exp.
XRCC6 Act.
XRCC6 Exp.
RAN Act.
RAN Exp.
4
2
0
−2
−4
PN
PPN
MES
PMES

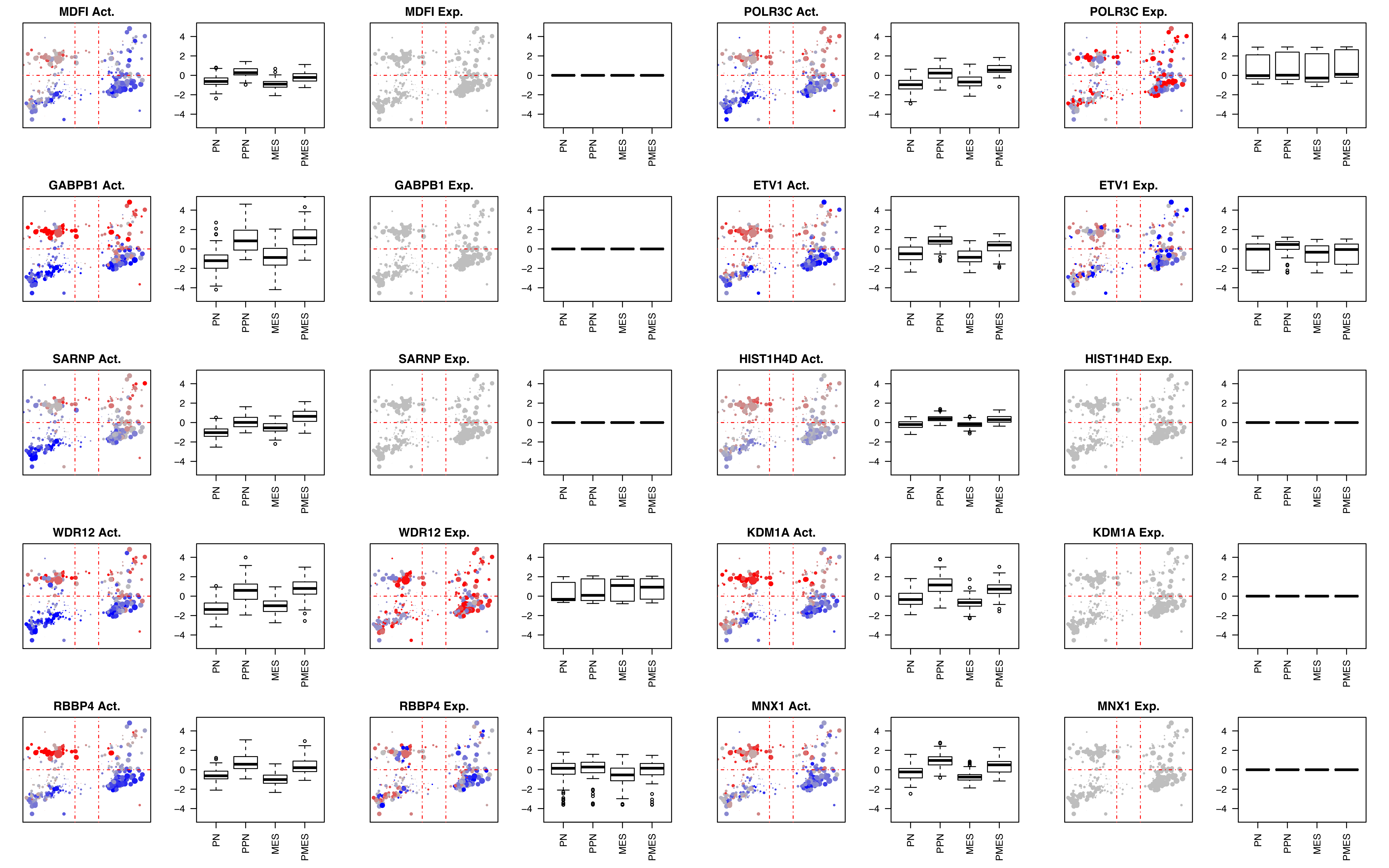
MDFI Act.
MDFI Exp.
POLR3C Act.
POLR3C Exp.
GABPB1 Act.
GABPB1 Exp.
ETV1 Act.
ETV1 Exp.
SARNP Act.
SARNP Exp.
HIST1H4D Act.
HIST1H4D Exp.
WDR12 Act.
WDR12 Exp.
KDM1A Act.
KDM1A Exp.
RBBP4 Act.
RBBP4 Exp.
MNX1 Act.
MNX1 Exp.
4
2
0
−2
−4
PN
PPN
MES
PMES

**Proteins aberrantly activated in the PRO- quadrants.**

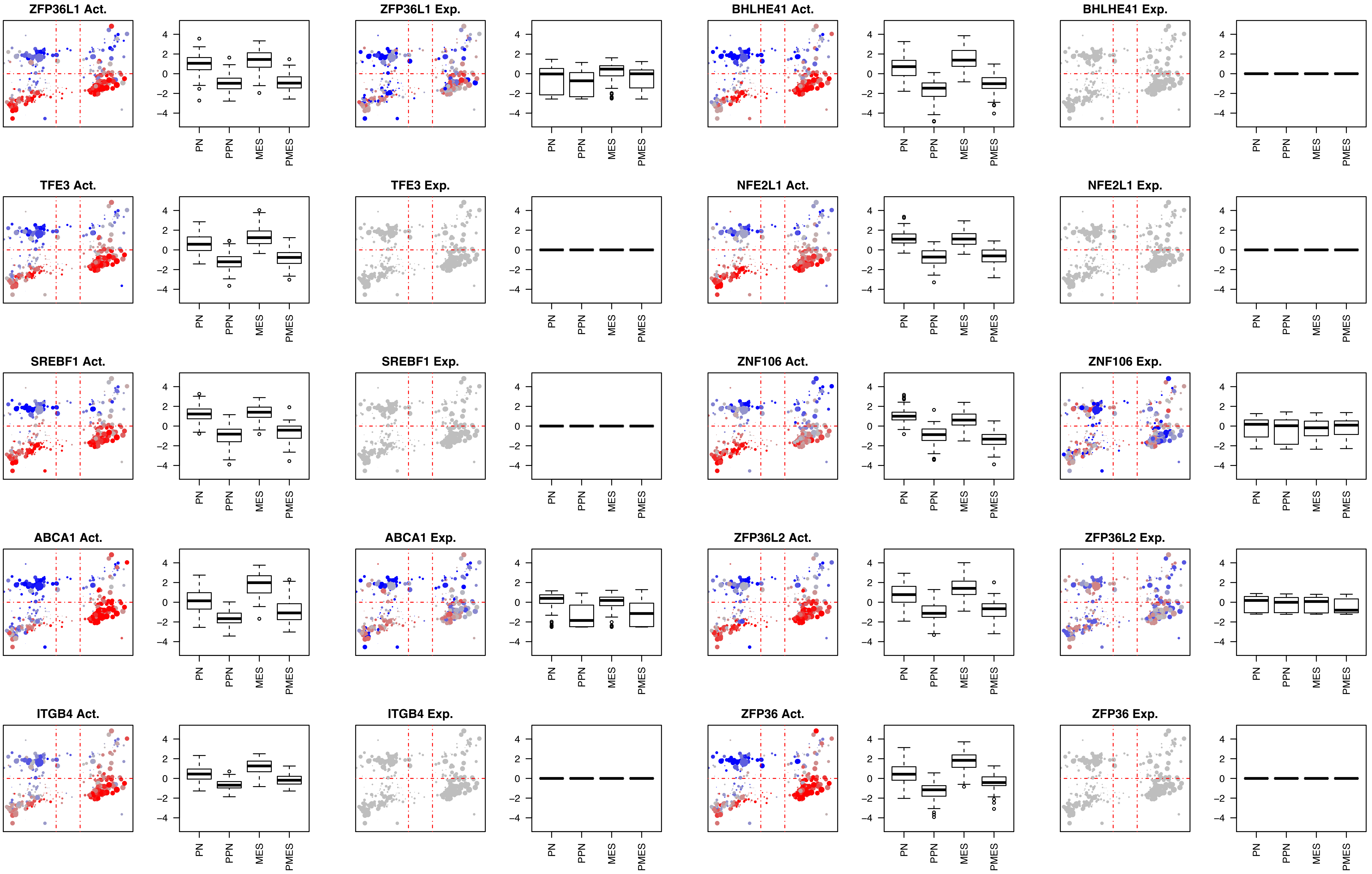
ZFP36L1 Act.
ZFP36L1 Exp.
BHLHE41 Act.
BHLHE41 Exp.
TFE3 Act.
TFE3 Exp.
NFE2L1 Act.
NFE2L1 Exp.
SREBF1 Act.
SREBF1 Exp.
ZNF106 Act.
ZNF106 Exp.
ABCA1 Act.
ABCA1 Exp.
ZFP36L2 Act.
ZFP36L2 Exp.
ITGB4 Act.
ITGB4 Exp.
ZFP36 Act.
ZFP36 Exp.
4
2
0
-2
-4
PN
PPN
MES
PMES

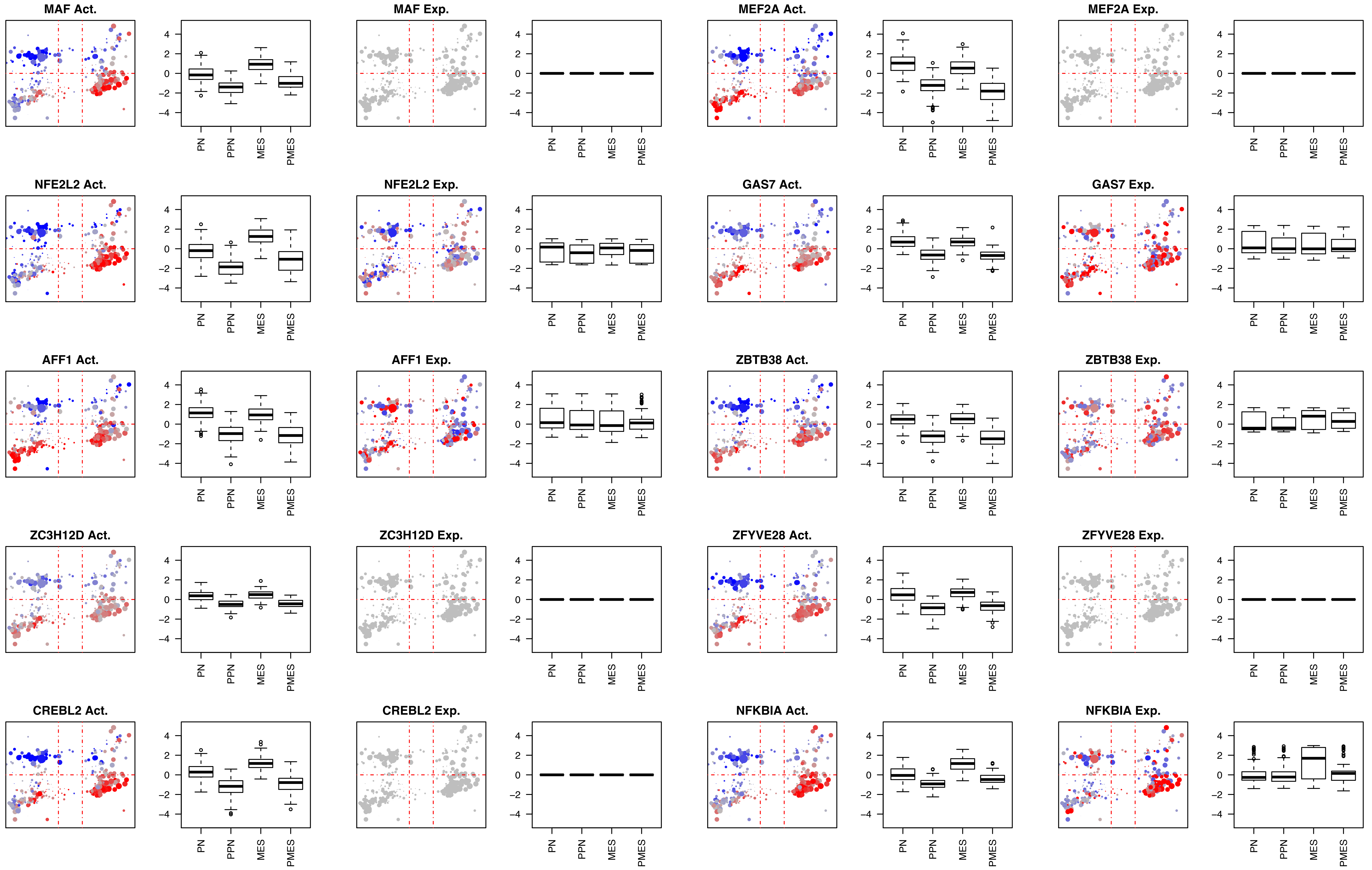

MAF Act.
MAF Exp.
MEF2A Act.
MEF2A Exp.
NFE2L2 Act.
NFE2L2 Exp.
GAS7 Act.
GAS7 Exp.
AFF1 Act.
AFF1 Exp.
ZBTB38 Act.
ZBTB38 Exp.
ZC3H12D Act.
ZC3H12D Exp.
ZFYVE28 Act.
ZFYVE28 Exp.
CREBL2 Act.
CREBL2 Exp.
NFKBIA Act.
NFKBIA Exp.
4
2
0
−2
−4
PN
PPN
MES
PMES

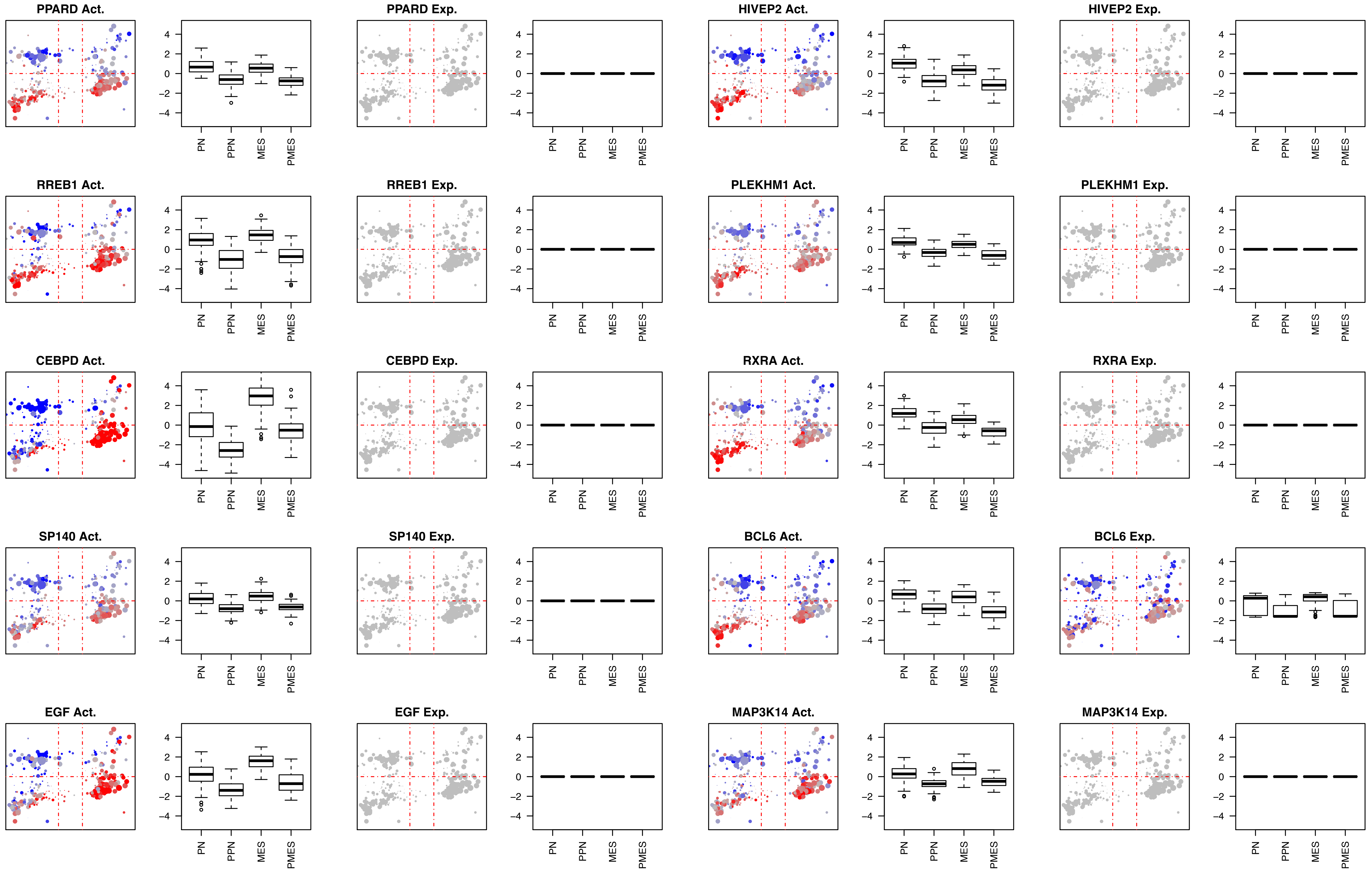
PPARD Act.
PPARD Exp.
HIVEP2 Act.
HIVEP2 Exp.
RREB1 Act.
RREB1 Exp.
PLEKHM1 Act.
PLEKHM1 Exp.
CEBPD Act.
CEBPD Exp.
RXRA Act.
RXRA Exp.
SP140 Act.
SP140 Exp.
BCL6 Act.
BCL6 Exp.
EGF Act.
EGF Exp.
MAP3K14 Act.
MAP3K14 Exp.
4
2
0
−2
−4
PN
PPN
MES
PMES

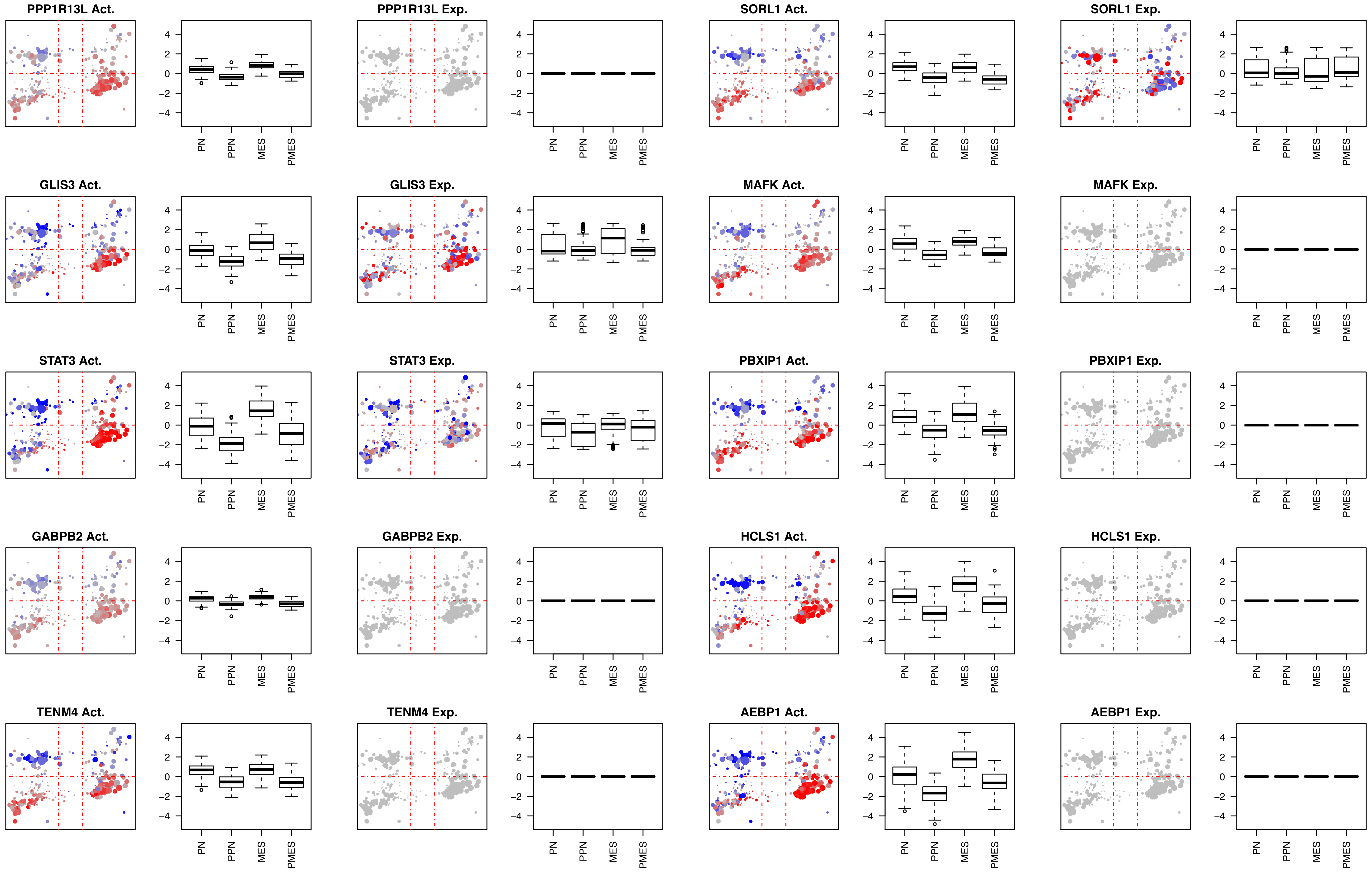
PPP1R13L Act.
PPP1R13L Exp.
SORL1 Act.
SORL1 Exp.
GLIS3 Act.
GLIS3 Exp.
MAFK Act.
MAFK Exp.
STAT3 Act.
STAT3 Exp.
PBXIP1 Act.
PBXIP1 Exp.
GABPB2 Act.
GABPB2 Exp.
HCLS1 Act.
HCLS1 Exp.
TENM4 Act.
TENM4 Exp.
AEBP1 Act.
AEBP1 Exp.
4
2
0
-2
-4
PN
PPN
MES
PMES

### Fig5SupFig-6

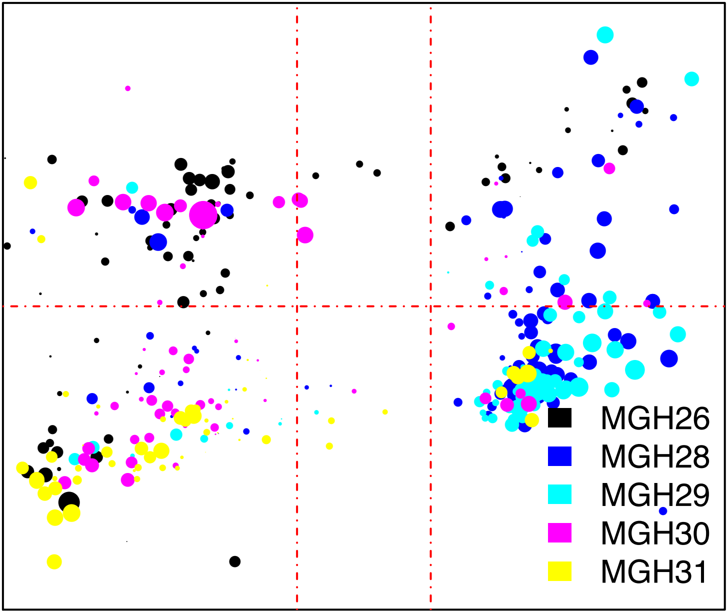

Fig7SupFig-1

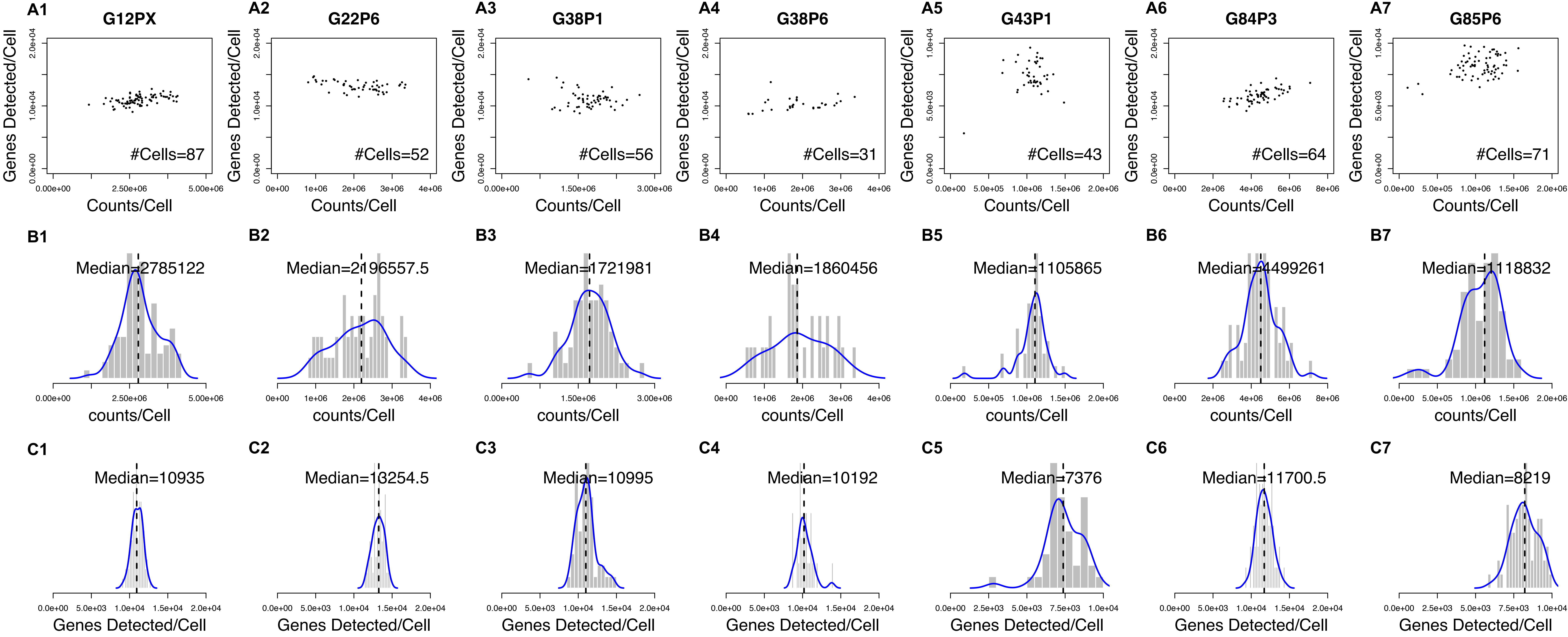

Fig7SupFig-2

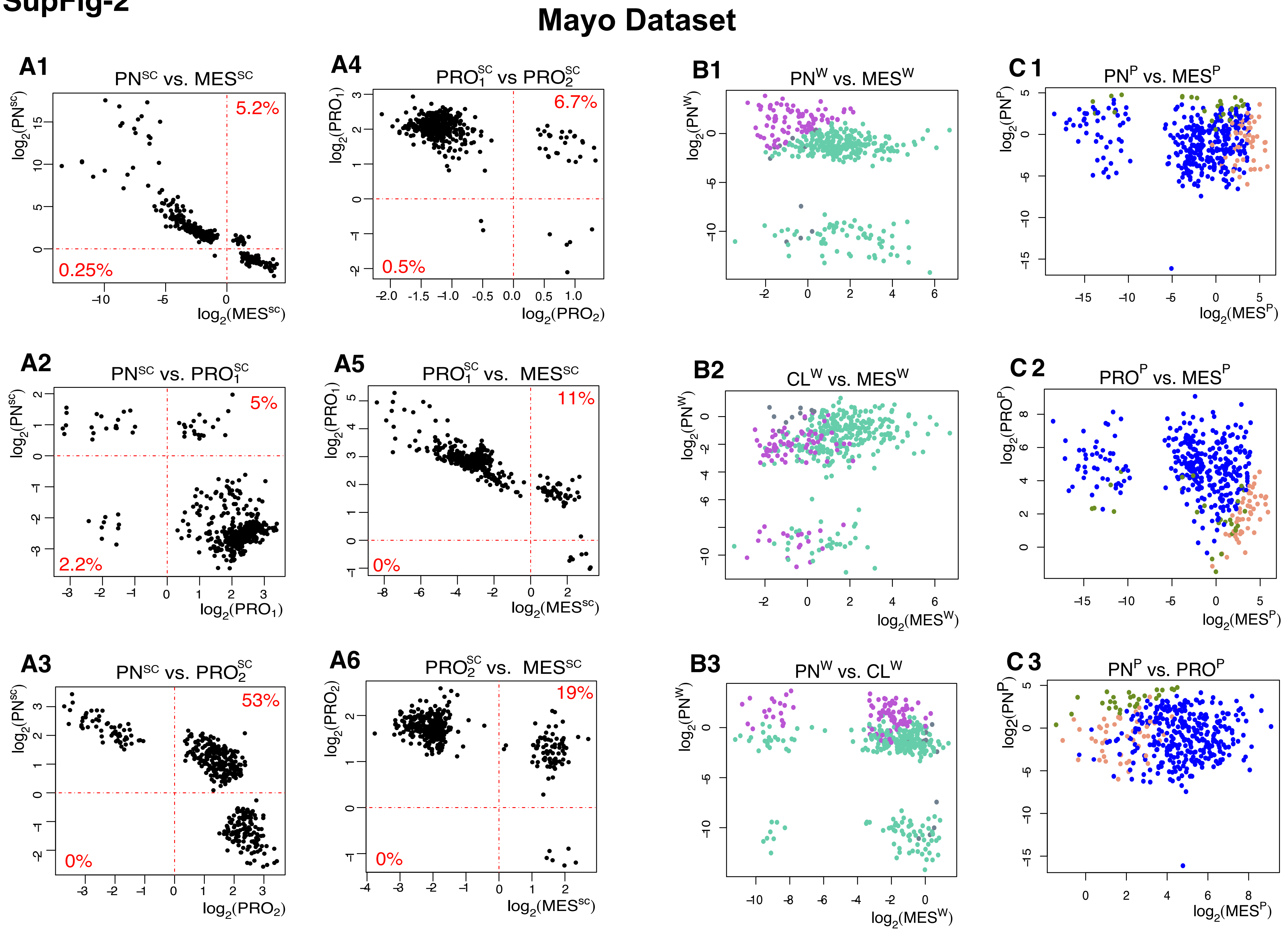

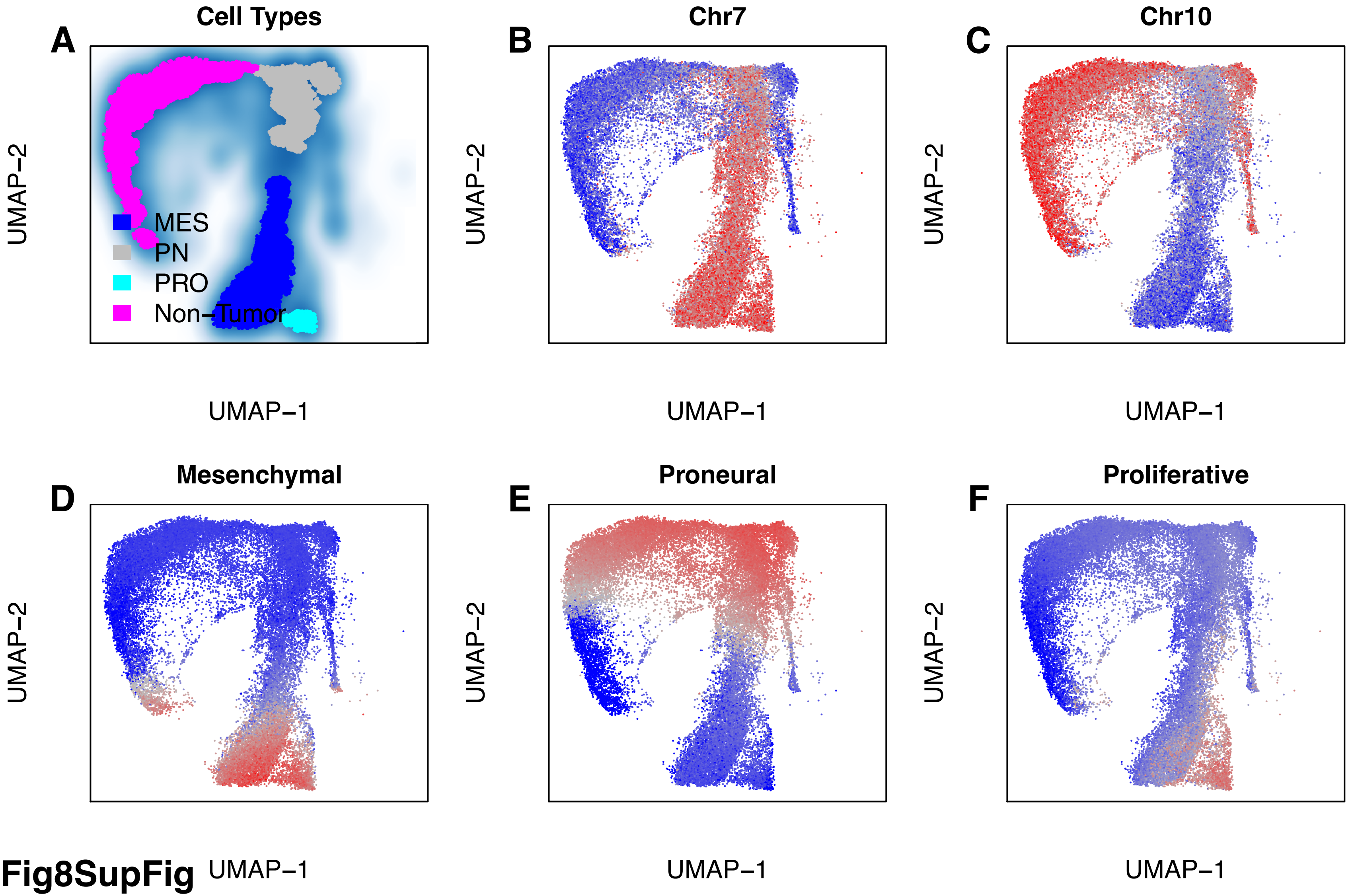
Cell Types
A
MES
PN
PRO
Non−Tumor
UMAP−2
UMAP−1
Chr7
B
UMAP−2
UMAP−1
Chr10
C
UMAP−2
UMAP−1
Mesenchymal
D
UMAP−2
UMAP−1
Proneural
E
UMAP−2
UMAP−1
Proliferative
F
UMAP−2
UMAP−1

Fig8SupFig
