## Supplementary Information for "Single-cell based elucidation of molecularly-distinct glioblastoma states and drug sensitivity"

**Fig1SupFig.** Independent gene-expression-based cluster analysis of multiple datasets—including (a) RNA-Seq profiles from patient-derived samples in (Wang et al., 2017) and (Phillips et al., 2006), (b) RNA-Seq from single cells dissociated from patient resections in (Patel et al., 2014) and (Darmanis et al., 2017), and (c) scRNA-Seq from single cells dissociated from orthotopically implanted PDX models—produced mutually-inconsistent subtypes and low Silhouette-Score clusters. In contrast, metaVIPER-based protein activity analysis (Ding et al., 2018a), using ARACNe-inferred regulatory networks (Basso et al., 2005) from five GBM datasets, identified four novel, highly-conserved subtypes with high Silhouette Scores. Single-cell analyses, using the OncoTreat algorithm (Alvarez et al., 2018), predicted that sensitivity to clinically-relevant drugs is subtype-specific, as experimentally confirmed in patient-derived explants and PDX models.

**Fig2SupFig.** Clustering analysis on Phillips and Wang samples. For single-pass solution, silhouette scores were used to determine optimal number of clusters at both protein activity and gene expression level (A-D). Detailed clustering analysis results, including sample pairwise distances, silhouette scores and subtype annotation in the original studies were visualized in (E-H). At protein activity level, for the optimal 2-cluster solution, ROC curves of cross-cohort random-forest classification analysis were shown in I and J Further, cluster-specific activated proteins in Phillips and Wang datasets were shown in K and J, respectively. The 2-cluster scheme can be further split by iterClust iterative clustering analysis, in both Phillips (M) and Verhaak (N) datasets.

**Fig3SupFig.** Different iterClust schemes (k=2->4, k=3->4) on (Patel et al., 2014) dataset.

**Fig4SupFig-1.** For (Patel et al., 2014) dataset, pairwise GSEA score towards single cell (PN, MES, $PRO_1$ and $PRO_2$) subtypes (see Figure 3) were shown in (A1-A6). Cells were color-coded according to corresponding single cell subtypes. Pairwise GSEA score towards Wang and Phillips subtypes were shown in (B1-B3) and (C1-C3), respectively. Cells were color-coded according to Wang or Phillips annotation.

**Fig4SupFig-2.** For (Darmanis et al., 2017) dataset, pairwise GSEA score towards single cell (PN, MES, $PRO_1$ and $PRO_2$) subtypes (see Figure 3) were shown in (A1-A6). Pairwise GSEA score towards Wang and Phillips subtypes were shown in (B1-B3) and (C1-C3), respectively. Cells were color-coded according to Wang or Phillips annotation.

**Fig4SupFig-3.** Annotating Phillips (A) and Wang (B) bulk samples using single cell classifier (see STAR Methods), further visualizing on the 4-quardrant plots. Samples were color-coded according to clusters determined in (Fig2SupFig MN).

**Fig5SupFig-1.** The activity and expression of master regulators reported in (Verhaak et al., 2010).

**Fig5SupFig-2.** The activity and expression of top 40 proteins aberrantly activated in the PN, PPN, MES and PMES quadrants.

**Fig5SupFig-3.** The activity and expression of master regulators reported in (Carro et al., 2010).

**Fig5SupFig-4.** The activity and expression of top 40 proteins aberrantly activated in the Proneural (PN+PPN) and Mesenchymal (MES+PMES) quadrants.

**Fig5SupFig-5.** The activity and expression of top 40 proteins aberrantly activated in the PRO+ and PRO- quadrants.

**Fig5SupFig-6.** (Patel et al., 2014) single cells were color-coded with corresponding patient IDs on the 4-quardrant plot.

**Fig7SupFig-1.** Quality control for Mayo datasets, including saturation (A1-A7), mapped reads (B1-B7) and transcriptome complexity (C1-C7).

**Fig7SupFig-2.** For combined Mayo datasets, pairwise GSEA score towards single cell (PN, MES, $PRO_1$ and $PRO_2$) subtypes (see Figure 3) were shown in (A1-A6). Pairwise GSEA score towards Wang and Phillips subtypes were shown in (B1-B3) and (C1-C3), respectively. Cells were color-coded according to Wang or Phillips annotation.

**Fig8SupFig.** Clustering analysis identified 4 clusters, which were further annotated as MES, PN, PRO, and non-tumor populations. Copy number of chromosome 7 and 10 was used to separate tumor-related and non-tumor populations (B, C). Tumor-related clusters were further annotated by single cell classifier (see STAR Methods) as MES, PN and

PRO subtypes.

**Fig2SupTable.** Summary of GBM-related regulatory networks used by metaVIPER.

**Fig3SupTable-1.** Summary of iterClust result for Patel single cell dataset.

**Fig3SupTable-2.** Single cell classifier: subtype-specific activated TFs.

**Fig3SupTable-3** Comparing Carro MRs to single cell classifier.

**Fig7SupTable** Summary of processed GBM samples provided by Mayo Clinic.
